# A Dirichlet-multinomial mixed model for determining differential abundance of mutational signatures

**DOI:** 10.1101/2024.03.07.583837

**Authors:** Lena Morrill Gavarró, Dominique-Laurent Couturier, Florian Markowetz

## Abstract

Mutational processes of diverse origin leave their imprints in the genome during tumour evolution. These imprints are called *mutational signatures* and they have been characterised for point mutations, structural variants and copy number changes. Each signature has an *exposure*, or abundance, per sample, which indicates how much a process has contributed to the overall genomic change. Mutational processes are not static, and a better understanding of their dynamics is key to characterise tumour evolution and identify cancer weaknesses that can be exploited during treatment. However, the structure of the data typically collected in this context makes it difficult to test whether signature exposures differ between samples or time-points. In general, the data consist of (1) patient-dependent vectors of counts for each sample and clonality group (2) generated from a covariate-dependent and compositional vector of probabilities with (3) a possibly group-dependent over-dispersion level. To model these data, we build on the Dirichlet-multinomial model to be able to model multivariate overdispersed vectors of counts as well as within-sample dependence and positive correlations between signatures. To estimate the model parameters, we implement a maximum likelihood estimator with a Laplace approximation of the random effect high-dimensional integrals and assess its bias and coverage by means of Monte Carlo simulations. We apply our approach to characterise differences of mutational processes between clonal and subclonal mutations across 23 cancer types of the PCAWG cohort. We find ubiquitous differential abundance of clonal and subclonal signatures across cancer types, and higher dispersion of signatures in the subclonal group, indicating higher variability between patients at subclonal level, possibly due to the presence of different clones with distinct active mutational processes. Mutational signature analysis is an expanding field and we envision our framework to be used widely to detect global changes in mutational process activity.

**Author Summary:** The genome is permanently subject to alterations due to errors in replication, faulty replication machinery, and external mutational processes such as tobacco smoke or UV light. Cancer is a disease of the genome, characterised by an abnormal growth of cells that harbour the same set of “clonal” mutations. In turn, these mutations might transform how cells accrue new “subclonal” mutations or the extent to which they tolerate them. The mutational signature framework lets us extract the information of which mutational processes have been active, and in which intensity, in creating a set of mutations. We extend this framework to statistically test the change in the relative intensity of mutational processes between conditions. In samples of 23 cancer types of the PCAWG project, we test the difference between mutational processes that contribute to mutations prior to cancer onset (clonal group), and upon cancer onset (subclonal group), whilst keeping into consideration patient-to-patient differences. We find differences in the majority of cancer types, and identify mutational processes which contribute preferentially to either group.

## 1 Introduction

Cancer is a disease of the genome, which is permanently subject to alterations due to external factors, to inevitable errors in replication, and to faulty replication machinery [1, 2, 3, 4, 5]. Cancer cell populations grow by clonal expansions. A typical tumour carries several clones – populations of cells with a common genotype. A mutation shared by all cancer cells in the genome is called a clonal mutation, in opposition to a subclonal mutation, which appears in subsequent clones. Fig 1 shows a diagram of clonal evolution in a cancer sample.

**Figure 1.**
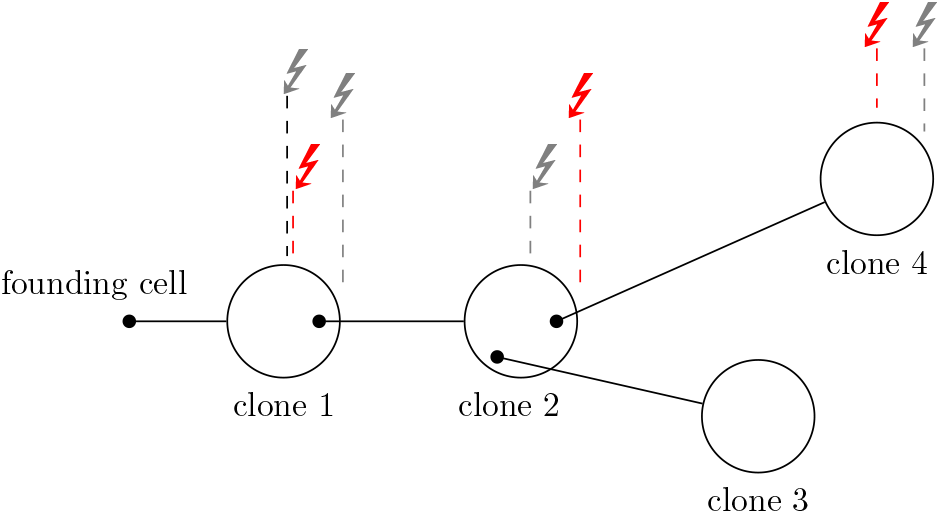
Example of a clonal tree. Four clones are drawn by big circles. A smaller black circle indicates the founder cell of the subsequent clone. The left-most dark circle is the MRCA cell of the tumour, which undergoes clonal expansion to create the first clone, which is defined by clonal mutations of a cancer cell fraction (CCF= 1). Subsequent clones are defined by subclonal mutations of CCF*<* 1. The mutations that define each clone might be of diverse origin: here we depict two active mutational processes (grey and red arrows) creating mutations at two distinct rates. Whilst the mutational process represented by grey arrows is more active at early stages of tumour development, in creating the mutations that define the first clone, the mutational process presented by red arrows has a constant mutation rate.

### Mutational signatures

Computational methods on large-scale, genome-wide, genomic data (commonly whole-genome sequencing, WGS) have enabled the use of *mutational signatures* as a proxy for mutational processes to measure the number of processes active in a tumour and quantify the number of mutations created by each one. Mutational signatures represent the processes of mutation and of any subsequent repair [6, 2, 7, 8]. They are particularly well established in the context of point mutation signatures, which are repeatedly re-defined and curated in the COSMIC database [9]. In COSMIC signatures, six types of mutations are considered (`C`*→*`A`, `C`*→*`G`, `C`*→*`T`, `T`*→*`A`, `T`*→*`C`, and `T`*→*`G`), as well as their immediate context (their 5’ and 3’ flanking bases), but generally without including information about their strandness. This leads to a classification into 96 (= 6 *·* 4 *·* 4) categories. Therefore, a COSMIC signature is a vector of probabilities of length 96 which sums up to one and indicates the preferences in creating each type of mutation. In the latest version of these signatures (v3.4, June 2023) there are 86 such signatures: SBS1 to SBS99, with some intermediate labels referring to deprecated signatures. As explained in more detail below, in a sample, each signature has a corresponding *mutational signature exposure*, indicating the number of mutations attributable to the signature in the sample. These are the key quantities we would like to model. Mutational signatures and their exposures are extracted assuming that the total number of mutations are a linear combination of signature contributions – the exposures being the coefficients in this linear combination. Section S1 provides an overview of signature extraction methods as one of their outputs is the exposure matrix **Y** corresponding to the response in the models we develop. Table 1 provides a list of notation used.

**Table 1:**
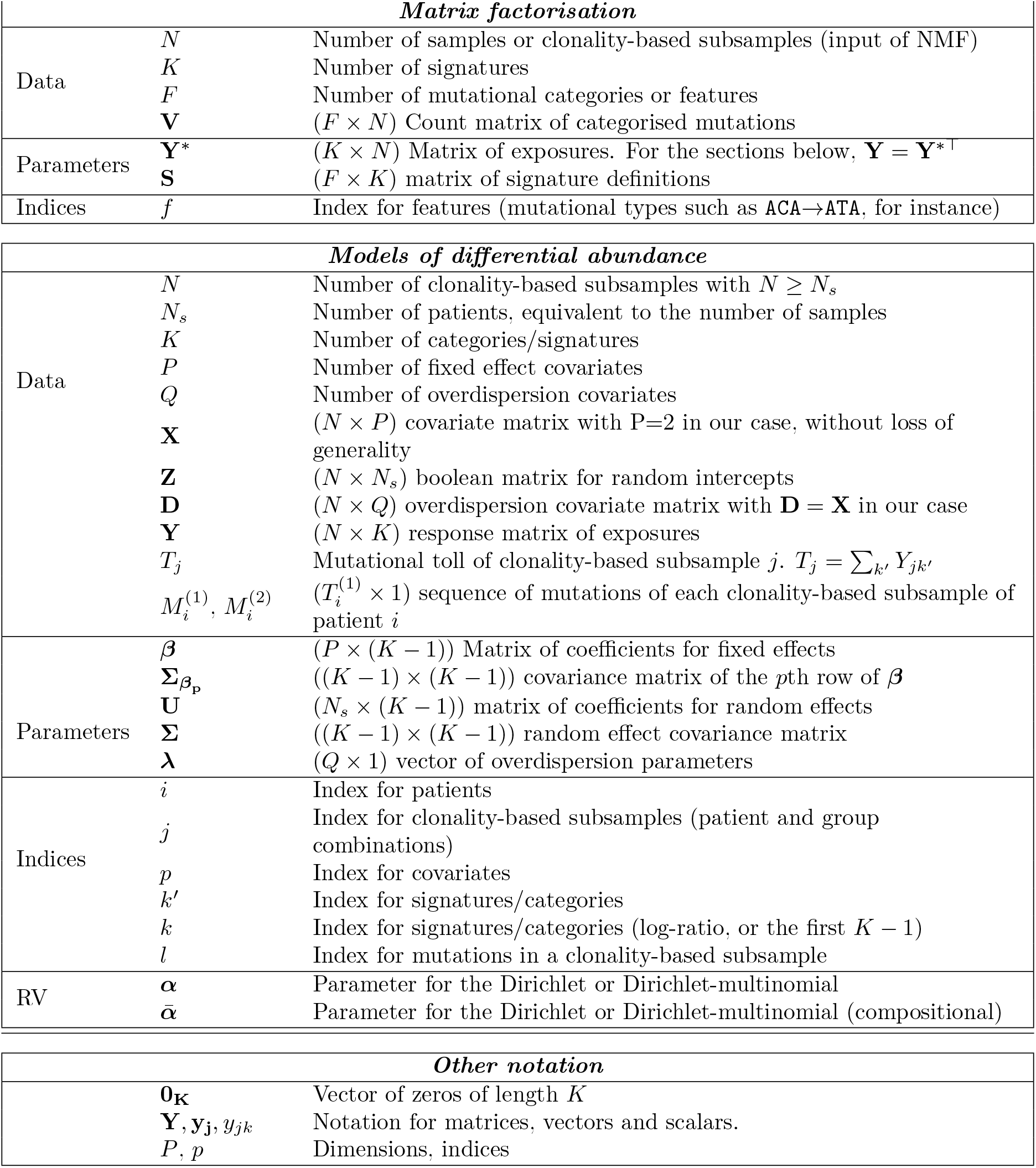
List of notation.

### A short history of signature analysis

Established between 2012 and 2013, the field of mutational signatures promised not only to bridge the gap between the observable, and possibly very complex, mutational landscape of a tumour at the time of sequencing and its aetiology, but also to be clinically relevant. In the first instance, patient stratification [10] based on exposures suggested treatment groups. This approach has delivered in the case of homologous recombination deficiency (HRD), where mutational signatures are used to determine suitability of PARPi treatment [11]. The use of signatures has, moreover, enabled to discern the nature of other mutational processes, notably APOBEC [12, 13]. However, the number of mutational signatures has been ever-increasing – at least partially due to the increased number of samples used for signature derivation – but few have an aetiology unequivocally linked to them, hampering the interpretation of results and even making the analyses difficult from a statistical point of view. Finally, several variations on single-base substitution signatures have been proposed and adopted: doublet-base substitution signatures [14], indel signatures [14] and copy number signatures [15, 16]. [17] list several statistical challenges in the field, including the difficult choice between alternative methods of signature extraction (each with different assumptions) and the inclusion of uncertainty quantification around exposures, which warrants new statistical methods. Even with simpler signature extraction methods the analysis of these data – such as differential abundance analyses – can be complex, and this is the focus of this paper.

### Motivating example

We wish to detect differential abundance between clonal and subclonal mutations in the PCAWG cohort, in a histopathological cancer type-specific analysis of 23 cancer types of 27 samples or more. Signatures exposures are extracted using quadratic programming as shown in [18]. The comparison to values obtained by considering `mutSigExtractor` [19], as well as to values reported in [20] and extracted using `sigProfiler` and without splitting the mutations into two groups, yield comparable results (Fig S1). Signatures are extracted using the subset of COSMIC signatures that were considered to be active in each cancer type, taken from [14]. Mutations are classified as clonal and subclonal following [21], considering as clonal mutations which appear in all cells of the tumour, and as subclonal mutations only appearing in a fraction of cells. Details on the assignment of mutations to these categories are beyond the scope of this paper, as this is a step that requires inference of the clonal population structure of the model, done in a per-sample basis and requiring careful consideration – in [20] 11 clonal structure identification methods are used, from which a consensus is derived.

### Signature exposures as compositional data

In the case under consideration – clonal and subclonal mutations – as well as in most comparisons of mutational signature exposures, the total number of mutations in each clonality-based subsample is not of interest; instead, the crucial information is the relative allocation of these signatures into categories (mutational signatures). This renders their analysis suitable in a compositional setting. [22] noted that the data can be encoded, without loss of information, using log-ratios. Compositional data in the form of probabilities can be transformed to log-ratios and back using the additive log-ratio transformation (ALR) and inverse additive log-ratio transformation (ALR^*−*1^) respectively (see Section S2 for an introduction to compositional data). In short, compositional data are relative data which must be analysed in relative terms, an important consideration that has usually been overlooked in the mutational signature literature (with few exceptions: [23, 24], only the latter mentioning compositionality), as well as in several other quantitative branches of the sciences where compositional data are frequent, and which can lead to spurious correlations and negative biases.

### Previous work

Two papers to date – HiLDA [23] and TCSM [24] – have shown statistical analyses specific to the comparison of exposures in two groups. TCSM tests for differential abundance between two groups of samples following signature extraction and by simulation through the logistic-normal distribution. In HiLDA, the Dirichlet-multinomial is used to model signatures in an MCMC based inference framework. Neither are suitable for a post-hoc analyses of exposures (e.g. using COSMIC signatures) nor do they include random effects. As compositional data arise in multiple situations, models that analyse equivalent data in other disciplines are more widespread. Recently, [25] implemented a Dirichlet-multinomial model with mixed effects, of which the model described here can be seen as an intension. Table S1 proposes a list of existing methods for either studying the dynamics of mutational signatures or, more generally, detecting differential abundance in compositional data.

## 2 Methods

In this paper, we propose an estimator for the Dirichlet-multinomial mixed effect model with a multivariate and unconstrained structure for the random effects as well as a group-specific overdispersion parameter to detect differences in mutational signatures between groups of observations. We also opted for the Laplace analytical approximation (LA) to evaluate the high dimensional integrals induced by the random effect structure. Our results might benefit from the good properties of LA in regards to bias and mean-square errors [26] as well as convergence rate and coverage [27]. Speed is also an attractive feature of the Laplace approximation compared to alternatives [28], as well as the fact that no parameters for integration, such as the number of quadrature points, have to be specified.

The remaining of this paper is organised as follows: the Methods section presents our model and its implementation with Laplace approximation, describes the operating characteristics of our estimator by Monte Carlo simulations, and compares the results to those of existing methods. The Results section is dedicated to the application of our model to the PCAWG dataset, first by using the simple six point-mutation categorisation, and then by using mutational signatures as categories. Finally, we present our conclusions and a discussion on the biological implications of the results and future directions, and we provide some guidance on each step of the pipeline of the R package `CompSign`, where our models are made available.

### 2.1 Mixed effects Dirichlet-multinomial regression

#### Data

Our goal is to compare signature exposures between two sets of samples. For simplicity we assume that we have paired observations for *N*_*s*_ patients and thus *N* = 2*N*_*s*_ observations in total, although our method is general and can also be applied to un-paired cohorts of different sizes, or to cohorts containing multiple samples per patient. As for each patient a single biological sample is taken, in this document we use the word “sample” to denote the biological sample, and it is therefore used interchangeably with “patient”. In turn, each sample is split into two “clonality-based subsamples”, which populate the first (clonal) and second (subclonal) groups of subsamples. These clonality-based subsamples correspond to the response in the model. Let *K* denote the number of signatures or categories, *i* index patients (or any other group defined by the mixed effects), *k* index either the first *K −* 1 signatures or *K −* 1 signature log-ratios, and *j* index clonality-based subsamples. Let *P* be the number of covariates. Comparing two groups and including a shared intercept leads to *P* = 2.

Our data are as follows. We have two exposure matrices of counts corresponding to the clonal and subclonal (clonality-based) groups. Each element in each of the two matrices corresponds to the same patient and the same category or signature. Let’s denote these matrices by 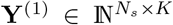 and 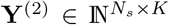 where (1) and (2) refer to clonal and subclonal groups, which might have been independently derived. In the case under consideration, for each sample and clonality-based group (i.e., for each clonality-based subsample), mutations are classified according to their trinucleotide context, and signature exposures are estimated independently. We row-concatenate these matrices to create **Y** *∈* ℕ^*N×K*^ where *N* = 2*N*_*s*_.

#### Fixed-effects

Firstly, we express the log-ratios of signature abundances as a function of the fixed effects. The fixed effects are captured by the **X** ∈ ℝ^*N×P*^ design matrix, and their corresponding coefficients are a ***β*** ∈ ℝ^*P ×*(*K−*1)^ parameter matrix. In the applications shown here **X** is a binary matrix of ones in the first row, and zeros or ones in the second (with a zero if the sample belongs to the clonal group, and a one otherwise). In practice, the implementation we propose does not have this constraint: **X** can take any value in ℝ and *P* can be greater than 2.

The first row of ***β, β***_0_, is a vector of length *K −* 1 which corresponds to the general abundance of signatures in the reference mutational group (clonal group), expressed as the ALR transformation using the last signature as reference, i.e. as the log-ratio of a signature to a baseline signature. Let 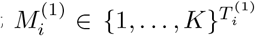 be the sequence of mutations for patient *i* in the clonal group, and 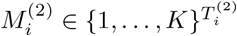 the equivalent for the subclonal group. Let these mutations be indexed by *l*. Thus, for *k < K* and a case without patient-dependence (i.e., no random effect), we have

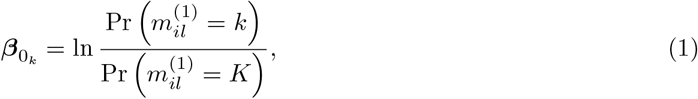

where 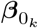 denotes the *k*th element of ***β***, and 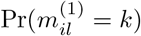 denotes the probability of the *l*th mutation of patient *i* in the reference group to have been generated by signature *k*. For *k < K*, this probability is then given by

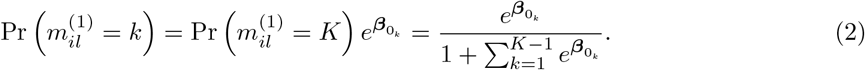

The second row of ***β, β***_1_, is our main parameter of interest, as it indicates whether there are shifts in the general abundance of signatures for samples of the subclonal group compared to the clonal reference in the ALR space. Similarly to above, the log-ratio of the probability of the *l*th count of patient *i* in the second group to have been generated by signature *k*, over the probability of having been generated by signature *K*, is

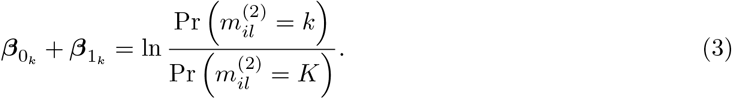

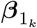 equal to zero indicates that the *k*th log-ratio is the same in both groups, indicating that the relative mutation rates of the signatures of both groups are equal. A zero vector of ***β***_1_ corresponds to cases without differential abundance between groups. A test for overall differential abundance can be computed by using the generalised Wald statistic *w* in a *χ*^2^ test with *K −* 1 degrees of freedom, combining all the estimates of ***β***_1_ and their correlations:

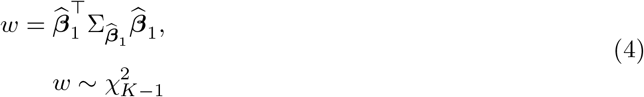

where 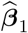 and 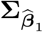 respectively denote the estimated ***β***_1_ parameter vector and its corresponding covariance matrix.

#### Random intercepts

Secondly, the log-ratios of abundances of signatures also depend on the random effects, in which we include information about the patient from whom each observation derives. In the PCAWG dataset, having one observation per patient and group, i.e. two observations per patient, we use the matrix 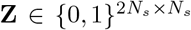 (more generally, 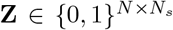) to indicate the patient to whom each observation belongs. The coefficients for **Z** are encapsulated in the matrix 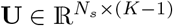, where each row of **U** corresponds to the patient-specific multivariate intercepts and where the values are, too, ALR-transformed. These intercepts are drawn from a multivariate normal distribution of mean zero (as the overall abundance of signatures in the first group is already captured by *β*_0_), and a covariance matrix which can be unconstrained or not. These multivariate random intercepts differentiate our implementation from the one of [25] (in which the computational Gauss-Hermite quadrature is used to approximate the random effect integrals) and allow us not only to model the potential within-patient dependence with more flexibility but also to have positive correlations between signatures.

#### Logit link

The linear combination of fixed and random effects is linked to the log-ratios of abundances as follows, in the commonly used logit link:

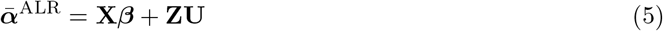

where 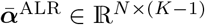 are row-wise ALR-transformed quantities. Therefore, each row can be transformed into a vector of probabilities 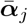 using the inverse-ALR transformation ALR^*−*1^, or generalised softmax transformation:

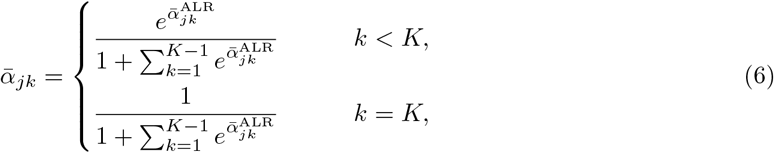

where 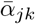 represents the probability of counts of the *j*th clonality-based subsample to have been generated from the *k*th signature. Moreover, we introduce the precision parameter vector ***λ***, such that, for the *j*th subsample, the Dirichlet-multinomial (DM) parameter ***α***_*j*_ is defined as product of the probability described above and the scalar 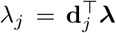, where **d**_*j*_ denotes the *j*th row of the (*N × Q*) **D** overdispersion predictor matrix, so that

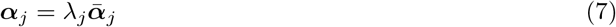

Without loss of generality, we model the overdispresion as a function of the groups only, so that **D** = **X** and the two values of the ***λ*** vector respectively correspond to the overdispersion level in the reference group and to the shift in overdispersion between both groups. Higher values of ***λ*** lead to more concentrated results, or lower overdispersion. In practice, we estimate log(***λ***), to ensure positivity. Lastly, the counts of the *j*th clonality-based subsample are drawn from a Dirichlet-multinomial distribution as follows

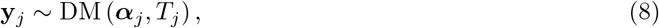

where 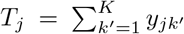 corresponds to the total number of mutations observed for clonality-based subsample *j*.

#### Model summary

Therefore, our mixed-effect Dirichlet-multinomial model can be summarised by the following equations:

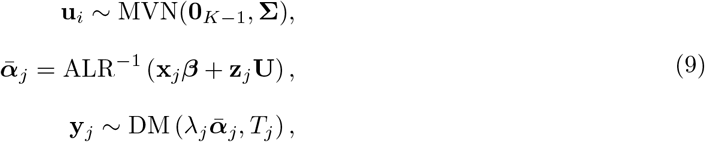

where **Σ** denotes the random effect correlation matrix.

The dependencies between parameters and elements of the data are displayed in the plate diagram proposed in Fig 2.

**Figure 2.**
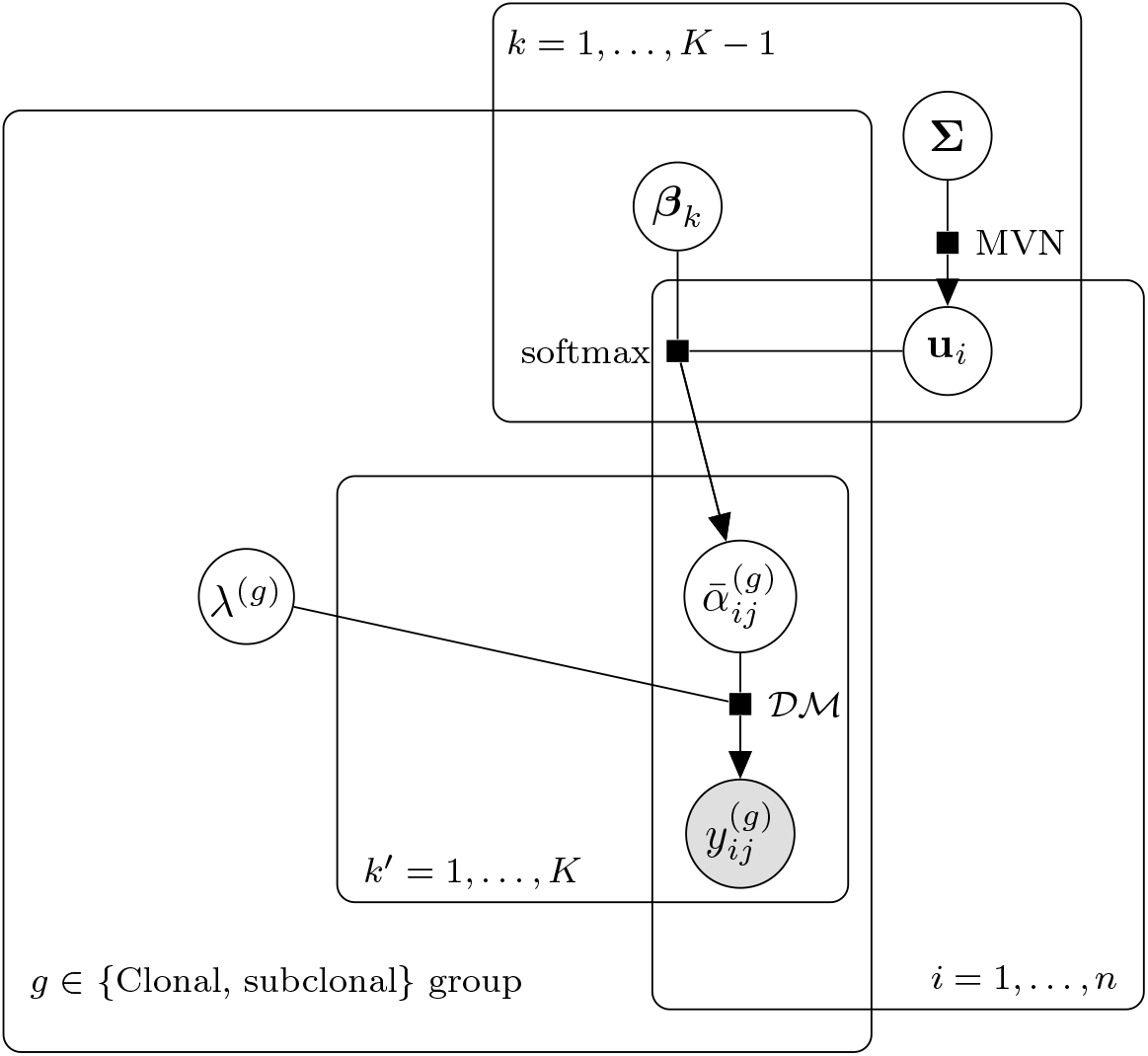
Plate diagram for the Dirichlet-multinomial model. The random intercepts **u**_*i*_ are independent of clonality-based grouping, and in log-ratio space, as are the coefficients ***β***. The overdispersion parameters ***λ*** are a function of the clonality-based grouping exclusively in this manuscript, but adjustable in their implementation. Similarly, **X** and **D** (the covariate matrices for the fixed effects and overdispersion) are shared here and reflect the clonality group *g*, but that need not be the case and they are specified independently in the model implementation.

### 2.2 Variants of the model and implementation in Template Model Builder

In the remaining part of this paper, we compare several variations of the model described above:

- `diagREDM` is a DM model that considers signature-dependent random-effects (independently for each signature log-ratio) and a clonality group-dependent overdispersion parameter *λ*,
- `singleREDM` is a DM model that considers signature-independent random-effects (a univariate random intercept drawn from one single *σ*) and a clonality group-dependent overdispersion parameter *λ*,
- `fullREDM` is a DM model that considers signature-dependent (and possibly correlated) random-effects and a clonality group-dependent overdispersion parameter *λ*.

All models considered in this paper are implemented using Template Model Builder (TMB) [29], a framework for maximum likelihood estimation in random effects models, in which the random effects are integrated out of the likelihood using the Laplace approximation. TMB has an R interface, and the model is written in `C++`. The mixed effects models presented in this paper have been assembled into the R package `CompSign` (https://github.com/lm687/CompSign), which includes vignettes with example data and instructions on how to run the functions and interpret the results. The code for reproducing the results of this paper can be found in https://github.com/lm687/CompSign-results and all datasets and inference results in https://zenodo.org/records/10546525.

### 2.3 Model assumptions

We assume in all cases that the mutational processes active in the clonal group are also clonal in the second group, possibly in different relative activity. As with most compositional models, scenarios where all exposures are zero in a group and non-zero in the other are problematic (Section S2.1), but this is not a scenario seen in any of PCAWG datasets to which the models have been applied. The exposures in the subclonal group might be representative of only a subset of the clones: as *β*_1_ represents a cohort-wide average, a large increase in the mutation rate of a process active in a single clone can result in a moderate *β*_1_ provided the single clone represents a small fraction of cells in the tumour. `diagREDM` assumes that mutational processes do not co-occur across patients, although the multivariate random effects paired with the clonality group-dependent overdispersion parameter ***λ*** allows for correlations to some extent. `fullREDM`, in that it does allow for such correlations or co-occurrences by estimating the covariance matrix of signature abundances, may suffer from convergence problems if some signature is present in very low abundance and the dataset includes few samples.

### 2.4 Simulation

#### 2.4.1 Bias and coverage of estimator

The bias and coverage of 95% confidence intervals for elements of ***β***_1_, the parameter vector of interest, as well as for elements of ***β***_0_, are assessed by means of Monte Carlo simulations considering 1000 simulated datasets of n=200 samples – a sample size in line with the number of patients often considered in such studies. These simulated datasets (A1-3, B1-4) represent cancer type signature exposures at clonality-based subsample level, with the same set of active signatures (although abundant to varying extents). B1-4 have been simulated to match biologically-relevant parameters (Section S3.1). Although the recovery of parameters ***β***_0_ and ***β***_1_ can suffer with increasingly lower values of *λ* (higher dispersion) (Figs S2, S3), both bias and coverage results are satisfactory in all biologically-inspired datasets (Figs S5, S6, S7, S8). When datasets have been created with independence between the random effects, `diagREDM` and `fullREDM` lead to equivalent results (Figs S5, S6). When correlated random effects are used, this can lead to biases in the estimation of elements of ***β***_0_ for `diagREDM` (but not for `fullREDM`), but this does not translate to biases or lower coverages in the elements of ***β***_1_, our parameter of interest, provided multivariate random effects are used (Fig S4, S6). In these simulations the number of mutations simulated in each clonality-based subsample are either representative of the cancer type from which representative parameters were chosen, or much lower than typical number of observed mutations (Fig S9). Indeed, we find that the results of the models are robust to lowering the number of mutations up to 10-fold (Fig S8). Due to the strong reduction in the number of parameters to estimate and their agreement in 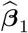, `diagREDM` thus appears as very attractive compared to `fullREDM`.

#### 2.4.2 Comparison with previous models of differential abundance

We compare the output of HiLDA and TCSM to that of `diagREDM`. To avoid simulating from the model, and because both TCSM and HiLDA take as input substitution categories (and not pre-computed exposures), we use a different approach in simulations C1-C3 (Fig S10). In a simulation inspired by [30], biologically-informed exposures are generated by mixing, in some proportion *π*, exposures from two groups of observations, and from those generating 96 substitution category data as input for HiLDA and TCSM. For `diagREDM`, exposures are re-computed from the substitution categories and those become the input for the model. The two groups of exposures for creating the mixture differ between simulation strategies (Section S3.3.1).

The results are markedly different, across simulation strategies and as parameters vary. Whereas HiLDA does not find any differential abundance, TCSM always does, and `diagREDM` is the only model with progressively higher true positive rates as the mixing proportion *π* increases (Table S2). `diagREDM` is consistently the fastest method (Fig S11, Table S3), often by several orders of magnitude. HiLDA, insofar as a Bayesian model, is the slowest, and we were unable to run it in datasets with a high number of mutations (Section S3.3.2). The coefficients that indicate differential abundance in each model differ, complicating the comparison of methods (see Section S3.3.3) but in all three cases a trend of change in these coficients in seen as *π* increases (Fig 3 for C1, and Fig S12, S13 for C2 and C3). For C1, the changes are apparent in `diagREDM` and TCSM even when datasets are mixed in a 8-to-92% proportion, whereas in HiLDA a 50-50% mixture is necessary. Whereas for `diagREDM` an increase in the number of mutations or the number of samples leads to higher statistical power (Fig S14), in TCSM it leads to a better recovery of signature definitions (Fig S15). In C1, the expected relative sparsity of signature exposures in datasets of high *π* is clearest in the results from `diagREDM`, but less in HiLDA and TCSM (Fig S16, Section S3.3.4).

**Figure 3.**
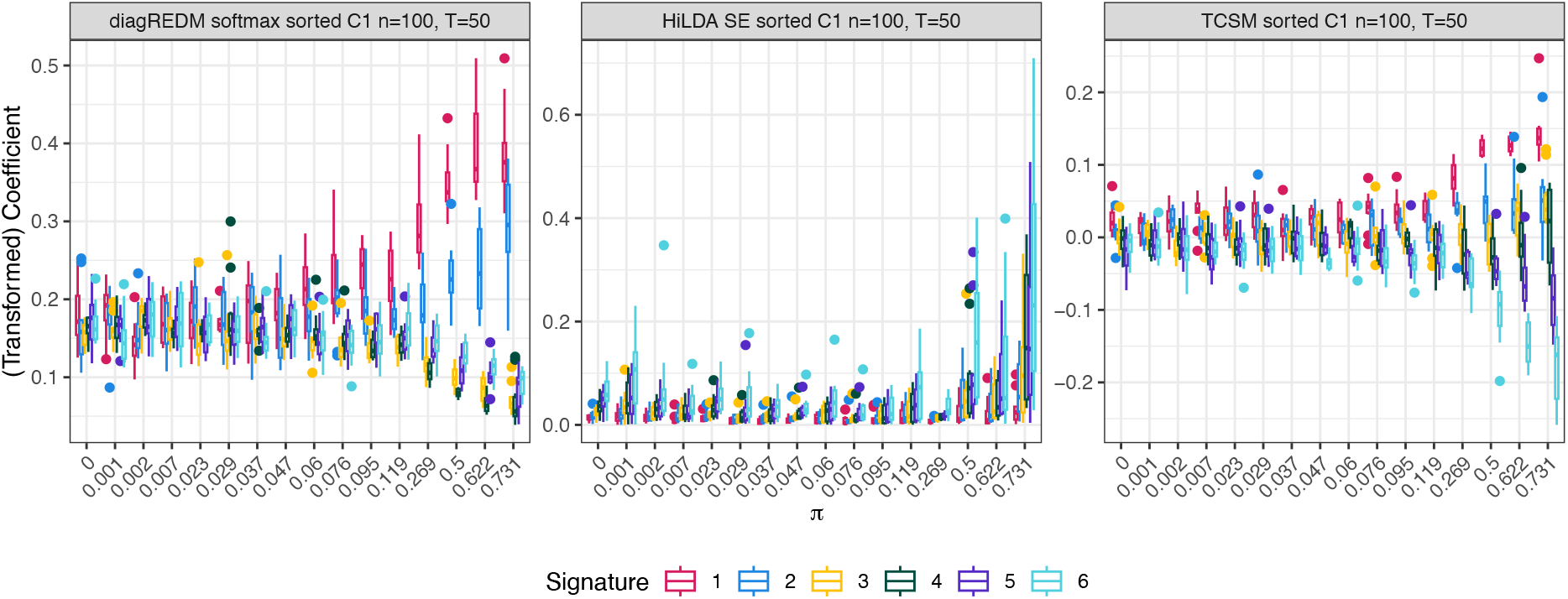
Comparison of coefficients that represent differential abundance for the models under consideration: `diagREDM`, HiLDA, and TCSM. Although `diagREDM` shows a marked increase in dispersion of softmax-transformed 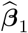 across signatures (indicating differential abundance) and the coefficients of TCSM show an increased deviation from zero (also indicating differential abundance), the results from HiLDA do not show such trends unless the mixing proporion *π* is very high.

#### 2.4.3 Effect of active signature selection on differential abundance

Unlike in HiLDA and TCSM, where a joint *de novo* signature extraction and differential abundance test is performed and the number of active signatures is a given parameter, the models suggested here require signature exposures to have been derived from a known set of active signatures prior to parameter estimation. The selection of active signatures is a topic of debate and careful thought is put in choosing the set of active signatures in each cancer type, or even sample. By generating simulated signature exposures using different strategies of active signature selection, we assess the robustness of our differential abundance model `diagREDM`. We find that, unless the number of active signatures is severely overestimated, leading to a high false positive rate, the differential abundance results are comparable across active signature selection strategies (Fig S17, Section S3.4).

## 3 Results

### Differential abundance of nucleotide changes

The type of mutations created at clonal and sub-clonal stages differs in distinct ways between cancer types. These divergent changes can already be observed at nucleotide level. To assess whether any changes in mutational aspects of clonal and sub-clonal mutations would be perceptible at the highest level of point mutation characterisation (i.e. into six substitution categories), we fit the Dirichlet-multinomial `fullREDM` model on data categorised by these single-nucleotide mutations. This allows us to determine general trends of mutation-type co-occurrence and differential abundance. Fig 4 shows 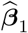, the estimate of ***β***_1_, for each cancer type. Each element of 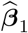 corresponds to the log-ratio of a mutation type {`C>A`, `C>G`, `C>T`, `T>A`, `T>C`} with respect to baseline category `T>G`, as obtained by our estimator. These elements are colour-coded and joined by a line, to enable the identification of similar trends of differential abundance across cancer types. Skin-melanoma offers an example of a cancer type with clear differential abundance in which the interpretation of coefficients is interesting: `C>T` appears to increase the least (or possibly to decrease) from clonal to subclonal mutations, followed by the same fold-change in three mutation categories: `T>A`, `T>C` and `T>G` (the baseline), and followed by the highest fold-change in `C>A` and `C>G`. Pan-cancer, the estimated ***β***_1_ coefficients for `C>T` (with respect to baseline category `T>G`) are generally not correlated with other coefficients across cancer types, whereas those of `T>C` and `T>A` are highly correlated (i.e. their lines follow each other, often sharing the same 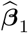). These results suggest patterns of differential abundance between clonal and subclonal mutations even at the lowest level of granularity, and we extend this approach to mutational signatures to be able to interpret the results further.

**Figure 4.**
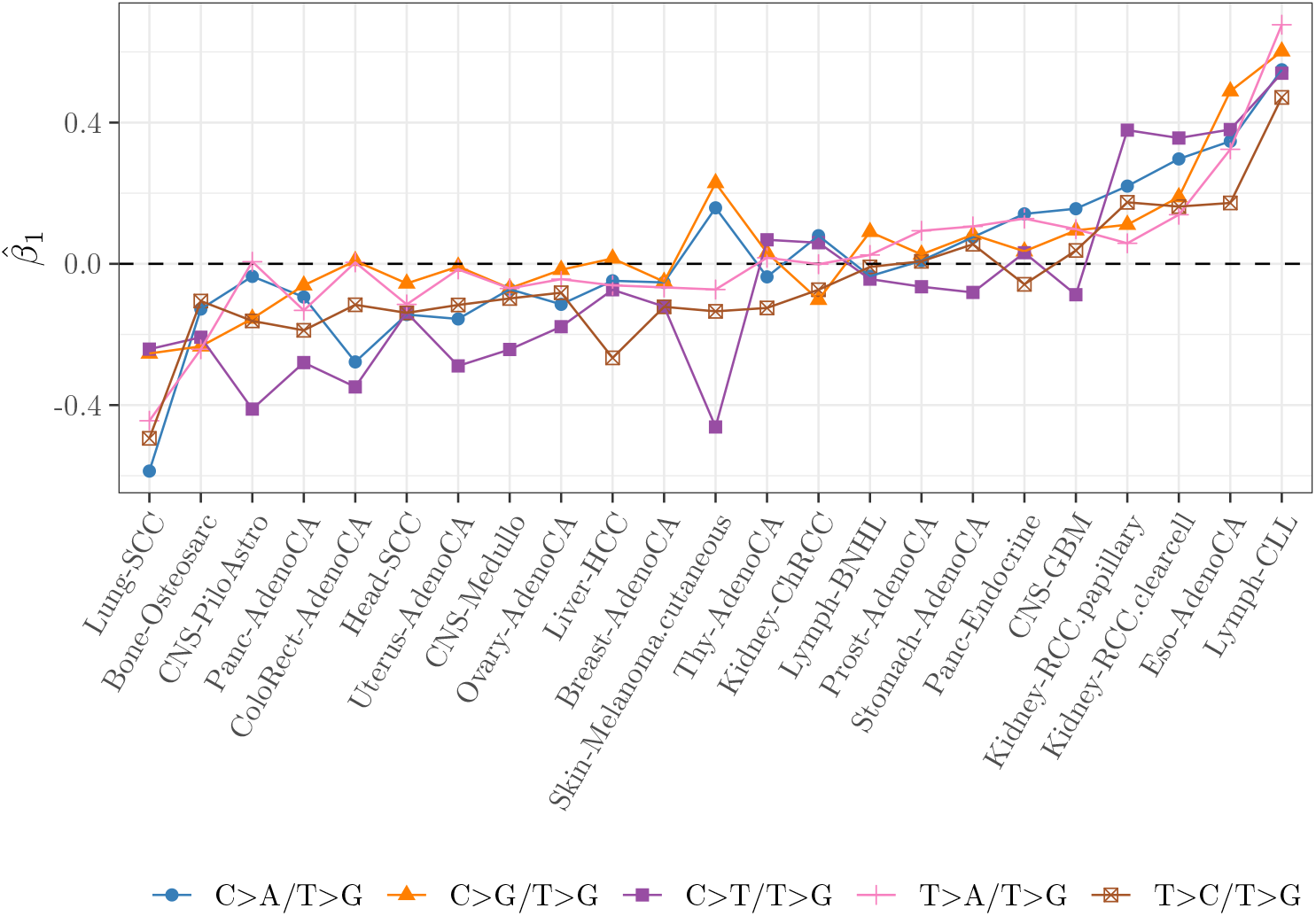
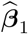 estimated using `fullREDM` on single-nucleotide mutations in 23 cancer types of the PCAWG cohort, using the mutation category `T>G` as baseline.

### Differential abundance of mutational signatures

In the remaining results of PCAWG data, the models have been fit with the subset of signatures considered to be active in each cancer type, according to [14] (Table S5). Using the `diagREDM`, all cancer types are differentially abundant when including all signatures, and most (17/23) are when including only non-exogenous signatures (following Benjamini & Hochberg adjustment). The six cancer types that cease to be differentially abundant are CNS-Medullo, CNS-GBM, CNS-PiloAstro, Kidney-ChRCC, Lung-SCC and Stomach-AdenoCA. These six cancer types do not differ in the number of samples (Welch Two Sample t-test; *p*-value = 0.08733) nor in the number of active non-exogenous signatures (Welch Two Sample t-test; *p*-value = 0.1858, with an average of 8.5 signatures in the group of differentially-abundant cancer types and of 5.5 in the group of non-differentially abundant cancer types). Using the multinomial with full random effects, `fullREM`, suggests differential abundance in all cancer types. There is no difference in the average number of mutations constituting the observed exposures 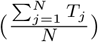 between differentially- and non-differentially abundant cancer types (*p*-value = 0.425, Welch Two Sample t-test on log_2_-transformed mutation toll), rather, it is the number of patient samples which can more strongly limit our ability to determine differential abundance, as discussed. The number of mutations for the patient-specific subsamples is found in Table S6. Plotting all 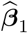 (Fig S18) or 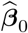 (Fig S19) gives a better indication of which signatures are behind these changes, as will be discussed.

### Differential precision of mutational signatures

Besides 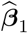 parameters indicating differential abundance, the models include group-specific overdispersion parameters, which may indicate differential precision. Fig 5 shows the log-transformed estimated precision parameters 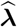, for the clonal and subclonal mutation groups in each cancer type. Higher values indicate higher precision, or lower overdispersion. In most cases (19/23) the subclonal mutations show higher overdispersion, or lower precision 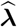, which is to be expected if subclones have different relative signature exposures or if patients diverge in the types of active mutational processes following cancer onset. There is a statistically-significant difference in precision in 12/23 cancer types (Breast-AdenoCA, CNS-GBM, CNS-Medullo, Kidney-ChRCC, Liver-HCC, Lung-SCC, Lymph-BNHL, Ovary-AdenoCA, Panc-AdenoCA, Panc-Endocrine, Prost-AdenoCA, Thy-AdenoCA; t-test with FDR adjustment). In all statistically-significant cases the precision is higher in the early mutation group. Head-SCC, Kidney-RCC (clear cell), Kidney-RCC (papillary) and melanoma are the cancer types where there is higher precision in the late mutation group, albeit not significantly. There are multiple additional reasons why overdispersion should be higher in subclonal than clonal mutations: the presence of subclones with distinct active mutational processes, signatures that reflect interactions between mutational processes in more advanced stages of cancer, or noise particular to mutations of lower coverages.

**Figure 5.**
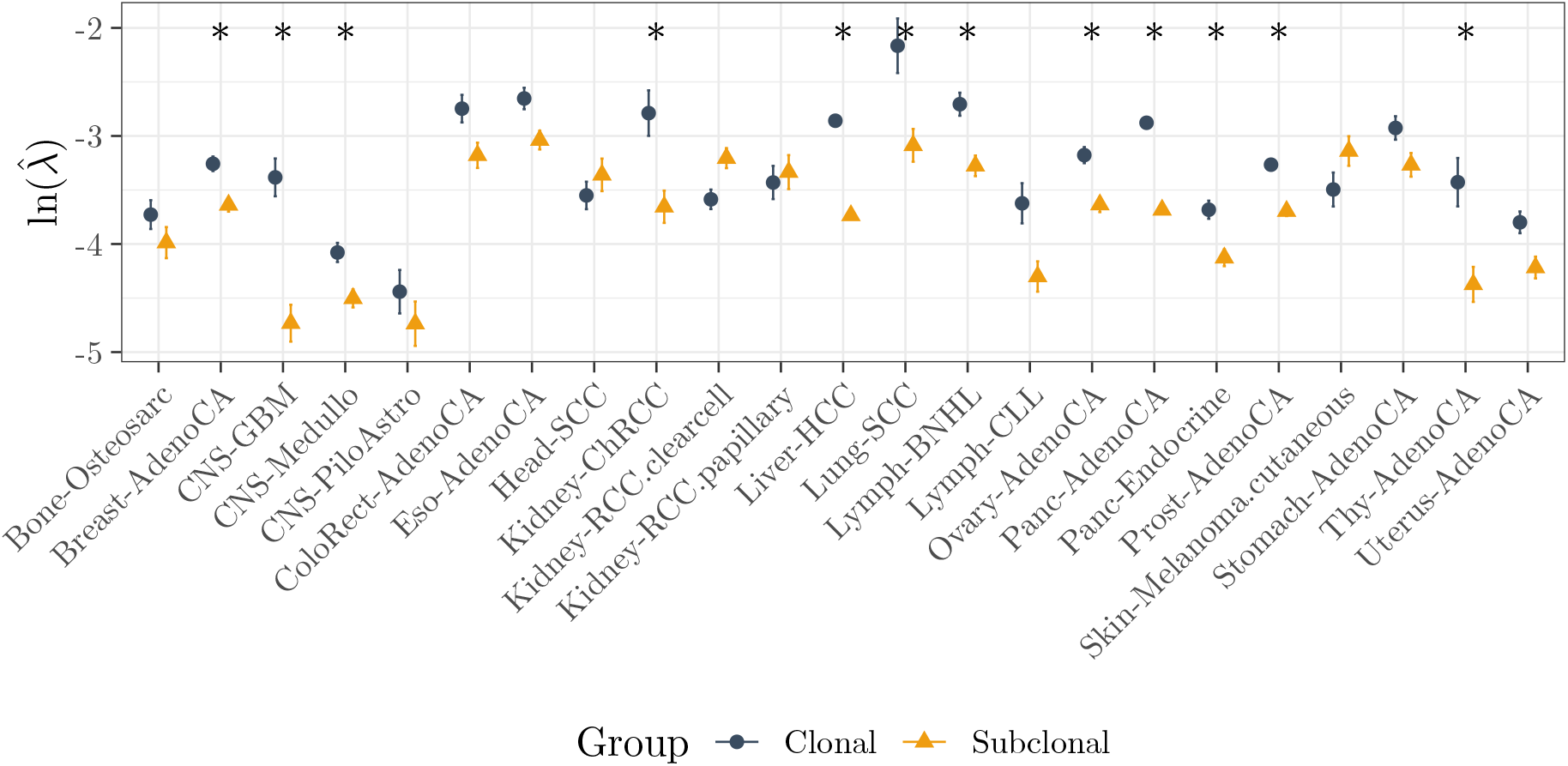
Estimated precision parameters ***λ***, for each of the groups, in each cancer type. Higher values indicate higher precision, or lower overdispersion. In most cases (19/23) the subclonal mutations show higher overdispersion rates, which are to be expected if subclones have different relative signature exposures. The bars indicate the *±* standard error. Asterisks indicate cancer types for which the precision parameters are statistically not the same (t-test, FDR correction).

### Identification of patient strata from their random intercepts

A third output of the model are the estimated patient random intercepts. TMB outputs their point estimates at the maximum (Fig S20), as well as values for the estimated correlation matrix if `fullREDM` is used (Fig S21). The correlation between two signatures in contributing to the random effects differs among cancer types, i.e. there is rarely a clear pattern of co-occurrence of signatures (see Fig S22 for selected pairs of signatures of interest). The extent to which patients can be categorised into strata according to their mutational signatures, based on the patient intercepts, also varies between cancer types: in Lymph-CLL at least two clear strata can be defined based on SBS9 (Fig S20), and so is the case in Prost-AdenoCA due to a range of signatures (but especially due to SBS3) whereas the Lung-SCC cohort appears much more uniform. The abundance of HRD signature SBS3 varies greatly among patients of Breast-AdenoCA, Eso-AdenoCA, Ovary-AdenoCA, Panc-AdenoCA and Prost-AdenoCA, which drives the patient stratification of these cohorts.

Having considered the cohort-wide results from the point of view of differential abundance, differential precision, and stratification based on the estimated random intercepts, we now focus on signature-specific results.

### Differential abundance results at the signature level

The coefficients 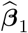 can give us an indication of how shared mutational processes behave in different cancer types. This is of particular interest in the signatures of unknown aetiology (u.a. henceforth). For this analysis, the vectors 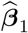 have been transformed using the inverse ALR transformation to be able to draw conclusions on the baseline signatures. These transformed 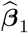 can still be used to compare two signatures between cohorts, as the log-ratios of abundance are preserved in the transformation. Importantly, correlations between 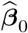 or 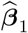 of signatures can reflect spurious and imposed correlations (e.g. a high correlation in signature abundance in softmax-transformed 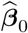 between two signatures across cancer types can be a reflection of the large variance in the abundance of a third signature, and the same is the case for 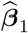).

Fig 6 displays scatterplots of softmax-transformed 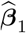 for pairs of signatures of interest (six pairs which include clock-signatures SBS1, SBS5 and SBS40, APOBEC signatures SBS2 and, SBS13, and two signatures of high abundance and active in multiple cancer types but with unclear aetiology, SBS8 and SBS18). In each plot, a point represents a cancer type. Tighter alignments along the identity line indicate a higher agreement in differential abundance, i.e. in the log-fold change in mutational signatures. Signatures that share the same 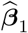 across cancer types, therefore, are hypothesised to share a biological mechanism. Outliers in an otherwise good agreement might indicate particularities of the relevant mutational processes in the specific cancer type.

**Figure 6.**
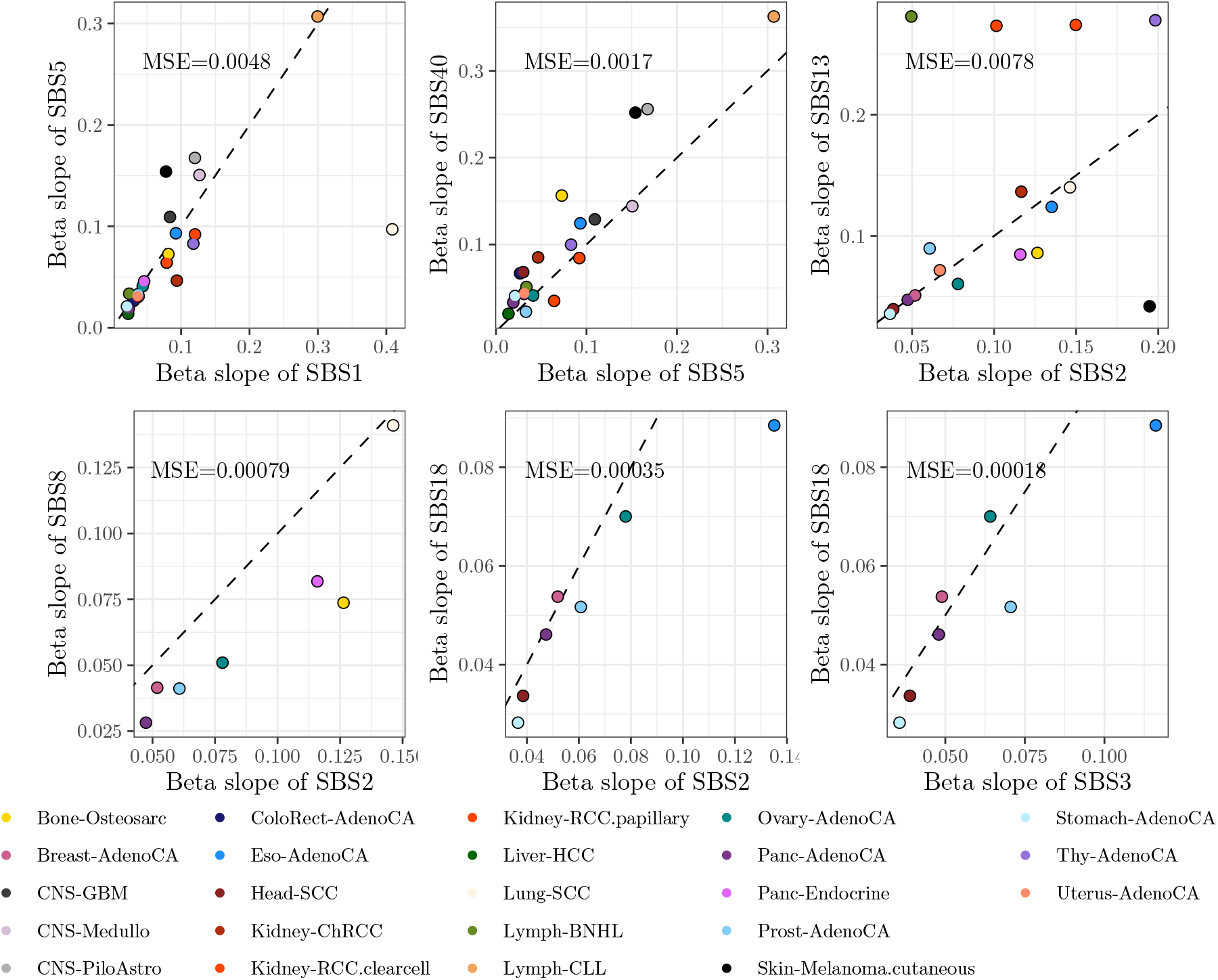
Selected pairs of signatures of interest and with constant ratios of (softmax-transformed) 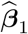. Each point represents a cancer type, and a tighter match along the identity line (the dashed line *x* = *y*) indicates a higher agreement in the differential abundance characteristics, i.e. a constant fold-change in their abundance. Only signatures which are shared between each pair of cancer types are shown in the plot. The mean squared error is shown for each pair of signatures. Note the very good matches between SBS1 and SBS5 (signatures of proposed constant mutation rate) and the consistently lower values for the softmax-transformed 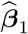 of SBS2 compared to SBS8.

### Concordance of clock signatures SBS1, SBS5 and SBS40

Reassuringly, the agreement of 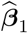 of known age-related signatures SBS1 and SBS5 [31, 14] is generally good. SBS40, sometimes described as an age signature, and a pervasive signature of high abundance, has 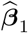 that match with those of SBS5. In the case of SBS1 and SBS5, an outlier (Lung-SCC) can be seen in seen in the first facet of Fig 6. Particularly, 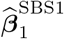 is much higher than 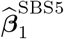, and SBS1 is found in very low abundance (Fig S19) – these two pieces of information warrant the question of whether SBS1 should be considered inactive in the PCAWG Lung-SCC cohort. In no other cancer type is SBS1 the signature of least abundance.

A less acute disagreement among clock signatures is found in the three CNS cancer types, where 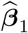 for SBS5 are too different from those of SBS1. In the three CNS cancer types, only signatures SBS1, SBS5 and SBS40 are are present in any two cancer types, and for all three the pattern of differential abundance shows the same relative increase: the 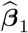 of SBS1 is always slightly lower than 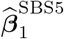, and all remaining signatures have a higher 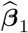. Whereas in CNS-Medullo SBS1 and SBS5 correlate in the patient intercepts (Fig S20-S22), they do not in the other two cancer types. In CNS-GMB SBS1 has a constant value of zero in the intercepts (i.e. no between-patient variability in the abundance of SBS1 with respect to SBS40). Contrarily, in all CNS cases the ratio of SBS5 and SBS40 varies greatly between patients. This finding applies across cancer types: we find that SBS1, SBS40 and SBS41 often do not contribute to the between-patient variability (see Kidney-RCC.papillary for the clearest example) whereas SBS5 does. These results are inconsistent with hypothesis that the random intercepts of SBS1 and SBS5 roughly track the age of the patients in all cancer types. Keeping in mind the discordant behaviour of SBS1 and SBS5 in the aforementioned cancer types, the comparison of any 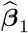 to 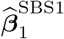 and 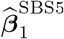 in the same cancer type is useful (Fig S23): where the belief that SBS1 and SBS5 represent signatures of constant mutation rate is justified, we can compare 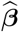 of signatures of interest with 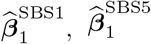, to determine the direction of change (i.e. an increase or decrease in the averaged mutation rate).

### Coordinated and separate behaviour of APOBEC signatures

Overall, there is a good agreement in 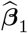between the two APOBEC signatures SBS2 and SBS13, with clear discordant 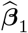 in the two renal cell carcinoma cohorts (Kidney-RCC clearcell and papillary), as well as in Lymph-BNHL and Skin-Melanoma. These four discordant samples are the only ones in which the patient random intercepts of SBS2 and SBS13 are poorly correlated, with correlations ranging from -0.03 to 0.36 (Fig S22), indicating that patients who present high values of SBS2 do not present high values of SBS13, or conversely. APOBEC signatures have been considered to be active [14] or not [32] in previous studies of kidney cancer. Their low abundance, discordant 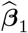, and discordant contribution to the random intercepts suggest that, in these cancer types, SBS13 might not be. Judging the equivalent results in melanoma, SBS13 is active but SBS2 is not.

### General increase in signature activity

Fig 7 plots the fraction of cancer types in which 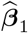 of a given signature is greater than 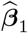 of every other signature, for each signature. This indicates the rank of 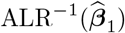 of the given signature. Therefore, analysed row-wise, a signature with many blue entries corresponds to a signature which has, consistently, the highest softmaxed-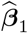 across cancer types, suggesting signatures that preferentially create subclonal mutations, or the mutation rate of which increases in subclonal stages of tumour development. On the other extreme, green rows indicate signatures with the lowest softmaxed-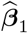, most likely indicating signatures which decrease in abundance over the course of tumour development, or the mutation rate of which remains constant. Signatures clustered according to the dendrogram on the right of the plot are suggested to share temporal dynamics. SBS1 and SBS5 do not rank differently across cancer types. SBS17a (u.a.) has higher coefficients than SBS17b (u.a.) in all occasions, and they in occasion rank lower than SBS1/5. The rank of SBS40 can vary extraordinarily, implying that, as a very flat signature, it could be capturing signal from other signatures. Some signatures tend to have the highest coefficients: SBS20 (POLD1 mutations), SBS31 (platinum chemotherapy), SBS33 (u.a.), SBS43 (artefact), SBS53 (artefact). On the other hand of the spectrum, SBS1 and SBS5 have comparatively low coefficients, which would indicate that there is an increase in the averaged mutation rate from clonal to subclonal stages in most mutational signatures. Although generally low, 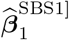 and 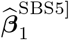 are not necessarily the lowest: signatures of undoubtedly exogenous origin (SBS7b and SBS7d, representing UV light) have consistently lower coefficients, as do SBS12 (u.a.), SBS15 (DMMR), SBS16 (u.a.), and SBS19 (u.a.), suggesting, whenever their aetiology is unknown, that they are, too, signatures of exogenous processes. SBS16 appears as clonal mutation – suggesting that the mutational process it represents was active historically, before tumour onset – in the two cancer types where it is present, Head-SCC and Liver-HCC. It has been linked to alcohol consumption [33] but this is not included in COSMIC [9]. This is the signature that solely drives the random intercepts in Head-SCC (Fig S21) indicating patient variability in its abundance, stratifying the samples into three groups according to these random intercepts (Fig S20).

**Figure 7.**
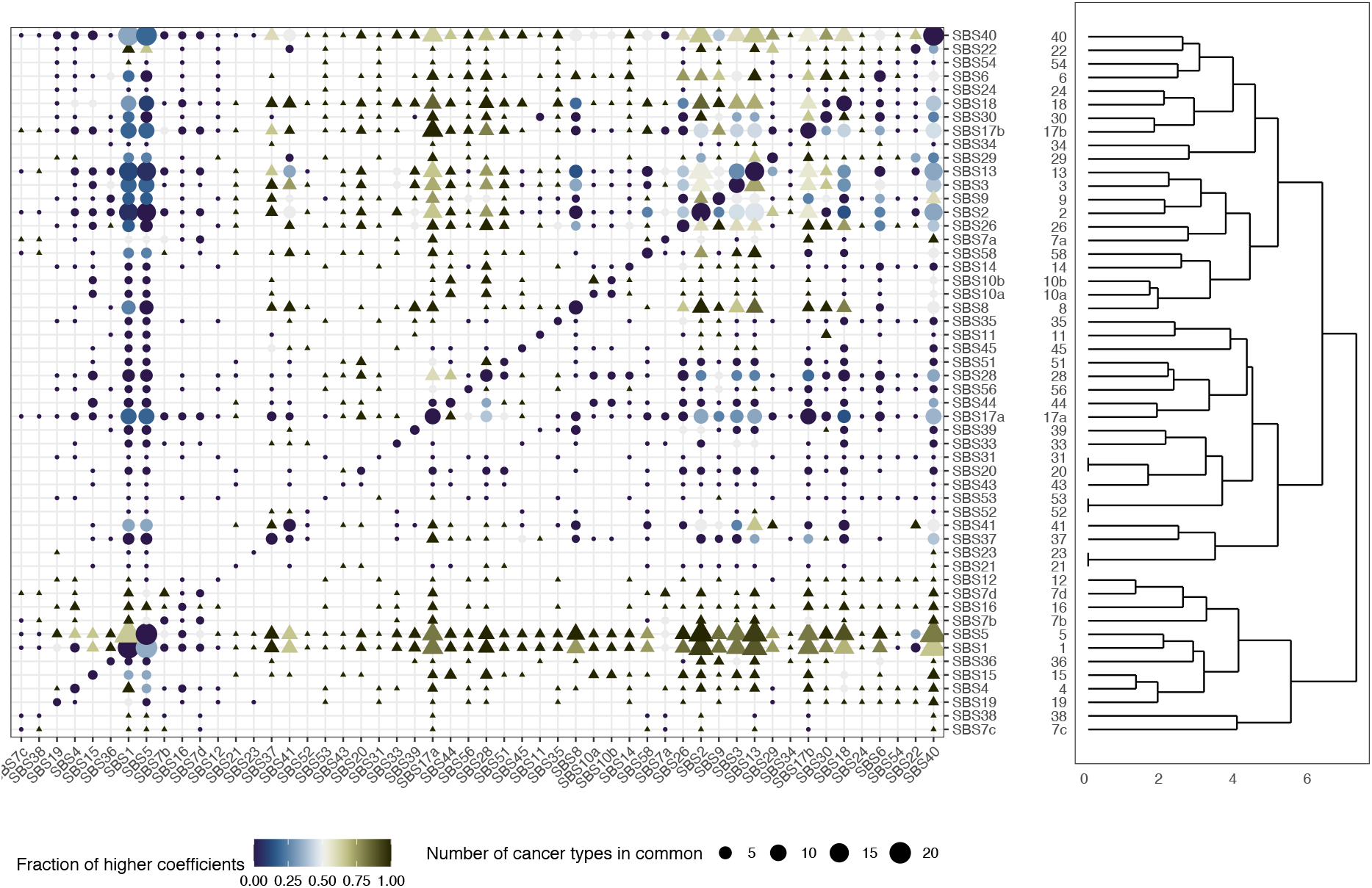
Fraction of cancer types in which the 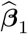 of any signature (in the *x* axis) are higher than those of another signature, for every other signature along the *y* axis. The vertical clustering is done on Euclidean distance using complete linkage. The size of the dots indicates the number of cancer types in which both signatures are active. The elements in the diagonal have a shared value of 0, and their size indicates the number of cancer types in which the corresponding signature is active.

### New insights on signature dynamics

Beyond the expected agreements, the most noteworthy trends are those of SBS8 and SBS18 and their interplay with APOBEC signatures and HRD. The aetiology of SBS8 is uncertain, although it has been linked to HR and NER deficiency [9]. Although SBS18 has been linked to reactive oxygen species, its aetiology is not yet clear, and is associated with MUTYH-related signature SBS36 [9]. SBS18 aligns well with the APOBEC signatures except in the case where 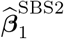 is highest (Eso-AdenoCA). Similarly, 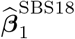1 closely matches 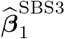. The 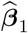 of SBS18 closely matches 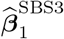. SBS3 and SBS8 display markedly different 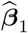(Fig S24), although they always have intermediate values of 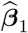 compared to other signatures. Interestingly, SBS2 shows consistently higher 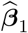 than SBS8. Such behaviour could be explained by at least four scenarios: by APOBEC initiation preceding a heightened mutation rate of SBS8, by a more subtle increase in the mutation rate of SBS8 than of SBS2, by an increase in SBS8 in only a fraction of clones (but an increase of SBS2 in all cells), or by an increase of SBS8 in a fraction of the patients (and a more common increase of SBS2). With very few exceptions, these signatures have intermediate values of 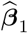. In fact, in several cancer types we find several shared values of 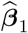 (seen as a plateau in the sorted 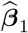 of Fig S18). In Panc-AdenoCA one such plateau includes SBS2, SBS13, SBS3 and SBS18 - these signatures are also included in the same plateau in Breast-AdenoCA. In the vast majority of cases such a plateau of 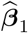 contains values only marginally higher than 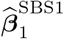 and 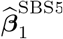, and is followed by a much lower number of signatures of very high 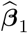.

### Tissue-specific differential abundance

Within-tissues analysis reveals which aspects of differential abundance are shared between cancer types of the same tissue and which are not. Often, the set of active signatures itself varies drastically within a tissue or organ (in the case of the three CNS samples, the two pancreatic samples, between Kidney-ChRCC and the two Kidney-RCC samples). This is to be expected in cases where the cell of origin differs.

With the pre-selected set of active signatures, CNS samples differ in every aspect except in the correlation between SBS1 and SBS40 in CNS-GBM and CNS-Medullo (Fig S21) – a correlation which is absent in CNS-Medullo – and the ordering of 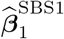 and 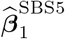. Indeed, we find that upon re-extraction of signature exposures and considering all signatures present in the tissue (Fig S25) the three CNS samples show only moderate commonalities in abundance 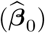 and none in differential abundance 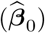. The two kidney RCC samples are very consistent in their differential abundance, showing SBS22 – the aristolochic acid exposure [9] – as the signature of lowest 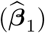, together with SBS40, and followed closely by the clock signatures. Upon signature re-extraction with all signatures present in the kidney, the general abundance is moderately similar even between ChRCC and RCC, but the differential abundance results differ completely. In the two RCC cancer types the opposite is true: their abundances differ slightly (e.g. in SBS29), but their differential abundance results are in perfect alignment except for, again, SBS29. In the case of Panc-AdenoCA, the random intercepts are clear in showing that, unlike in Panc-Endocrine, SBS3 plays an important role in explaining the variability between patients. In the analysis of all signatures, find a similar pattern to the ChRCC/RCC comparison in the pancreatic samples where, although the abundances are only markedly different in SBS3, SBS9 and SBS36 (showing shared and possibly tissue-related mutational processes that were active early in tumour development, or prior to it), the differential abundance patterns are not shared.

## 4 Discussion and deployment of *CompSign* in cancer genomics

Applied to the PCAWG dataset, the mixed-effects Dirichlet-multinomial model finds ubiquitous differential abundance between clonal and subclonal mutations, capturing similar behaviour in signatures known to be related, as well as discovering new correlations between signatures with regard to their temporal dynamics. Besides differential abundance, we find differential precision, with higher dispersion levels in subclonal signature exposures. In this work we begin to explore the true correlations between mutational signatures, and not the correlations induced by the compositional nature of these data. Validation of these results either experimentally or computationally and with orthogonal data would be of interest, and we welcome the recent papers in which mutational signatures are studied using *in vitro* models. The models can help answer a variety of questions in cancer genomics. To help in the deployment of these methods, we give some guidance for each step of the pipeline.

The signature exposure matrix **Y** is taken as input, but several choices need to be made in order to generate this matrix from a sequence of mutations, chiefly whether to extract signatures *de novo* – in which case the number of signatures will specified, possibly by computing the optimal number of signatures based on metrics (see e.g. [34]) – or to fit the signatures from a known set of signature definitions, such as the COSMIC signatures. The latter approach is simpler computationally and has the advantage that signature definitions and aetiologies are curated constantly in the COSMIC database, aiding in the interpretation of the output of the model. We show that the differential abundance results of the model are robust to the strategy used for signature extraction, but ultimately the signature extraction method is deserving of a conversation in its own right and needs to take into consideration the cancer type, the origin of the mutation data, and a variety of technical and clinical covariates. It is important not to include signatures which are nearly always zero. Although the multinomial model and its extensions support zero (count) exposures, including a signature which is never active or only active in the clonal and subclonal group can lead to problems in estimating ***β***_0_ and ***β***_1_. Alternative methods in the field, such as HiLDA and TCSM, have opted for extracting signatures *de novo* and determining differential abundance in a joint model. Indeed, one of the limitations of the models shown here is that there is no error propagation between signature extraction and differential abundance testing. On the other hand, in our models it is the incorporation of random effects, dispersion parameters and freedom in the choice of input signature exposures that allow for an in-depth characterisation of the differential abundance parameters.

The next step is the choice of model from the models considered here (`diagREDM`, `fullREDM`, `fullREM`). We recommend the use of `diagREDM` for most purposes, especially if the interest is in the analysis of differential abundance (and not, for instance, in the patient random effects that contribute to signature abundance). Especially, in cases where the number of samples is low and the number of mutations is high, estimating the covariance matrix in `fullREDM` can be difficult and slow. The R package includes additional models, not discussed here, where the number of dispersion parameters are either lower (i.e. |*λ*| = 1) or higher (patient-specific) than those discussed here, and can be suitable in cases where the number of subsamples per patient is high, or the number of covariates greater than two or continuous – for instance, in a regression setting where signature abundances are correlated with age or compared across chromosomes. The choice of baseline signature can be important for interpretability of the results. If using COSMIC signatures, we recommend using SBS1, SBS5 or SBS40 as baseline signatures, as they are active in most cancer types (simplifying between-cancer type comparison of coefficients), and they are suggested to be signatures of constant mutation rate. If there is good reason, these exposures could even be amalgamated together, and used as a single baseline category.

Finally, although the models give differential abundance results as a single *p*-value, they also provide the 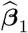coeficients, as well as all other coefficients in the model, which can be analysed in their own right. To account for the compositional nature of the model we suggest being careful in drawing any conclusions on the direction of change of signatures, and instead plotting the 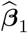 coefficients with their error and in increasing order, which gives an indication of which signatures experience a higher increase relative to each other, and makes clear which signatures lie at the extremes of the plot. In the same plot, it can be useful to see the position of the 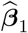 coefficients for any signatures of interest relative to the position of 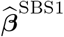 and 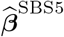, if those are not used as the baseline.

Immediate future directions along this line of research are to apply the proposed models to other cancer cohorts and extending the covariates considered - for instance, considering several timepoints in tumour development instead of two, interrogating chromosomes separately, testing the relevance of clinical covariates, or comparing exposures of samples with and without certain driver mutations. The models presented are publicly available and readily applicable to other types of latently compositional count data to determine differential abundance between groups, or perform other types of multivariate regression.

## Acknowledgements

L.M.G. was fully supported by the Wellcome Trust PhD programme in Mathematical Genomics and Medicine grant RG92770. D.-L.C. was partially supported by the Cancer Research UK grant C9545/A29580. Parts of this work were funded by Cancer Research UK core grant C14303/A17197 and A19274, which supported F.M. The funders had no role in study design, data collection and analysis, decision to publish, or preparation of the manuscript. We thank the patients and their families, without whom this work would have not been possible.

## Supplementary figures

**Figure S1:**
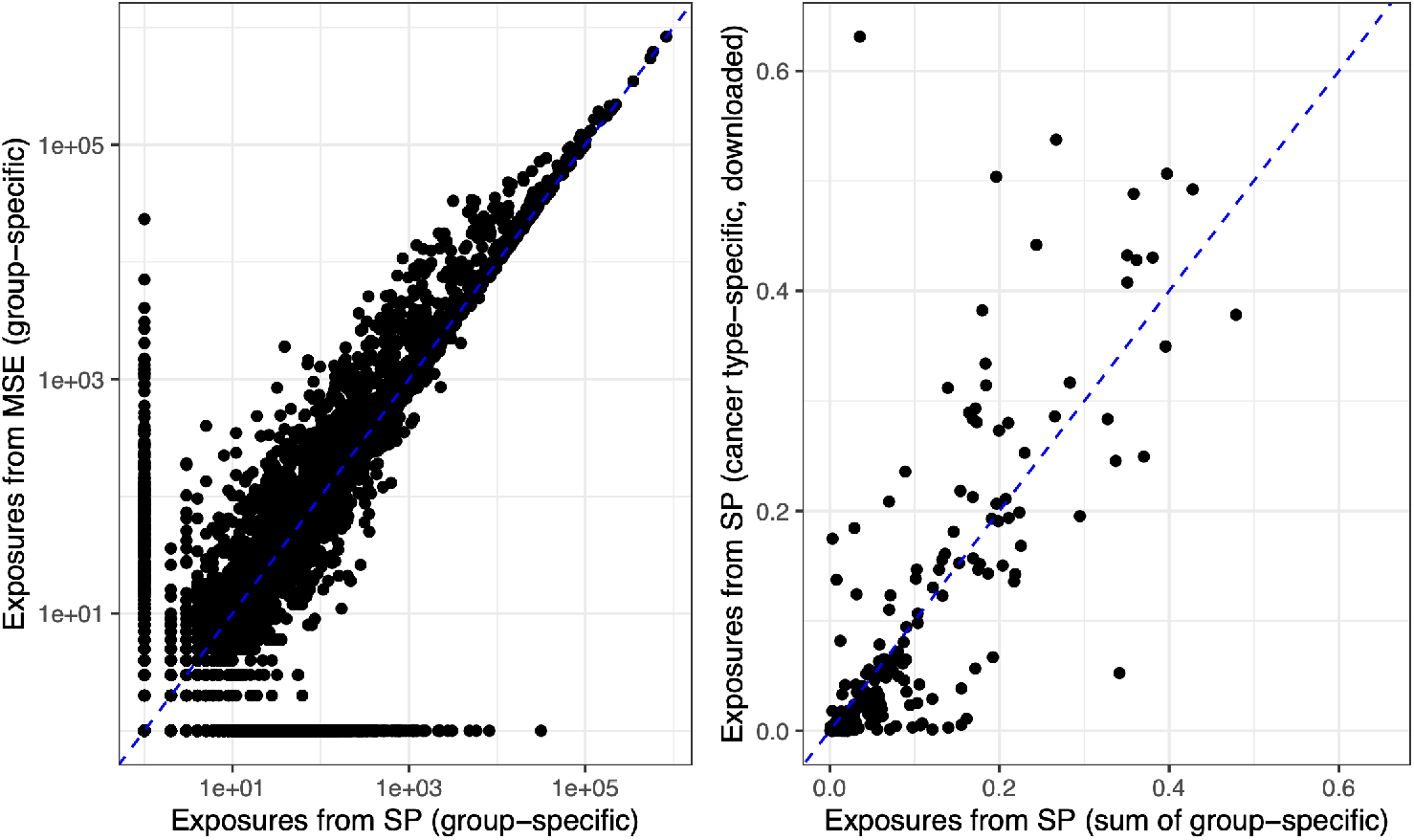
Left: comparison of patient-subsample exposures that have been extracted using quadratic programming (QP) and MSE. Right: comparison of exposures at the cancer-type level to those published in [14], in which each point represents a signature in a cancer type.

**Figure S2:**
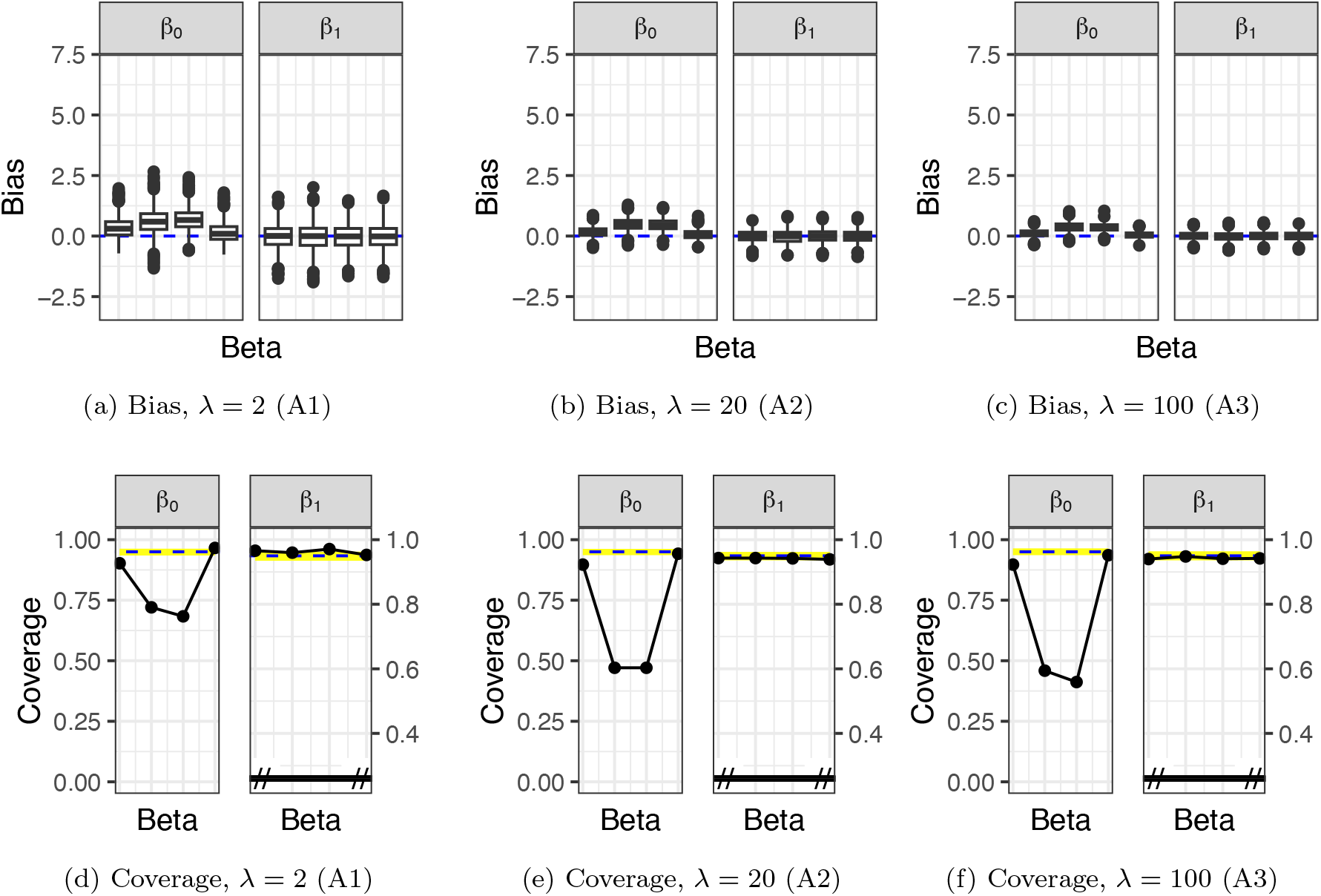
Estimation and inference when (wrongly) assuming a restricted model with non-correlated mixed-effects DM (`diagREDM`). Data simulated using a correlated DM with random intercepts with parameters *N*_*s*_ = 200, ∑_*j*_ *y*_*lj*_ = 180 *∀ l, d* = 5 (A1-A3). ***β***_1_ is well recovered, but ***β***_0_ is not, for the inability to model the correlated abundances of signatures in the first group. In all cases of overdispersion (all three columns) are ***β***_0_ poorly estimated, whilst the estimates of ***β***_1_ have no bias and good coverage. The highlighted area is the 0.025 *−* 0.975 quantile of 1000 binomial draws of the same size as the data with *p* = 0.95.

**Figure S3:**
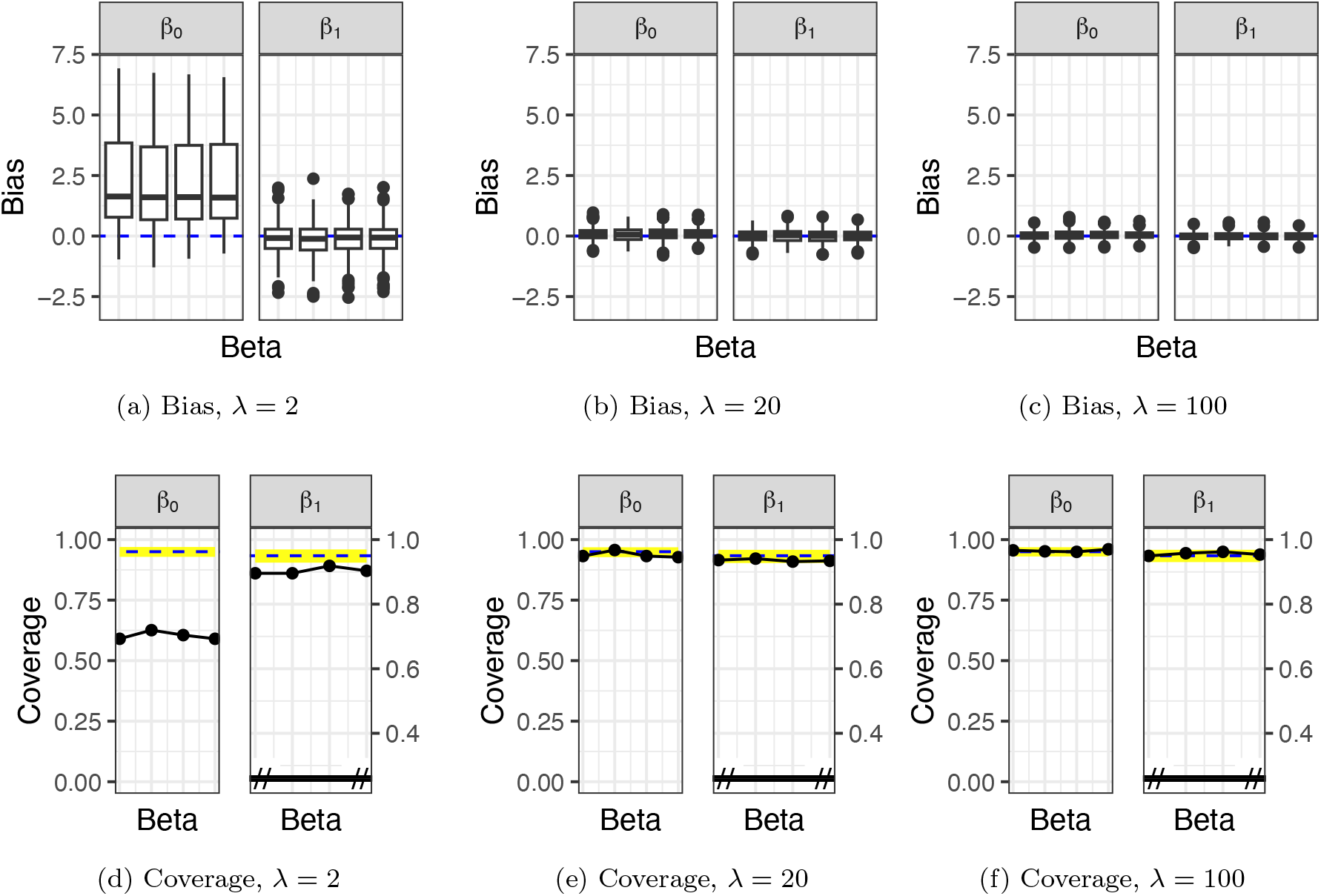
Estimation and inference when (correctly) assuming a model with correlated mixed-effects DM (`fullREDM`). Data simulated using a correlated DM with random intercepts with parameters *N*_*s*_ = 200, ∑_*j*_ *y*_*lj*_ = 180 *∀ l, d* = 5 (A1-A3). Both ***β***_0_ and ***β***_1_ are well recovered, except in the scenario of high overdispersion (first column; *λ* = 2). The highlighted area is the 0.025 *−* 0.975 quantile of 1000 binomial draws of the same size as the data with *p* = 0.95.

**Figure S4:**
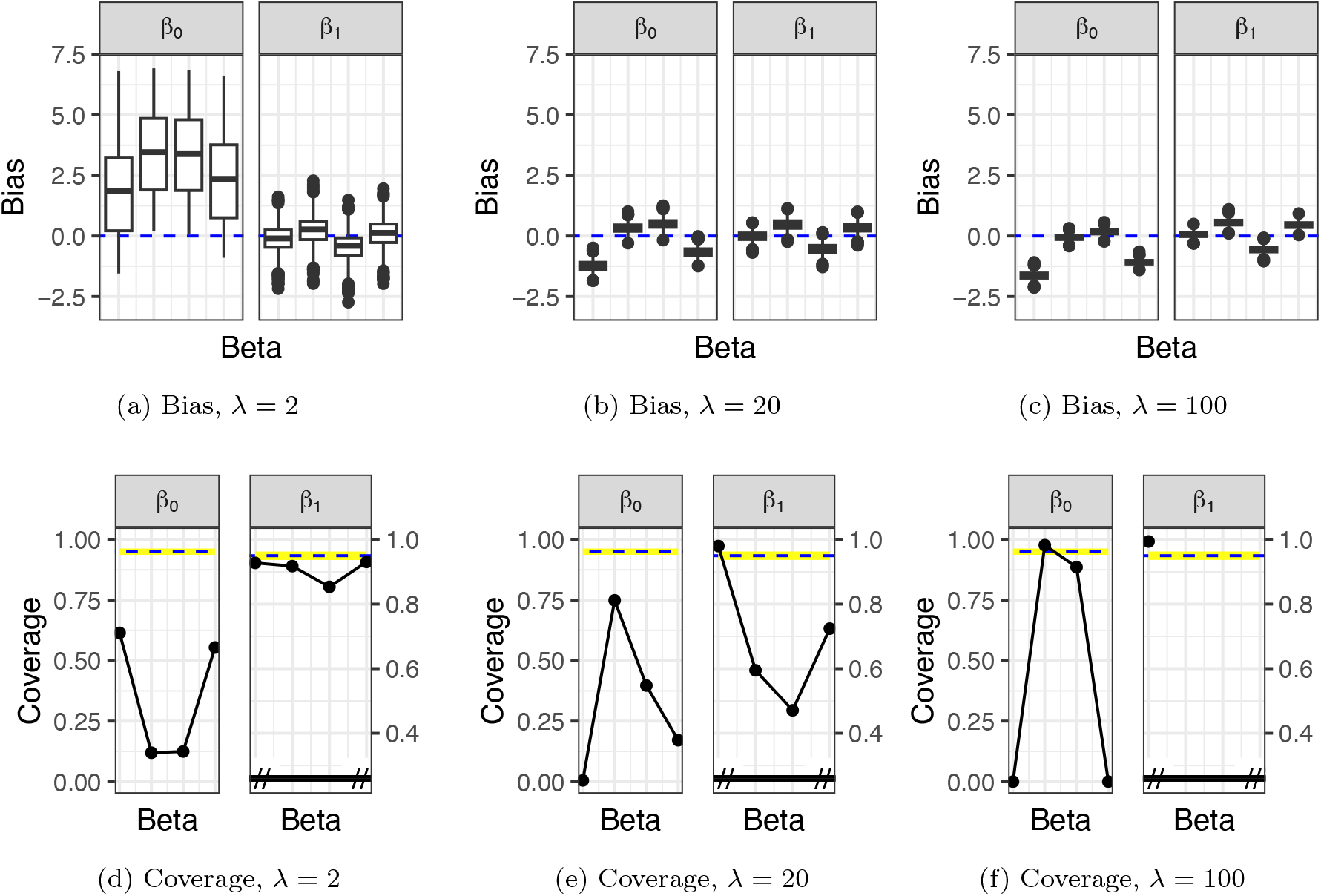
Estimation and inference when (wrongly) assuming a restricted model with a single intercept per patient (`singleREDM`). Data simulated using a correlated DM with random intercepts with parameters *N*_*s*_ = 200, ∑_*j*_ *y*_*lj*_ = 180 *∀ l, d* = 5 (A1-A3). Neither ***β***_0_ nor ***β***_1_ are well recovered, in all cases of overdispersion (all three columns). The highlighted area is the 0.025 *−* 0.975 quantile of 1000 binomial draws of the same size as the data with *p* = 0.95.

**Figure S5:**
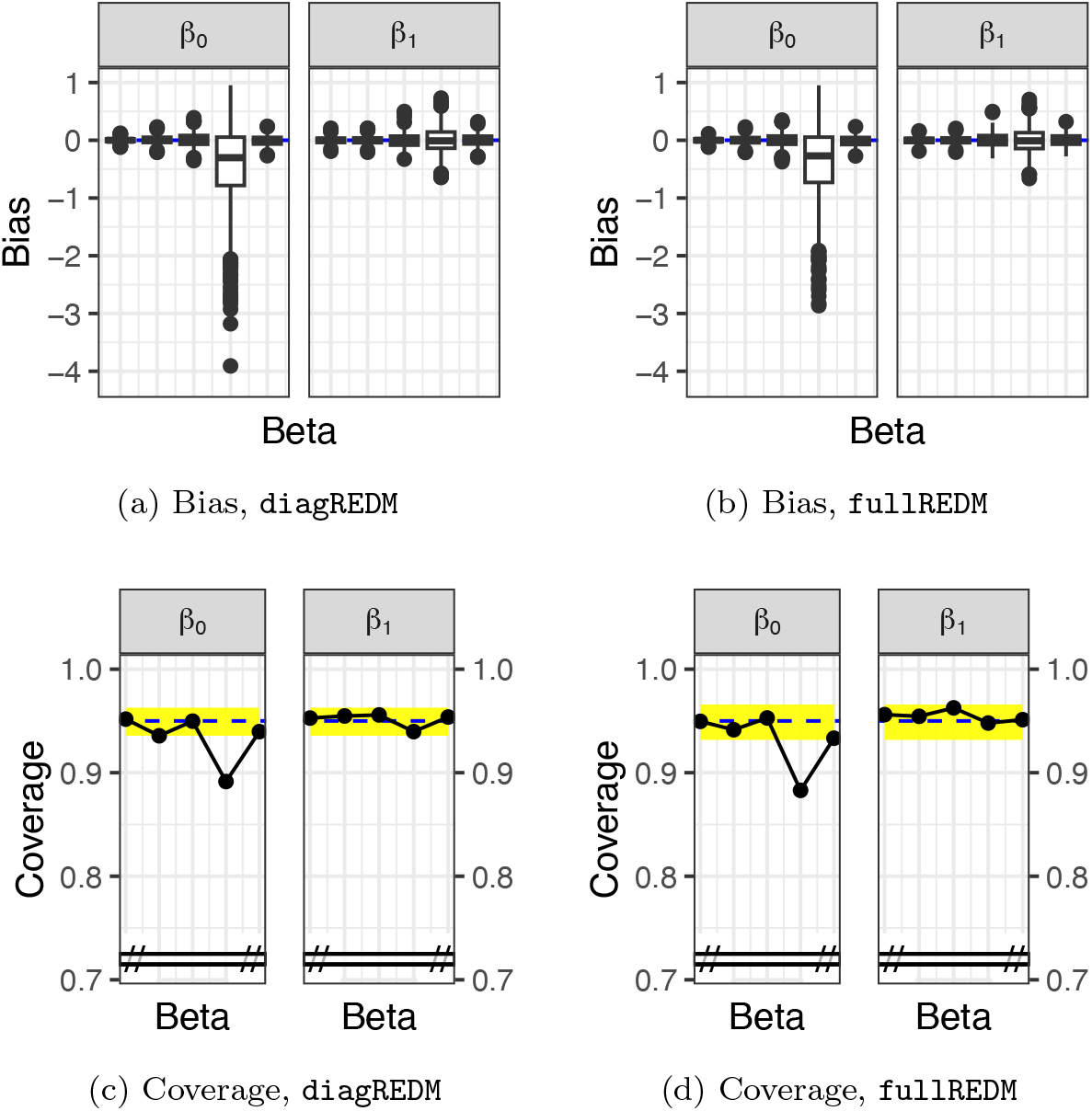
Estimation and inference using un-correlated mixed-effects DM (`diagREDM`) and correlated mixed-effects DM (`fullREDM`) for simulation (B1). Data simulated using previously-estimated parameters from the CNS-GBM cohort, with *N*_*s*_ = 200, no correlations, and a shared ***λ*** (average between the two estimated *λ*). All remaining parameters are taken as estimated. Given that uncorrelated data are simulated, both models give the same results for bias and coverage. Both ***β***_0_ and ***β***_1_ are well recovered, although the penultimate element of ***β***_0_ has some bias. The highlighted area is the 0.025*−*0.975 quantile of 1000 binomial draws of the same size as the data with *p* = 0.95.

**Figure S6:**
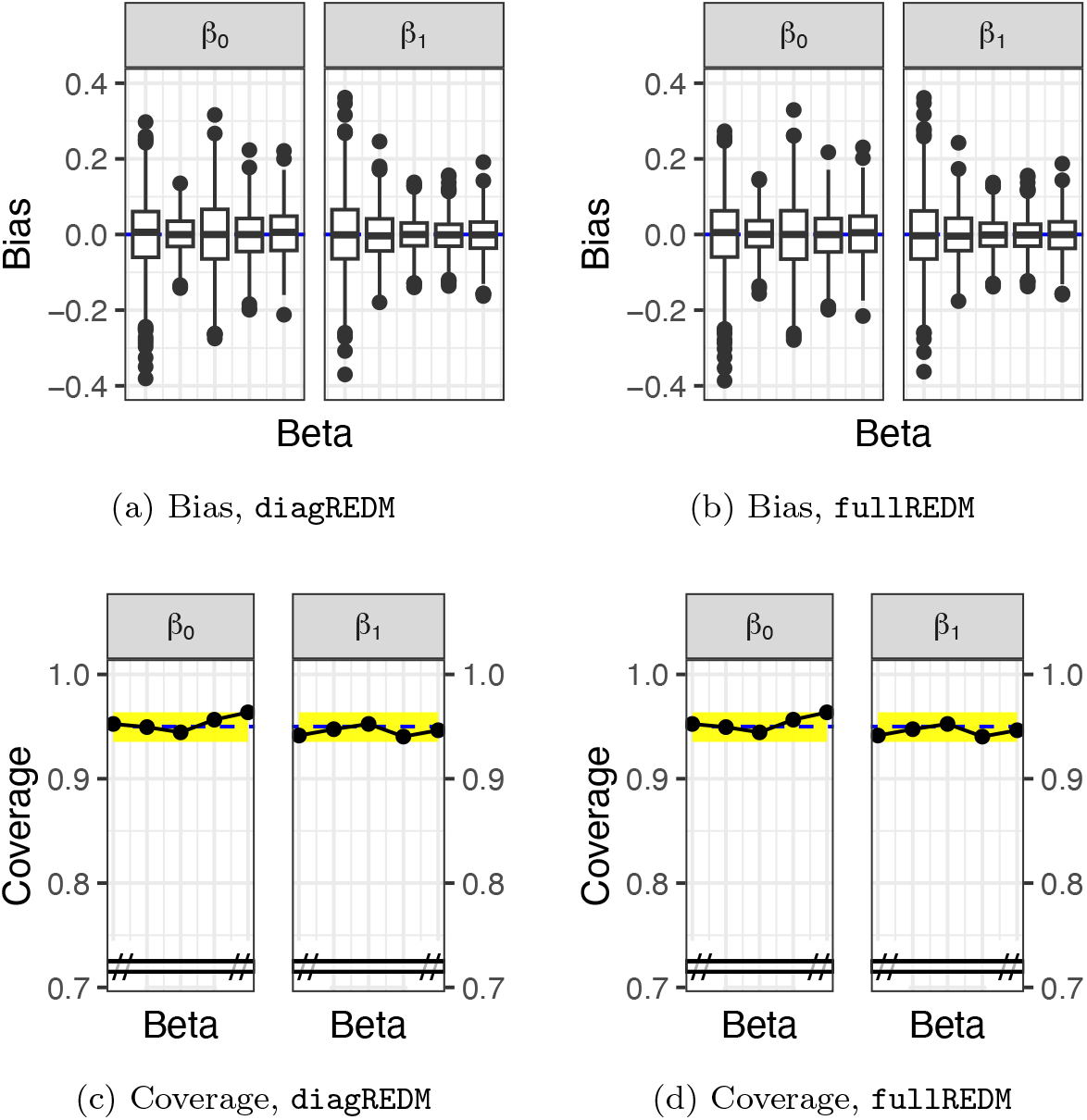
Estimation and inference using un-correlated mixed-effects DM (`diagREDM`) and correlated mixed-effects DM (`fullREDM`) for simulation (B2). Data simulated using previously-estimated parameters from the Lung-SCC cohort, with *N*_*s*_ = 200, no correlations, and a shared ***λ***. All remaining parameters are taken as estimated. Given that uncorrelated data are simulated, both models give the same results for bias and coverage. Bias and coverage results are satisfactory for both ***β***_0_ and ***β***_1_. The highlighted area is the 0.025 *−* 0.975 quantile of 1000 binomial draws of the same size as the data with *p* = 0.95.

**Figure S7:**
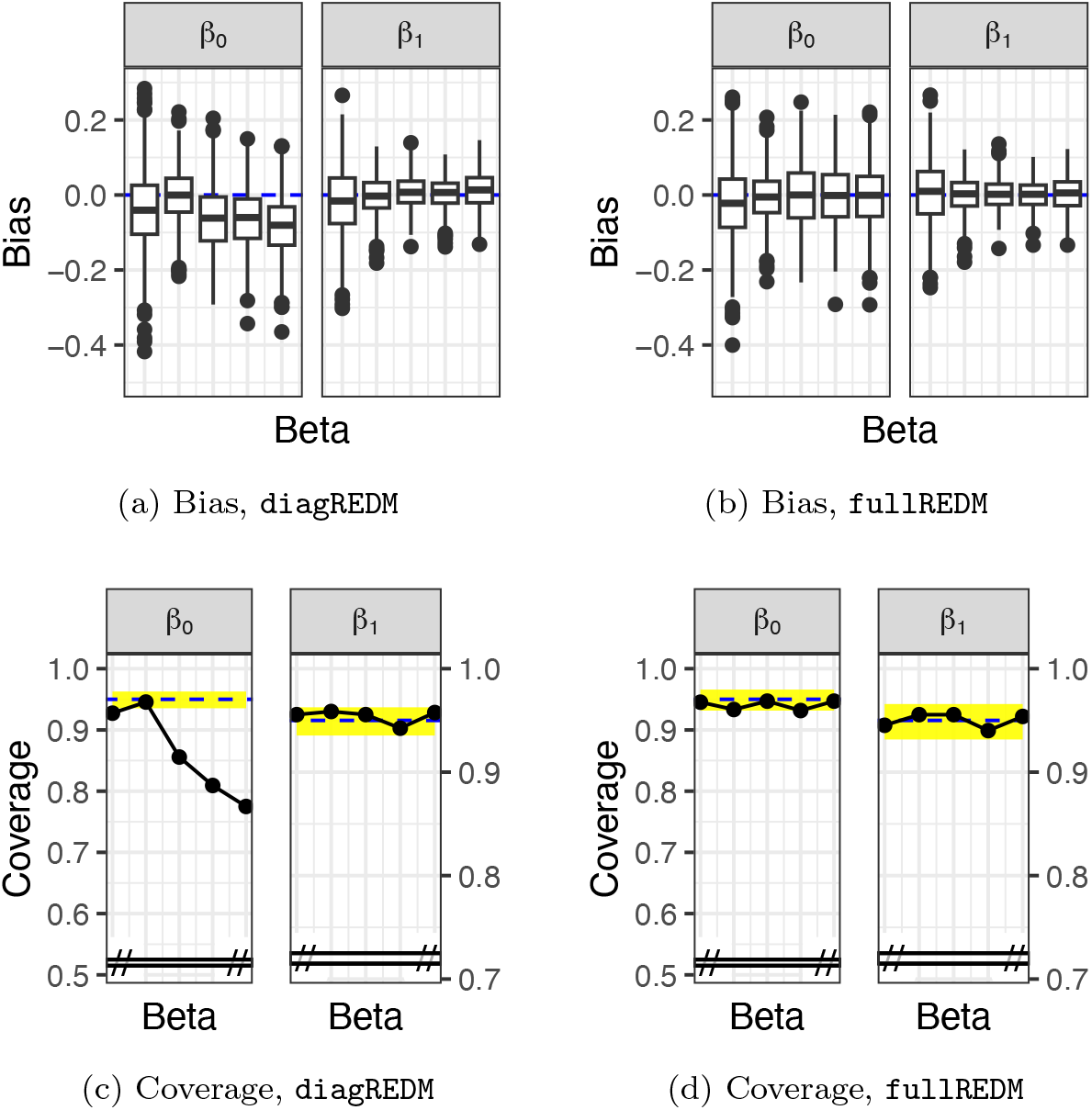
Estimation and inference using un-correlated mixed-effects DM (`diagREDM`) and correlated mixed-effects DM (`fullREDM`) for simulation (B3). Data simulated using previously-estimated parameters from the Lung-SCC cohort, with *N*_*s*_ = 200 and a shared ***λ*** (average between the two estimated *λ*). All remaining parameters are taken as estimated, including correlations. The results for bias and coverage differ between the models owing to the presence of correlations. Bias and coverage results are satisfactory in both cases for ***β***_1_, but there is bias and low coverage for ***β***_0_ in the non-correlated `diagREDM` model. The highlighted area is the 0.025 *−* 0.975 quantile of 1000 binomial draws of the same size as the data with *p* = 0.95.

**Figure S8:**
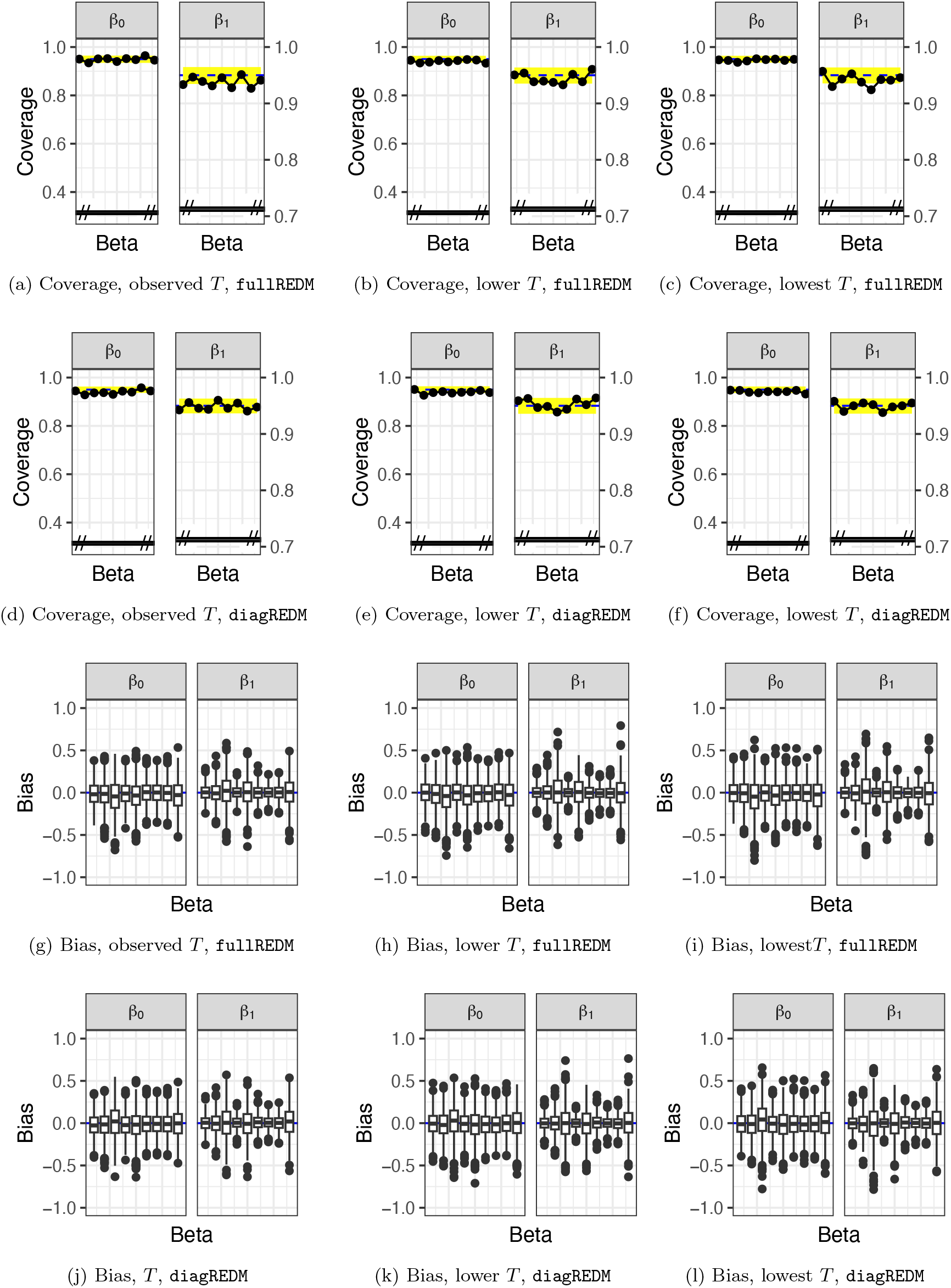
Bias and coverage as the number of mutations *T* in the simulations decreases (B4). Data simulated using previously-estimated parameters from the Prost-AdenoCA cohort, selecting the four most abundant signatures.

**Figure S9:**
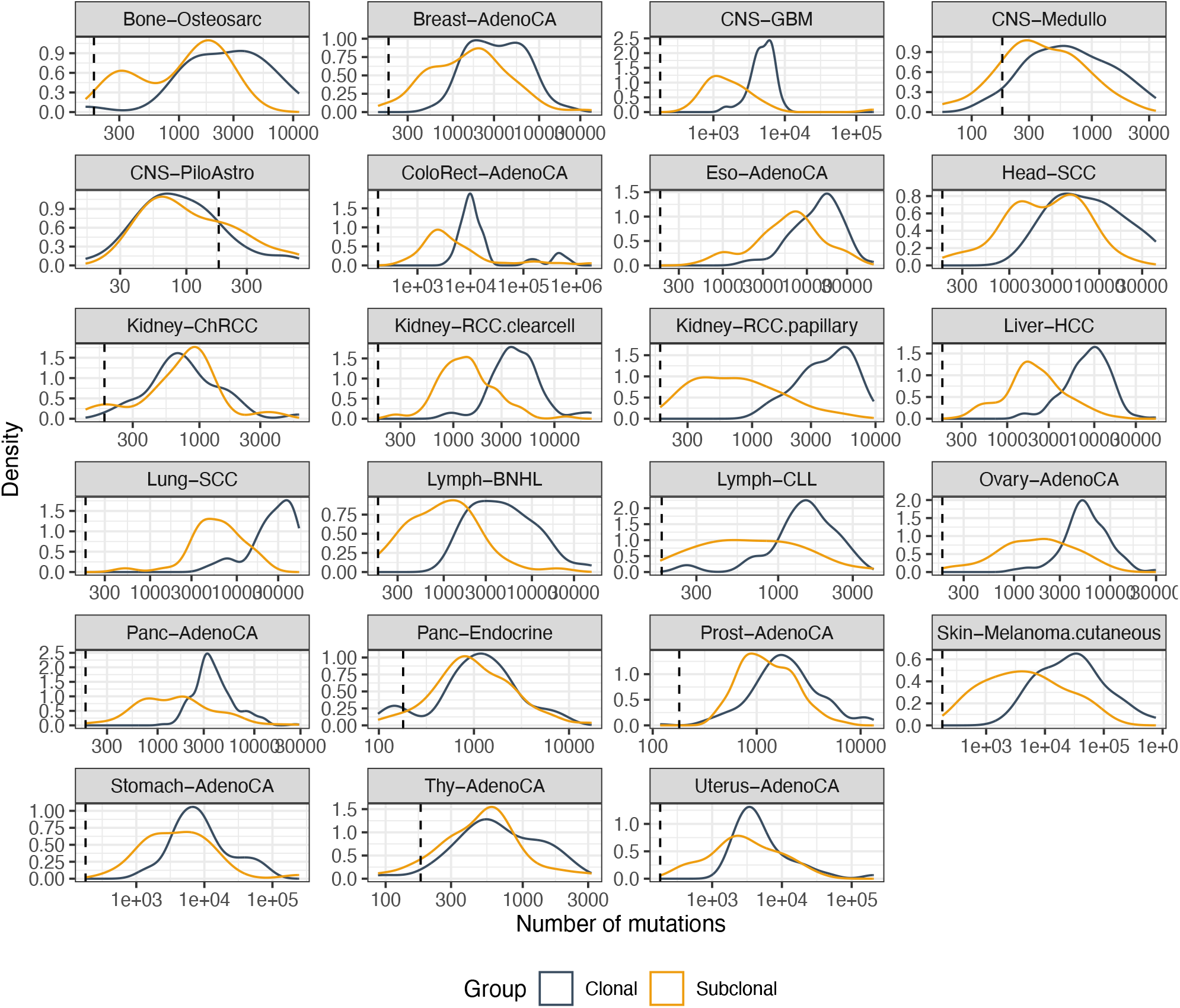
Density plots of the number of mutations in the patient-specific subsamples, for each cancer type.

**Figure S10:**
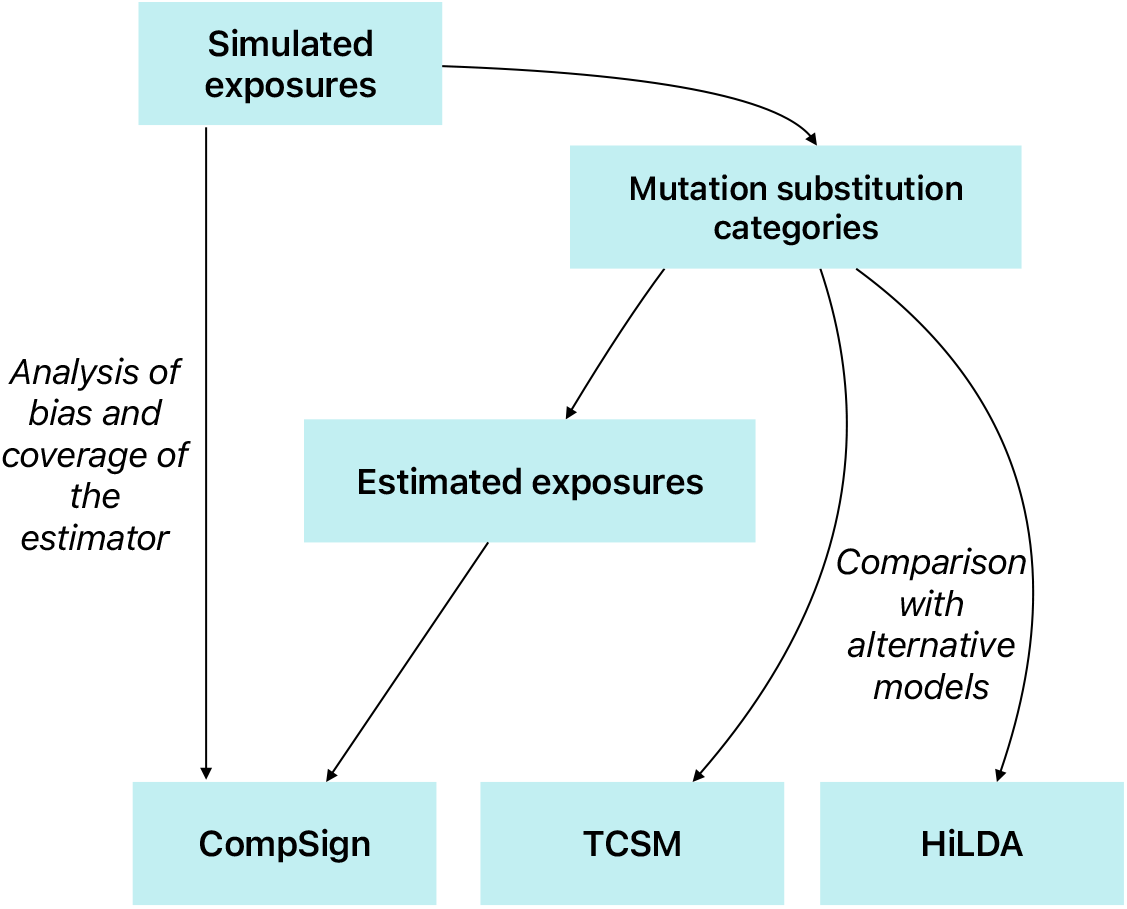
Framework for simulating data which can be used as input for TCSM and HiLDA, as well as `diagREDM` or other `CompSign` models.

**Figure S11:**
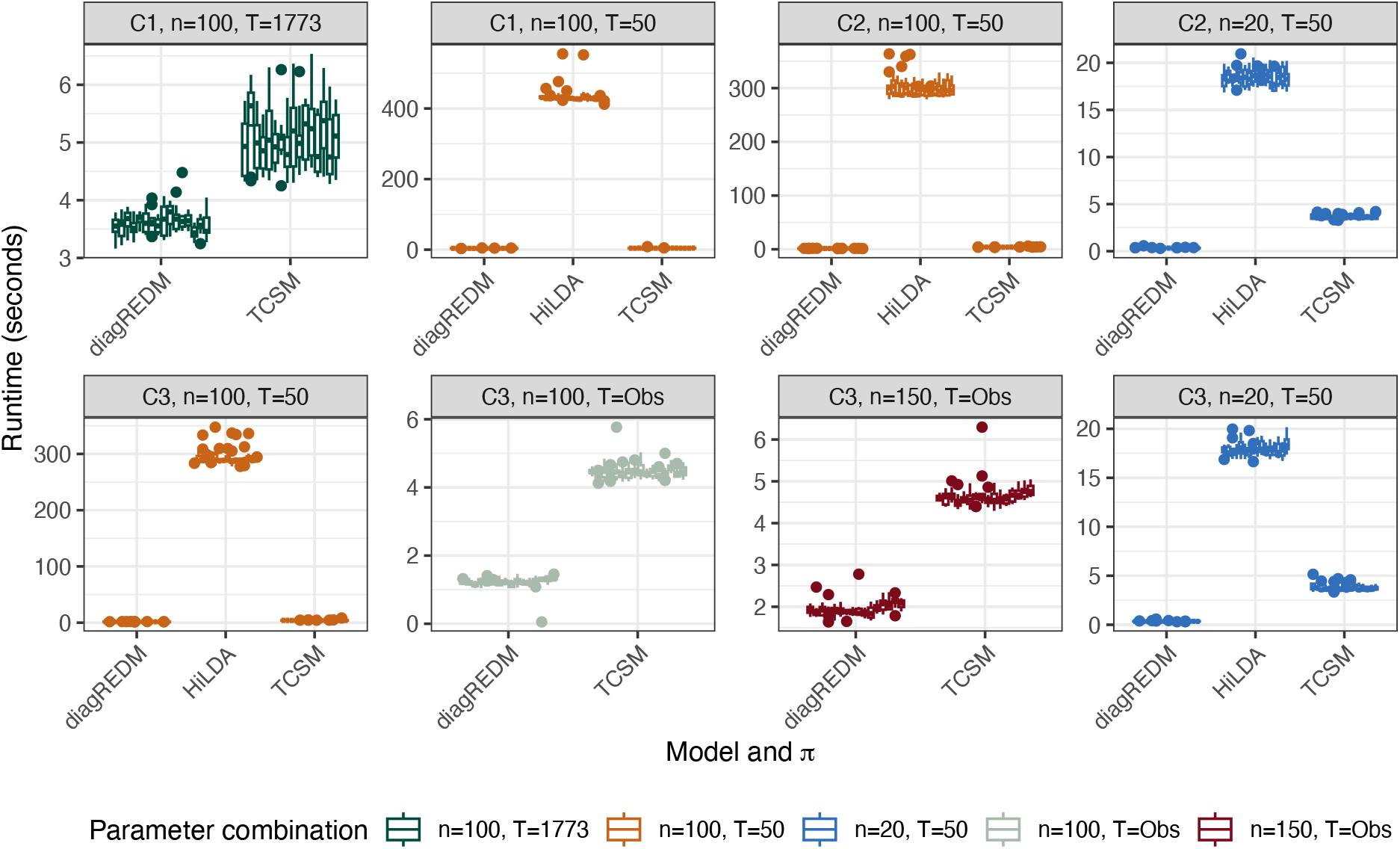
Runtime (in seconds) for each of the models under consideration, in simulations C1-3, as several parameters vary.

**Figure S12:**
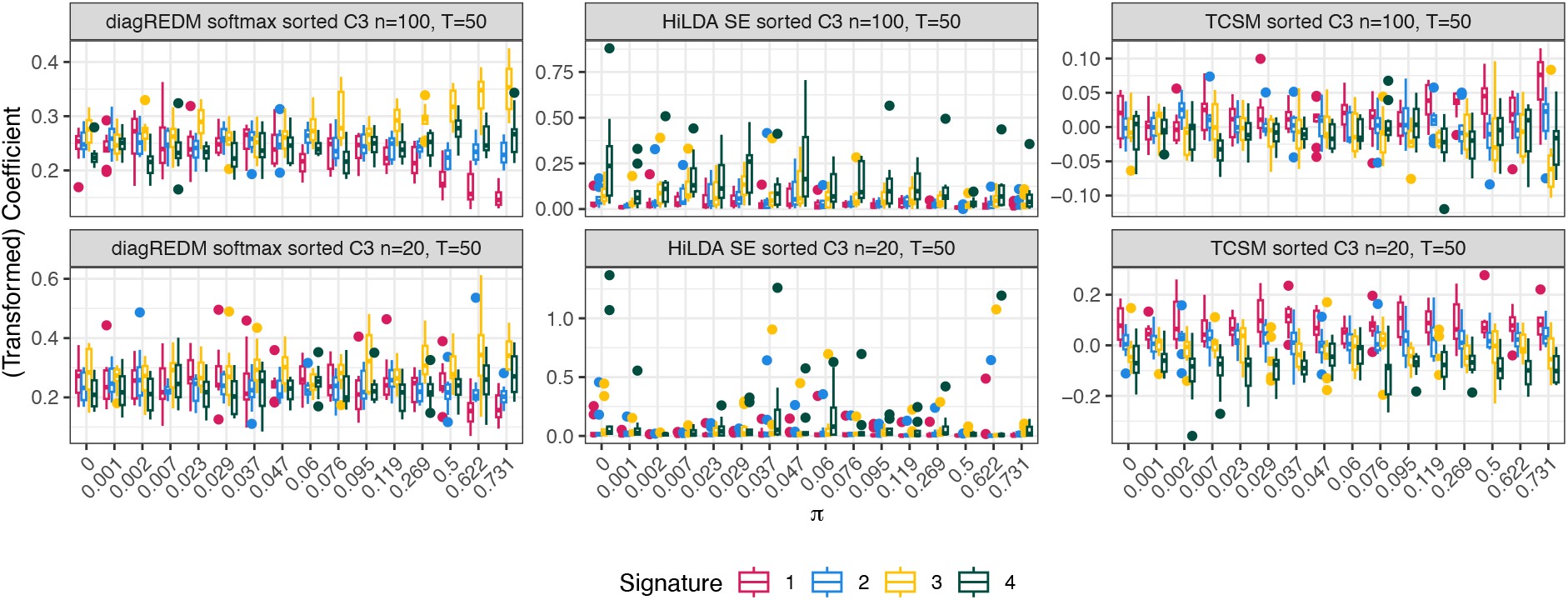
Comparison of coefficients that represent differential abundance for the models under consideration: `diagREDM`, HiLDA, and TCSM, in simulation C3.

**Figure S13:**
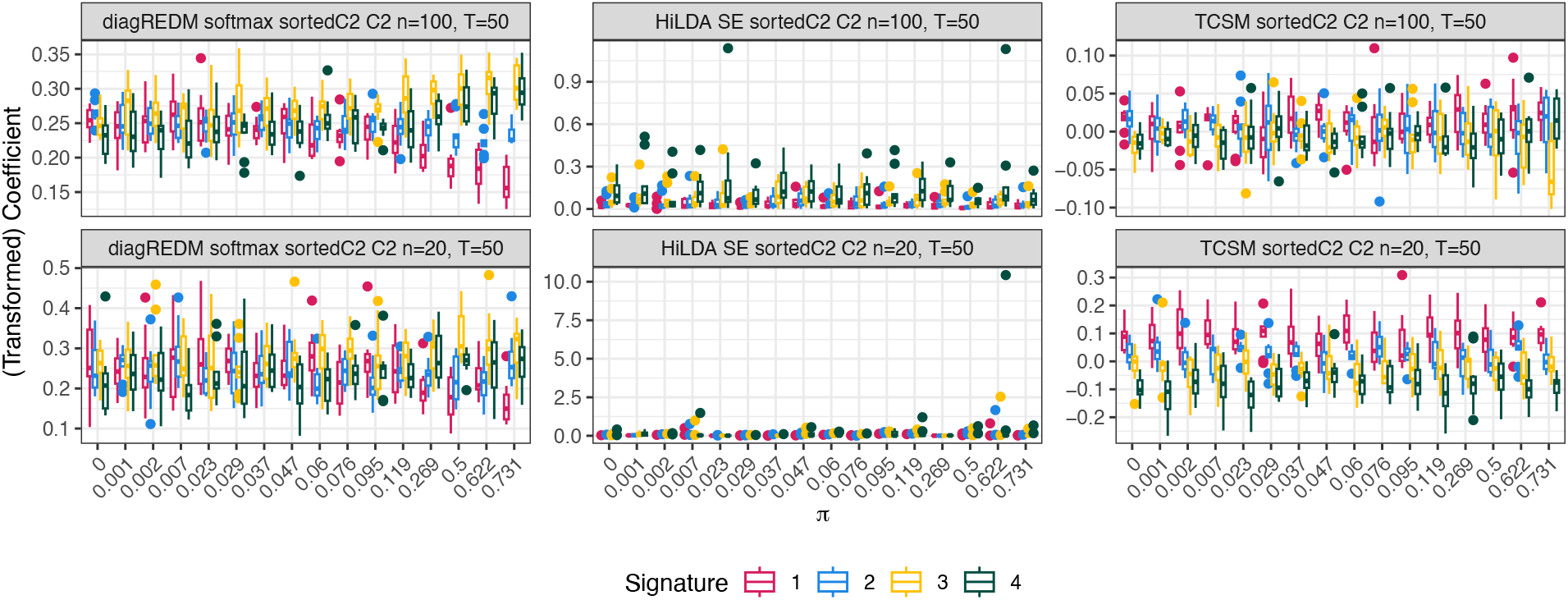
Comparison of coefficients that represent differential abundance for the models under consideration: `diagREDM`, HiLDA, and TCSM, in simulation C2. Note, for `diagREDM`, a poorer signal in C2 than in C3, as data in C3 are simulated with patient-specific intercepts which `diagREDM` models, increasing its statistical power.

**Figure S14:**
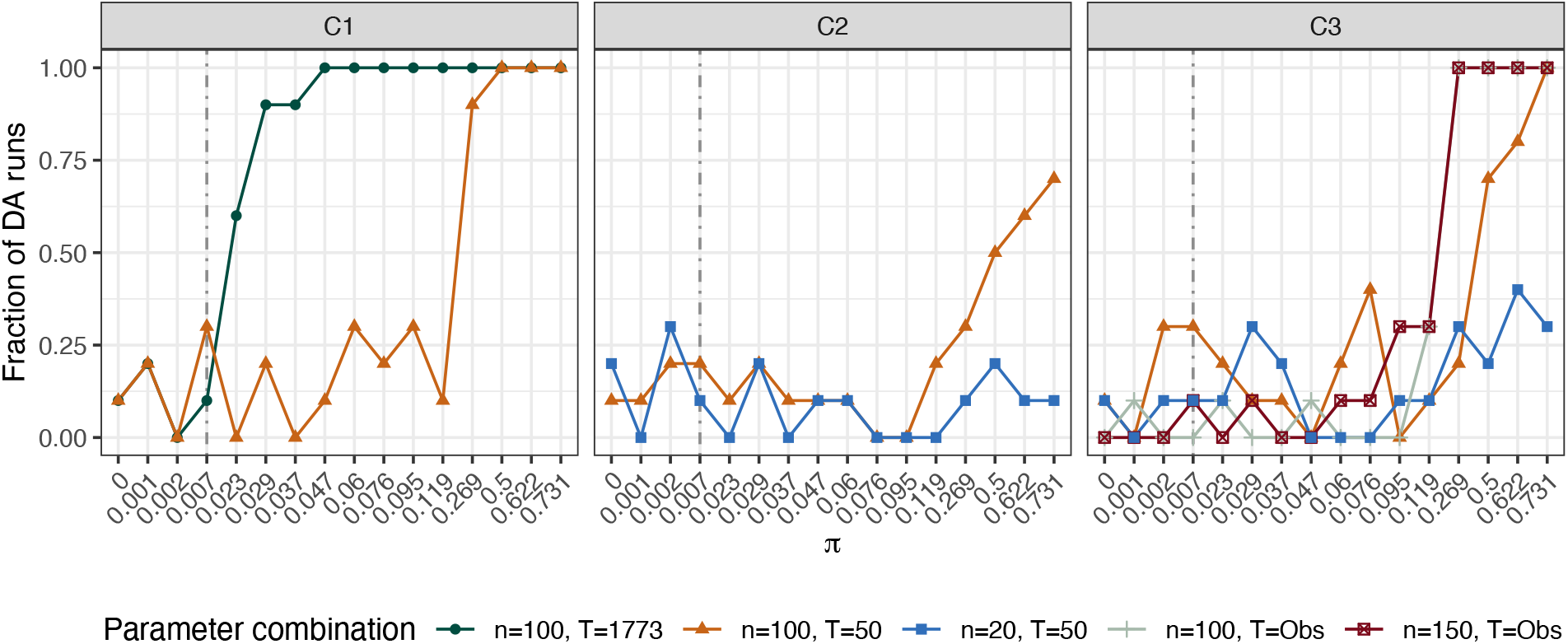
Results of `diagREDM` in simulations C1-C3, with varying parameters *T* and *N*_*s*_. On the *y* axis, the fraction of differentially abundant runs at a significance level of 0.05, as the mixing proportion *π* increases (*x* axis). Both an increase in the number of mutations (first facet) and in the number of patient samples (second facet) lead to higher statistical power.

**Figure S15:**
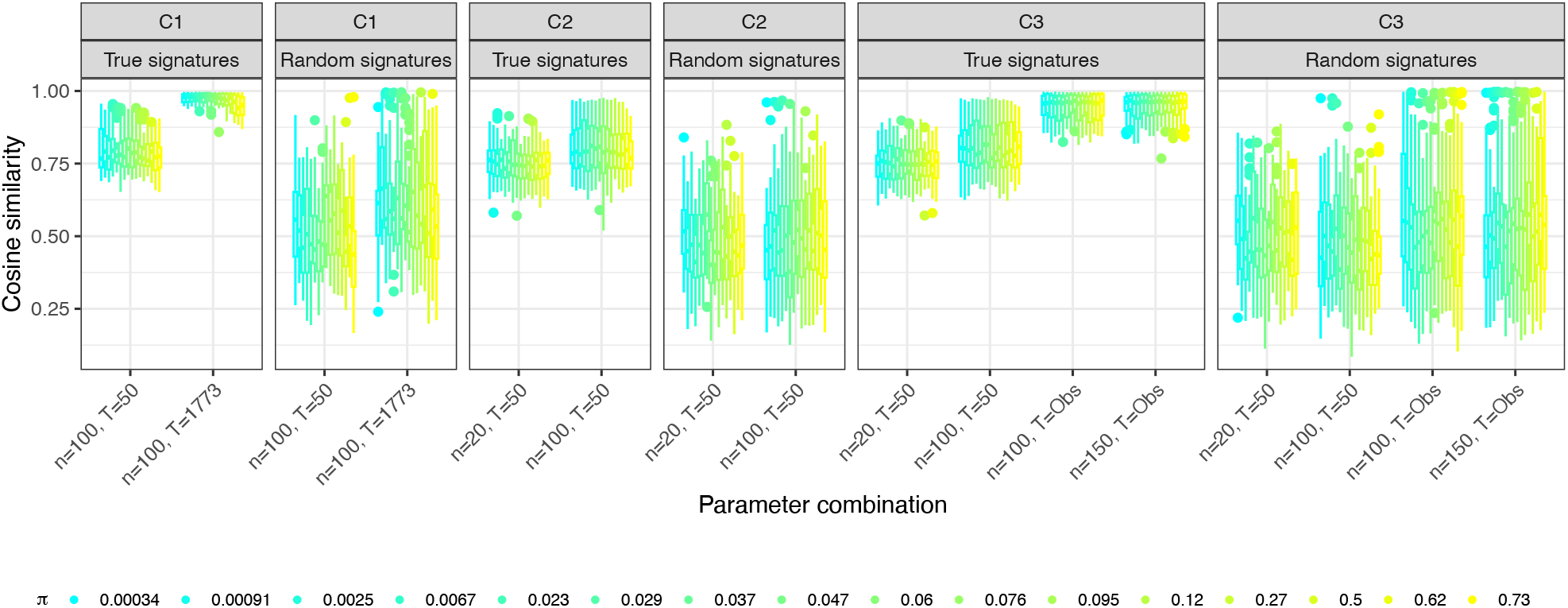
Results of signature extraction TCSM in simulations C1-C3. Higher cosine similarities indicate a better recovery of the ground-truth signatures used to simulate the input data for TCSM. The cosine similarities of the extracted signatures to the ground truth signatures are compared to the cosine similarities of extracted signatures to a set of COSMIC signatures chosen at random. In colour, *π* in increasing abundance. With higher numbers of mutations *T*, signature reconstruction improves in all cases, and so does with a higher number of patient samples.

**Figure S16:**
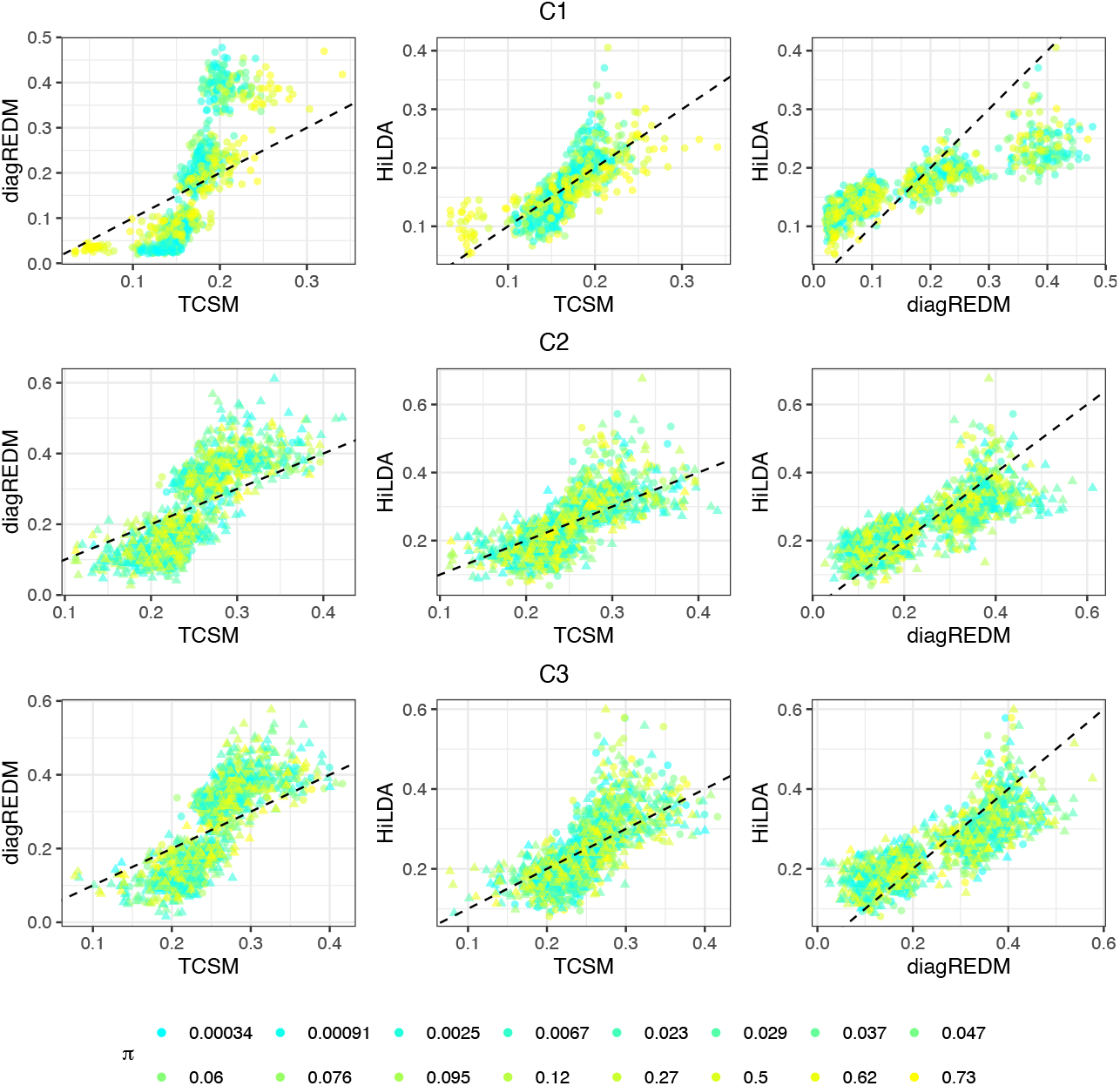
Comparison of abundances in the clonal group of simulated samples, for each of the three models and in all three datasets. Values further away from the identity line indicate higher discrepancies in signature abundance. Note the case in the first and third facets, where sparse signatures are simulated in the highest value of *π* (yellow) but where TCSM and HiLDA overestimate these abundances.

**Figure S17:**
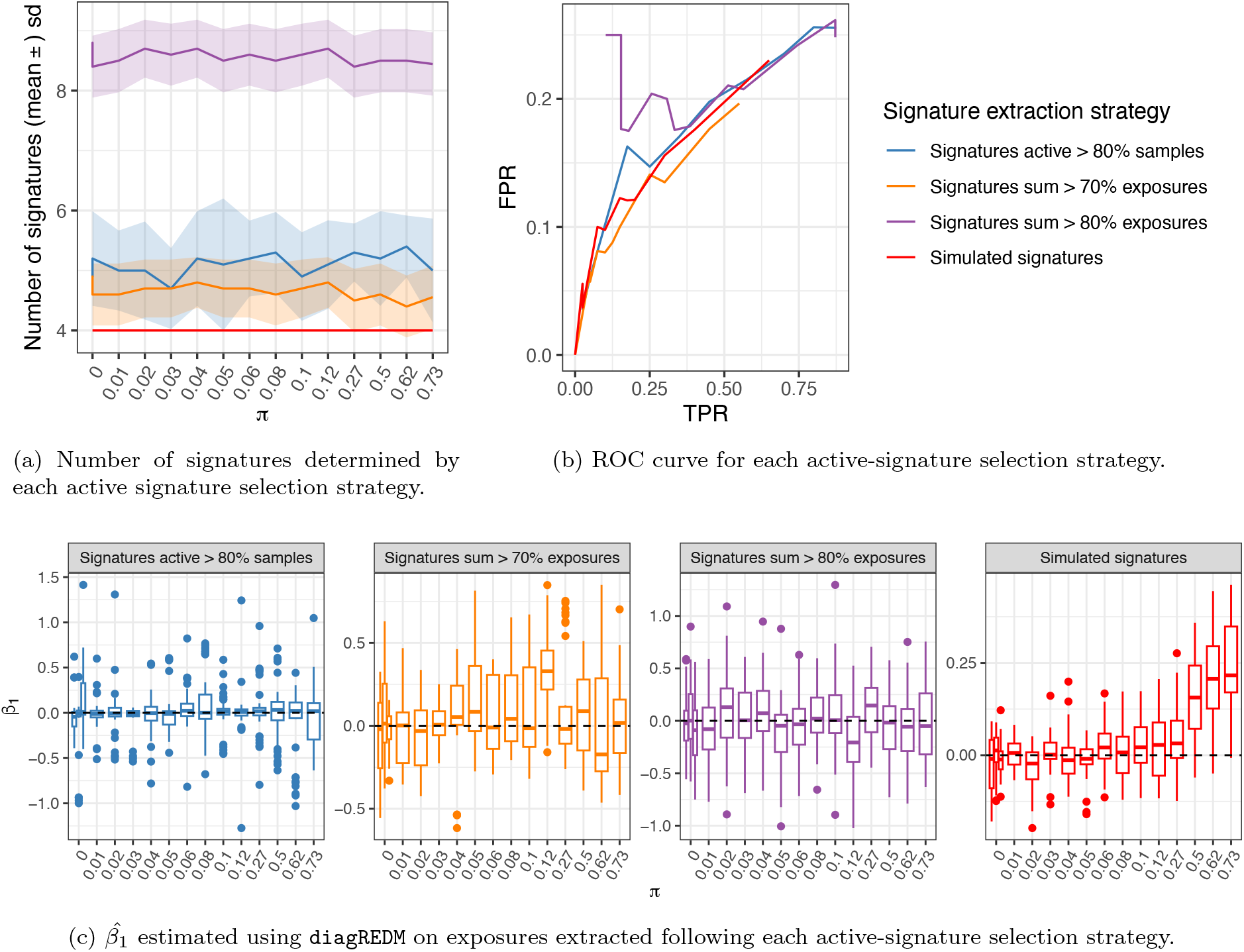
Assessment of the `diagREDM` model when several strategies for selecting the set of active signatures are used.

**Figure S18:**
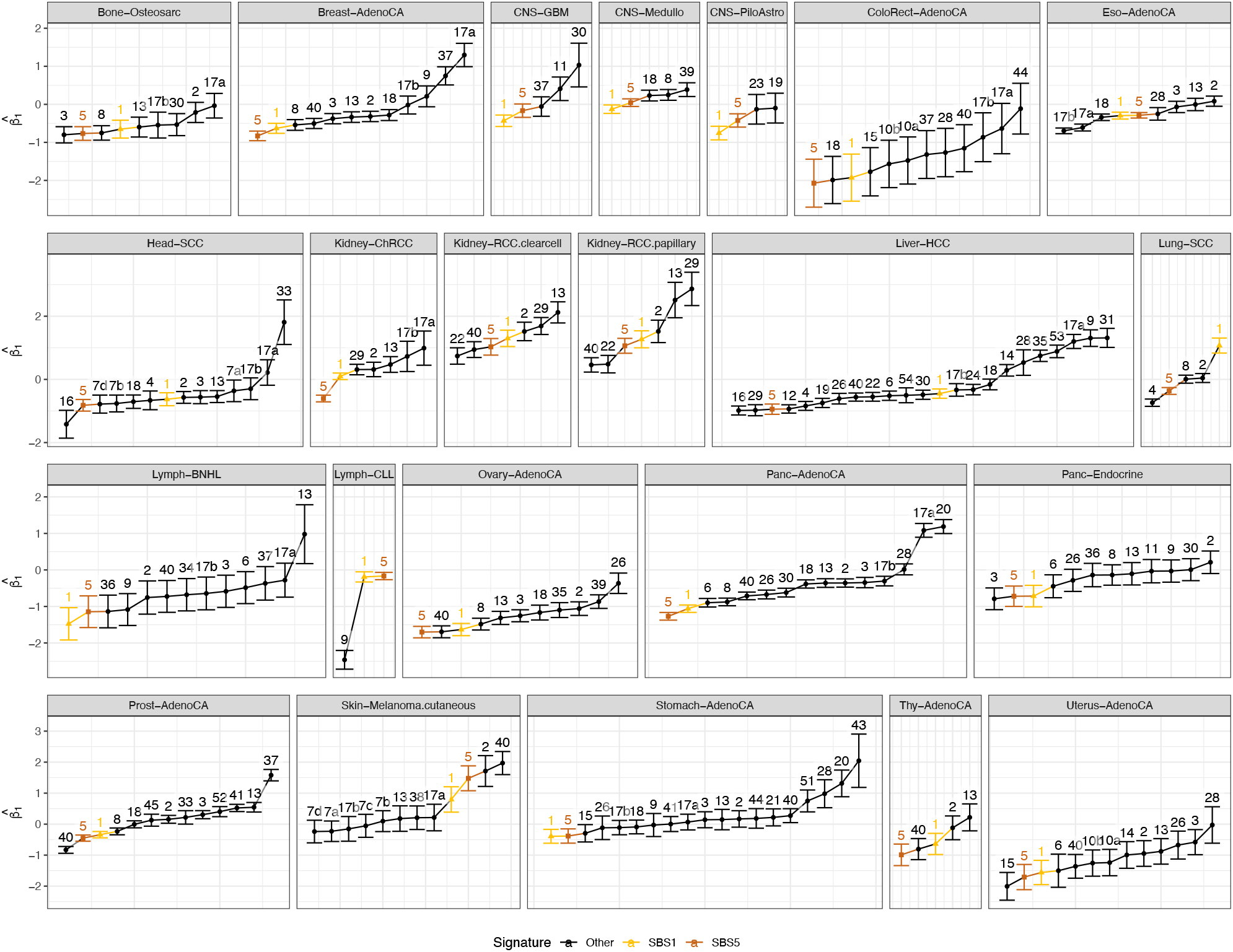
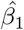 for all cancer types, indicating those in which SBS1 or SBS5 are the signature in the denominator of the ALR transformation that corresponds to the *β*_1_ coefficient – note that these two coefficients are nearly always adjacent, or at least of similar value.

**Figure S19:**
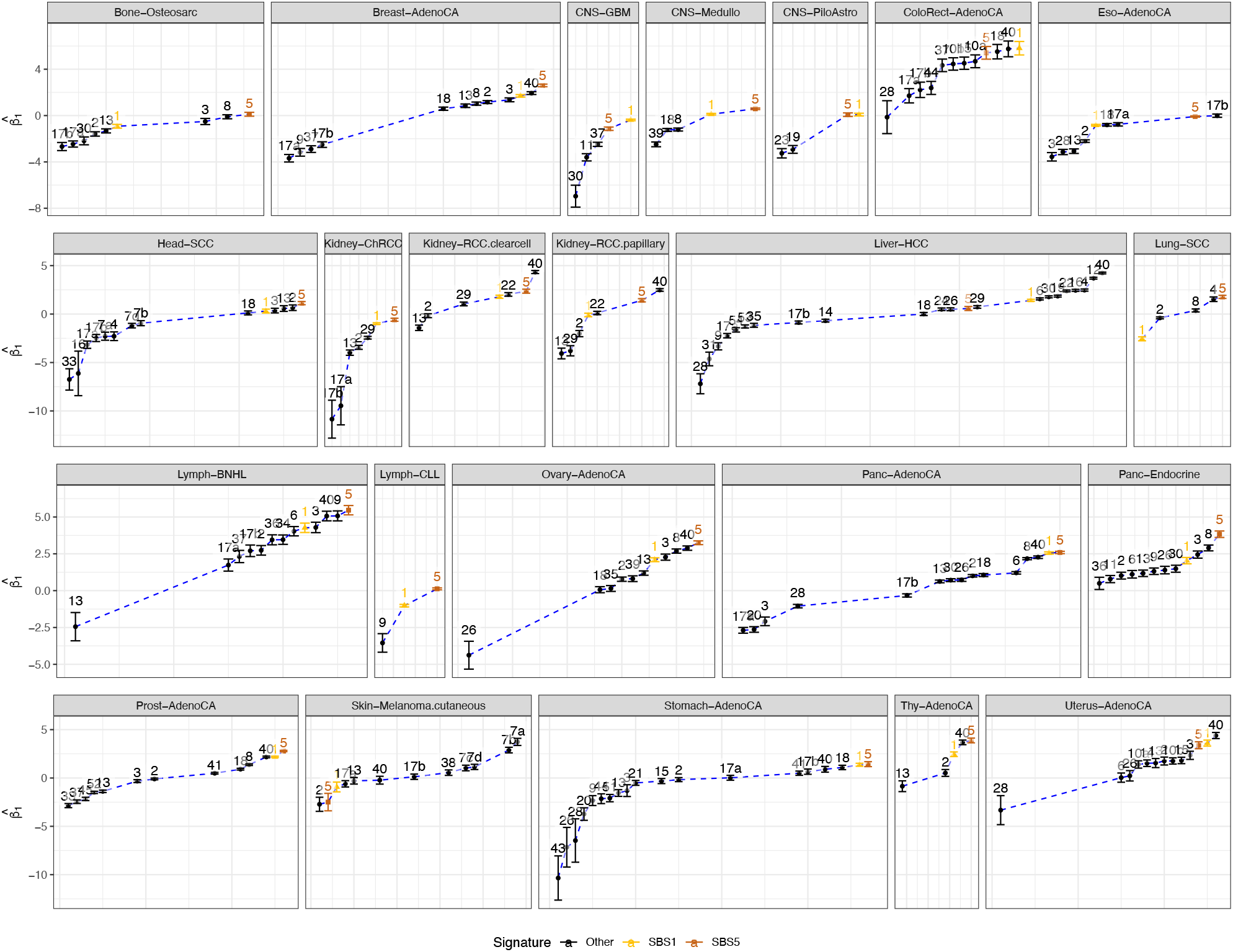
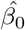 for all cancer types, with the abundance of SBS1 and SBS5 for reference. Lower values indicate lower abundance in the clonal group.

**Figure S20:**
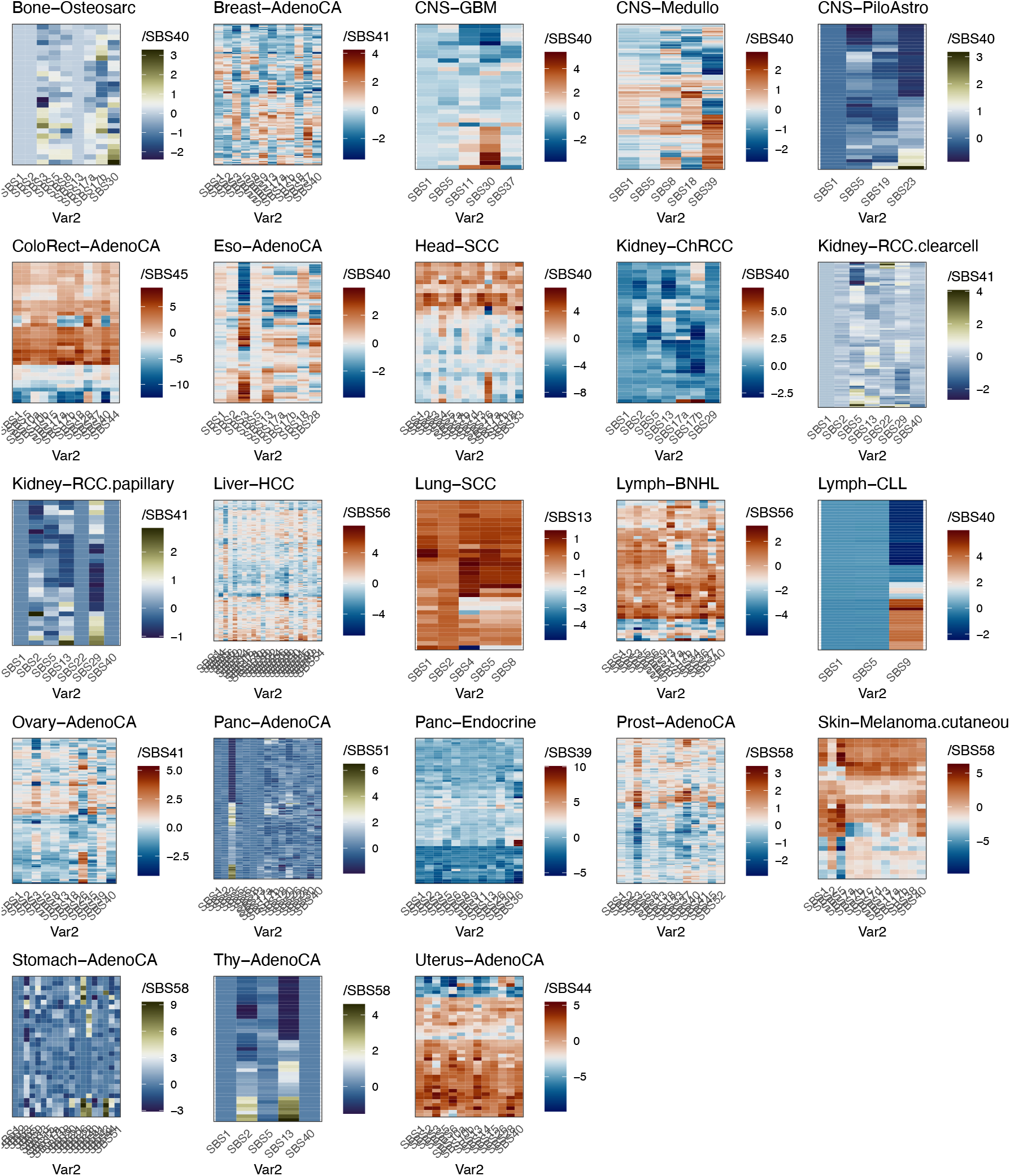
Values of patient random intercepts. In the red colour scheme, values from `fullREDM`, if converged, and values of `diagREDM` otherwise.

**Figure S21:**
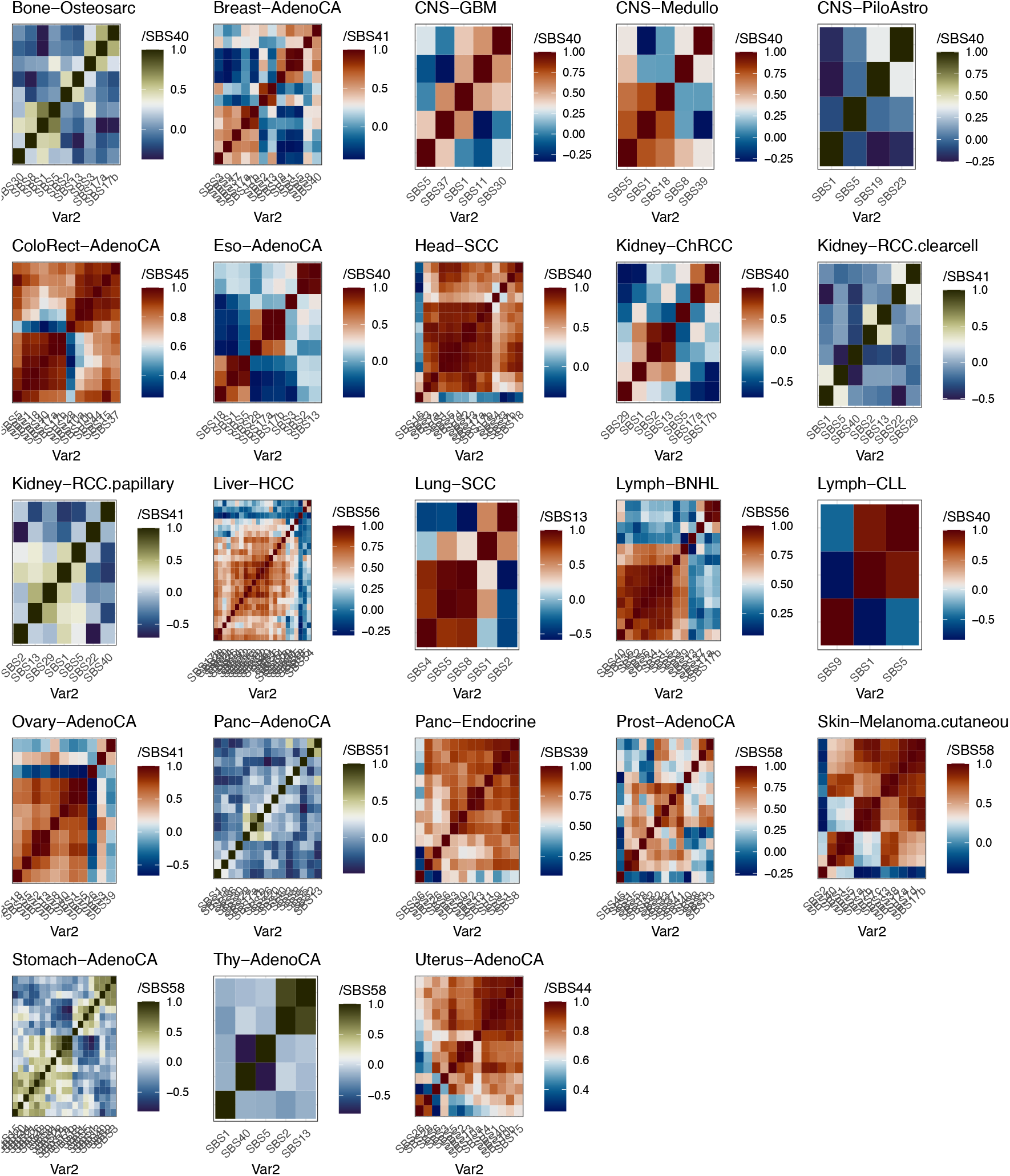
Estimated correlation matrix of patient random intercepts. In the red colour scheme, values from the estimates parameters of `fullREDM`, if converged, and correlation values computed from the random intercept of `diagREDM` otherwise.

**Figure S22:**
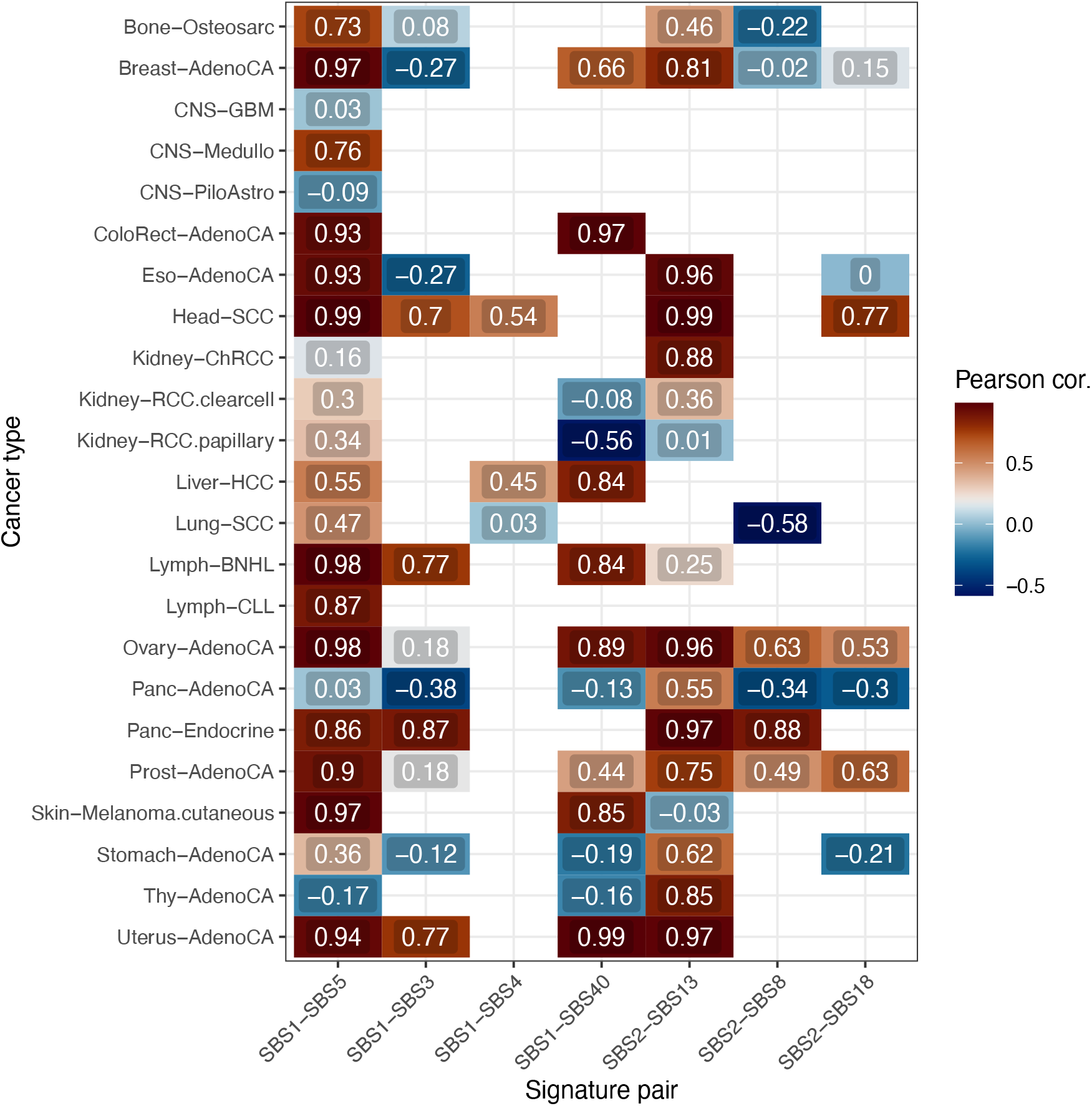
Estimated correlations of patient random intercepts, for selected pairs of signatures, i.e. a small subset of the data in Fig S21.

**Figure S23:**
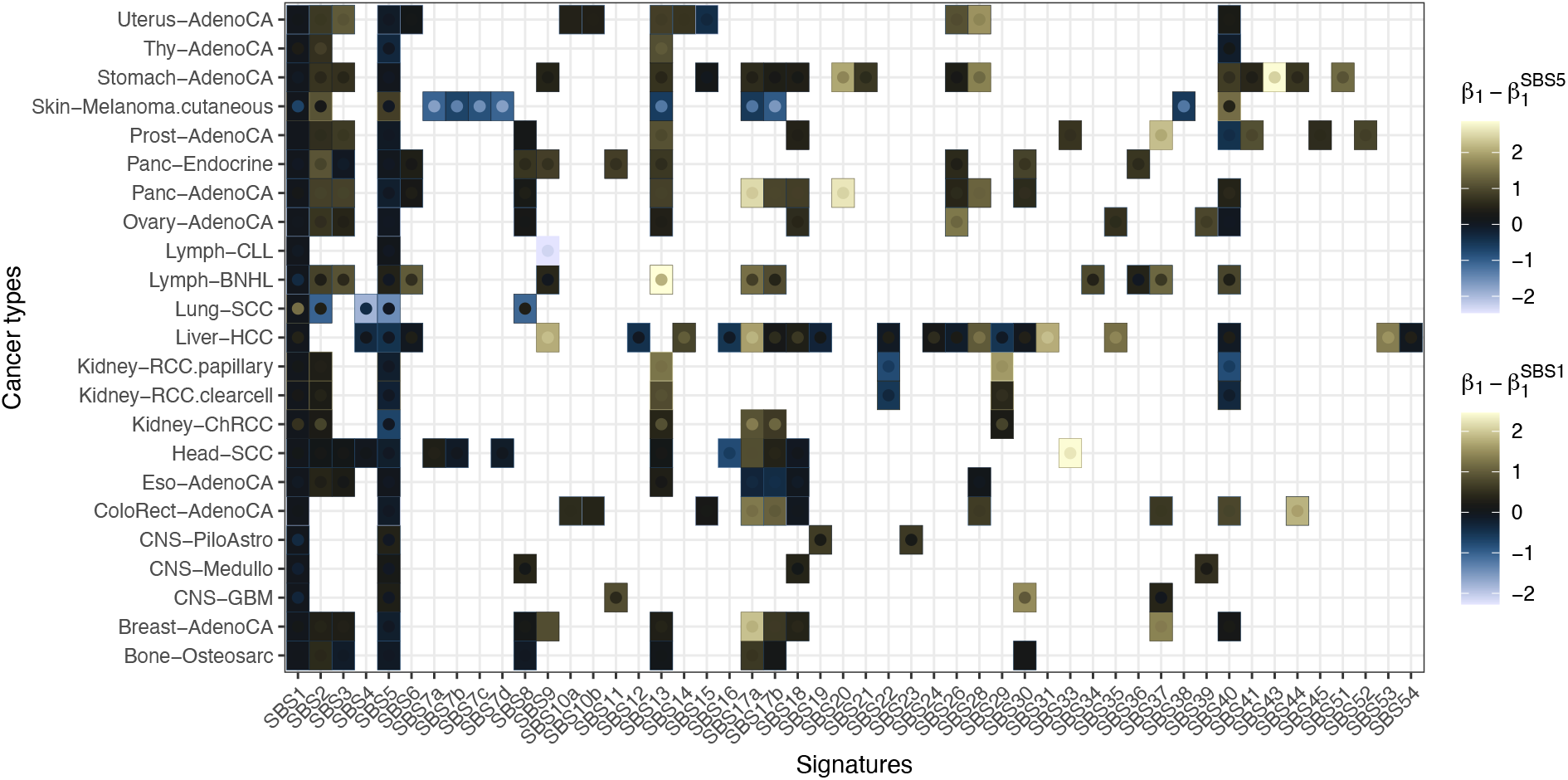
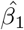 for each signature in each cancer type, subtracted by the 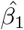 which corresponds to clock signatures SBS1 (rectangles) and SBS5 (points inside rectangles). Yellow colours represent 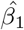 higher than those of clock signatures, and blue lower.

**Figure S24:**
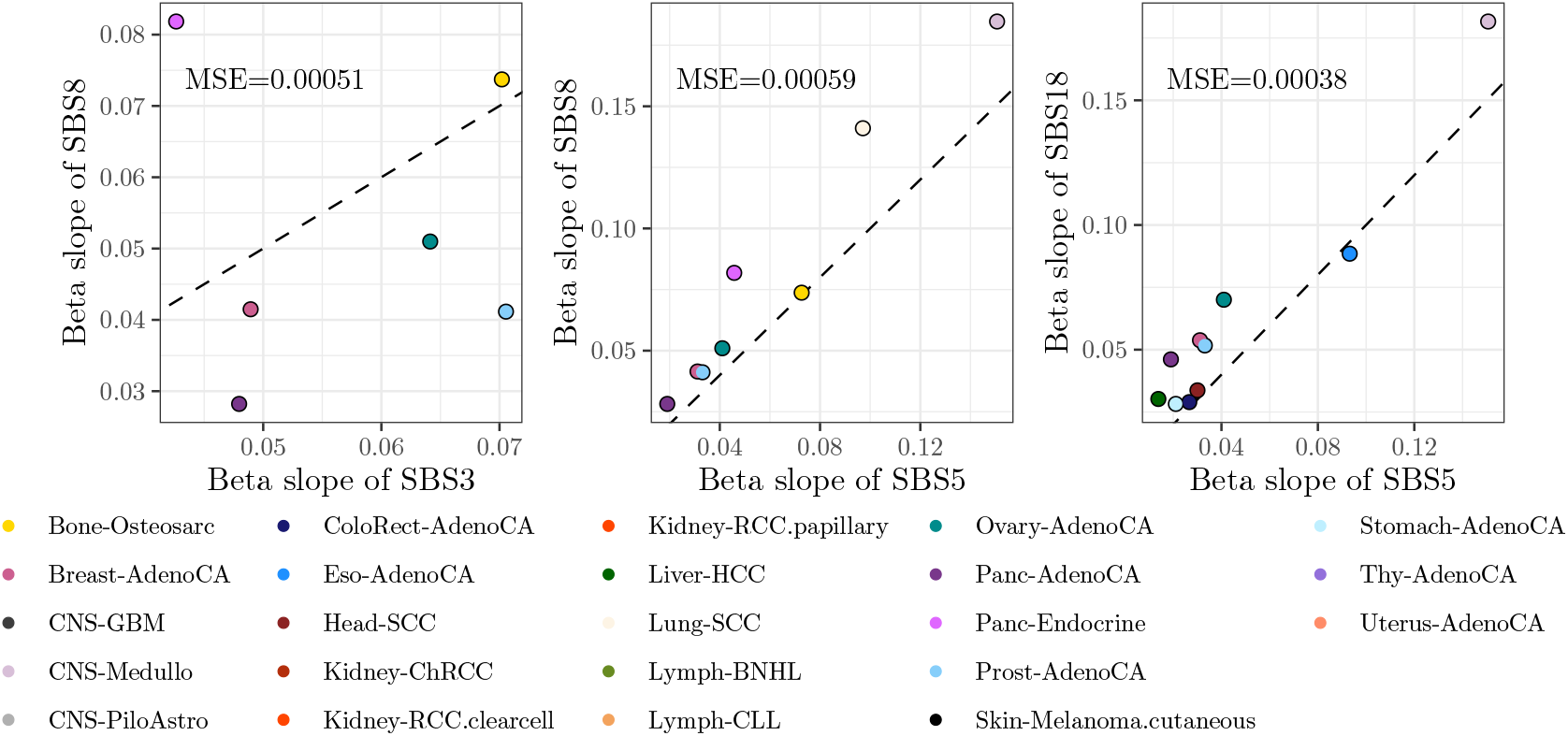
Plots additional to those of Fig 6, with selected pairs of signatures of interest, showing softmax-transformed 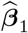 for each cancer type.

**Figure S25:**
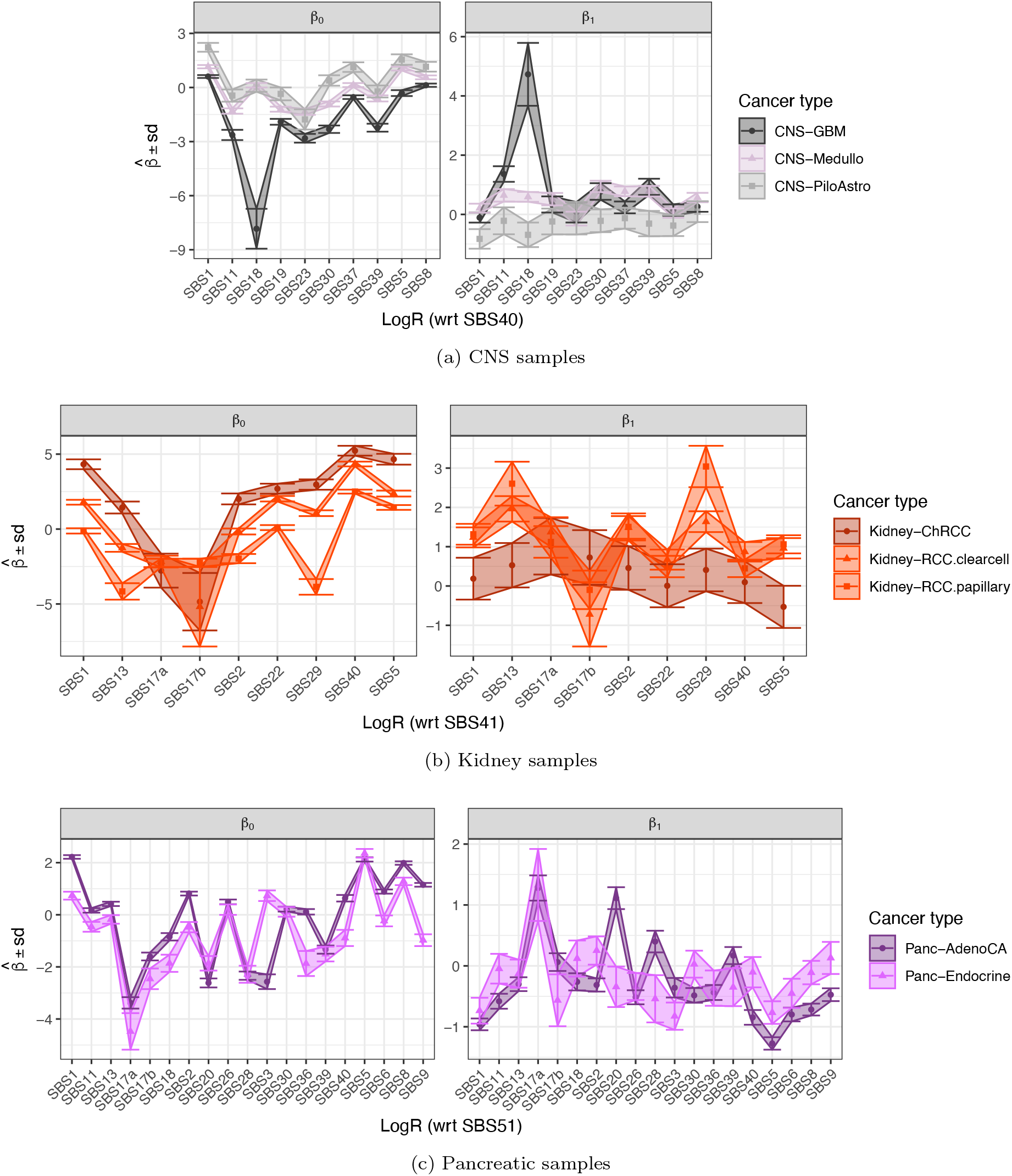
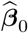 and 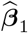 for cancer types of selected tissues. These values are derived once signatures have been re-extracted using all active signatures in each tissue.

**Figure S26:**
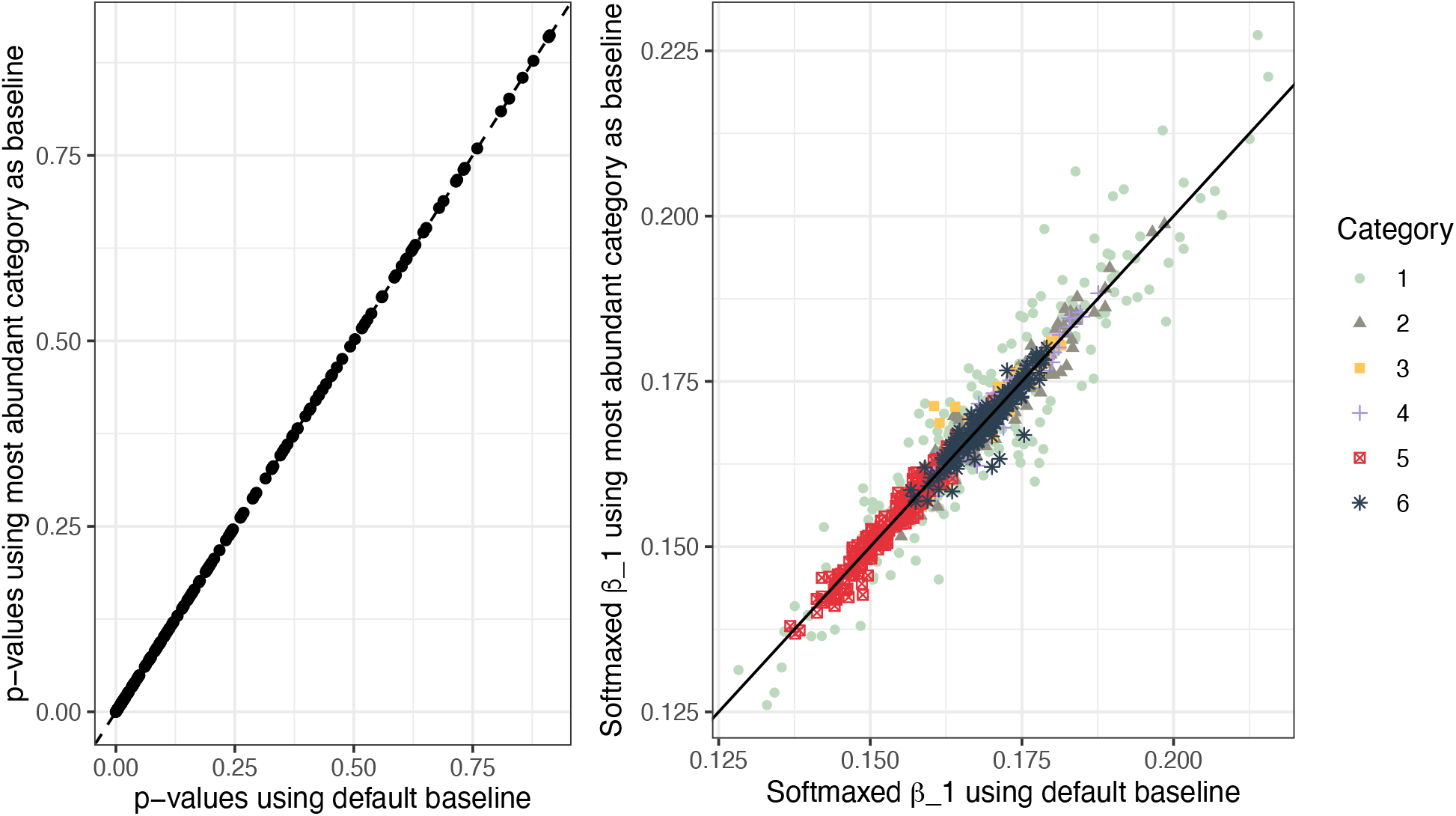
Left: comparison of *p*-values for datasets in which the baseline signature for ALR differs: along the x-axis, the last signature is used as baseline (as default) and this signature is made differentially abundant; along the y-axis, the signature of highest abundance is chosen. The two sets of *p*-values align perfectly. Right: comparison of softmaxed-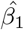 for the same datasets – note that without the softmax the results cannot be compared. There is a very good correlation between the two sets of 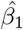 across all signatures (each in a different colour).

## Supplementary tables

**Table S1:**
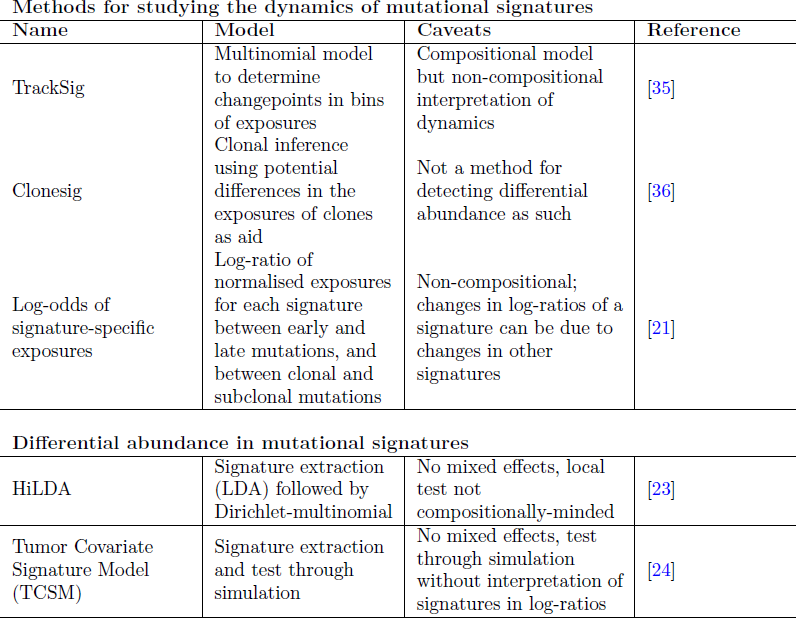
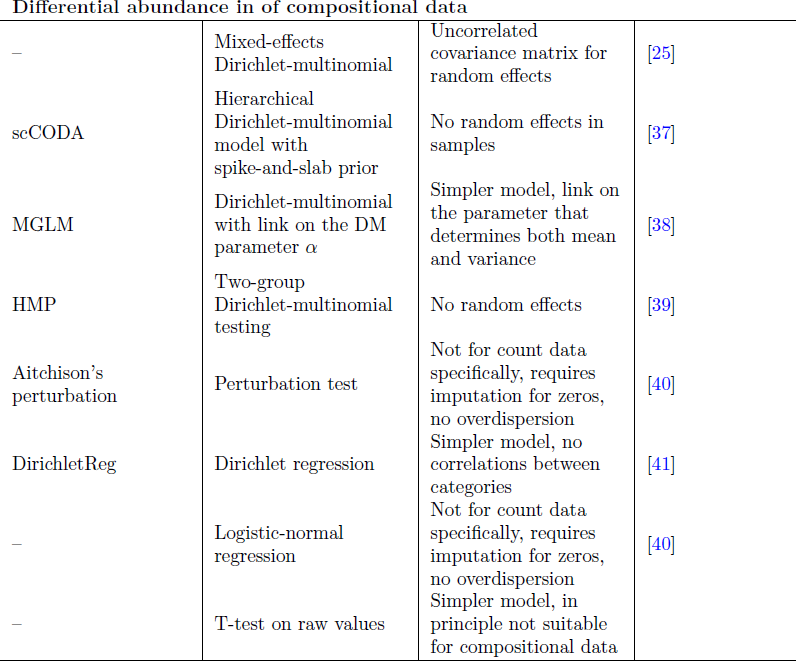
Methods for determining changes in the mutational spectrum, and general regression methods for compositional data.

**Table S2:**
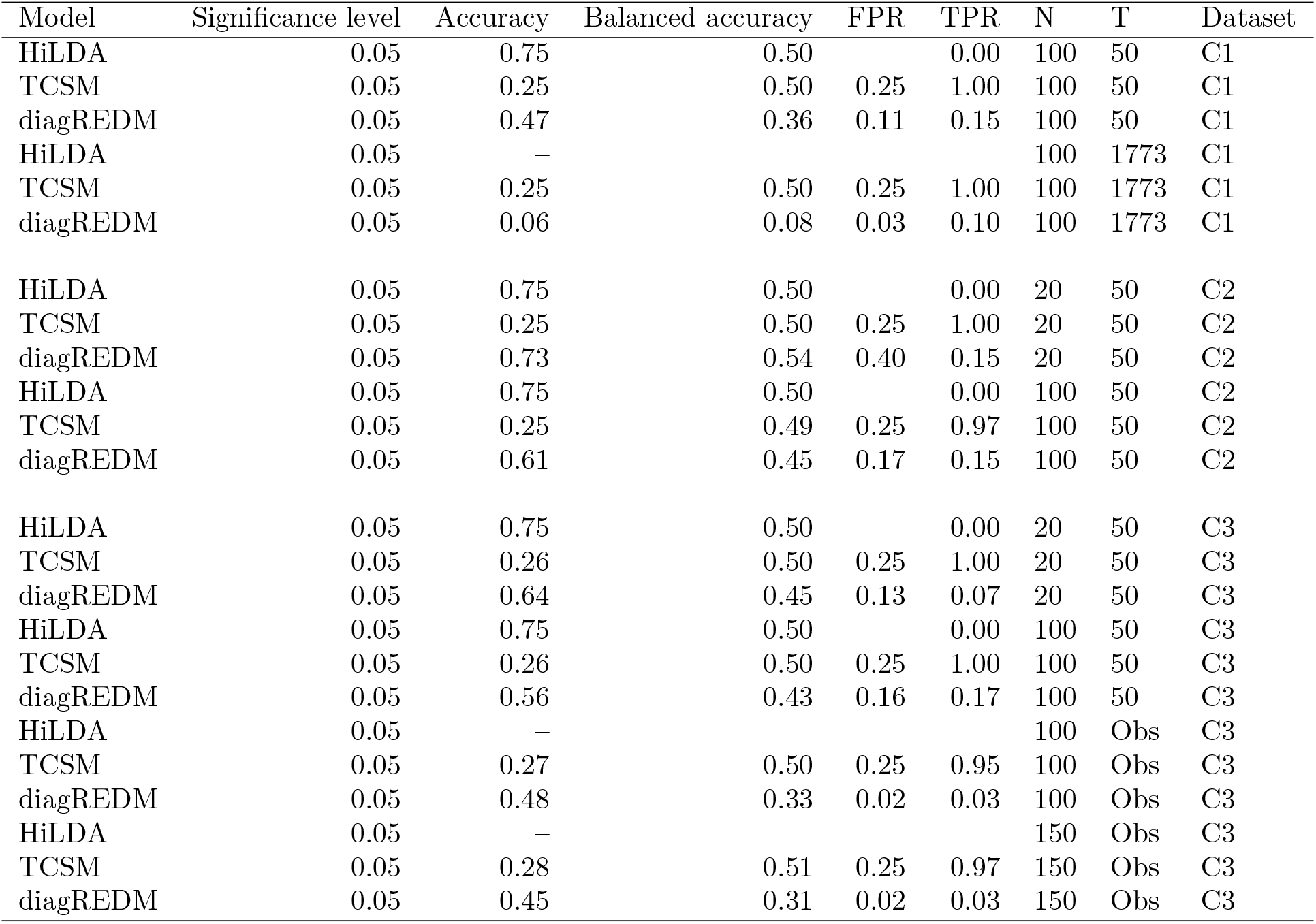
Performance of each model in each dataset, as parameters vary. In three simulations where the number of simulated mutations was relatively high we were not able to get results from HiLDA.

| Model | Significance level | Accuracy | Balanced accuracy | FPR | TPR | N | T | Dataset |
| --- | --- | --- | --- | --- | --- | --- | --- | --- |
| HiLDA | 0.05 | 0.75 | 0.50 |  | 0.00 | 100 | 50 | C1 |
| TCSM | 0.05 | 0.25 | 0.50 | 0.25 | 1.00 | 100 | 50 | C1 |
| diagREDM | 0.05 | 0.47 | 0.36 | 0.11 | 0.15 | 100 | 50 | C1 |
| HiLDA | 0.05 | – |  |  |  | 100 | 1773 | C1 |
| TCSM | 0.05 | 0.25 | 0.50 | 0.25 | 1.00 | 100 | 1773 | C1 |
| diagREDM | 0.05 | 0.06 | 0.08 | 0.03 | 0.10 | 100 | 1773 | C1 |
| HiLDA | 0.05 | 0.75 | 0.50 |  | 0.00 | 20 | 50 | C2 |
| TCSM | 0.05 | 0.25 | 0.50 | 0.25 | 1.00 | 20 | 50 | C2 |
| diagREDM | 0.05 | 0.73 | 0.54 | 0.40 | 0.15 | 20 | 50 | C2 |
| HiLDA | 0.05 | 0.75 | 0.50 |  | 0.00 | 100 | 50 | C2 |
| TCSM | 0.05 | 0.25 | 0.49 | 0.25 | 0.97 | 100 | 50 | C2 |
| diagREDM | 0.05 | 0.61 | 0.45 | 0.17 | 0.15 | 100 | 50 | C2 |
| HiLDA | 0.05 | 0.75 | 0.50 |  | 0.00 | 20 | 50 | C3 |
| TCSM | 0.05 | 0.26 | 0.50 | 0.25 | 1.00 | 20 | 50 | C3 |
| diagREDM | 0.05 | 0.64 | 0.45 | 0.13 | 0.07 | 20 | 50 | C3 |
| HiLDA | 0.05 | 0.75 | 0.50 |  | 0.00 | 100 | 50 | C3 |
| TCSM | 0.05 | 0.26 | 0.50 | 0.25 | 1.00 | 100 | 50 | C3 |
| diagREDM | 0.05 | 0.56 | 0.43 | 0.16 | 0.17 | 100 | 50 | C3 |
| HiLDA | 0.05 | – |  |  |  | 100 | Obs | C3 |
| TCSM | 0.05 | 0.27 | 0.50 | 0.25 | 0.95 | 100 | Obs | C3 |
| diagREDM | 0.05 | 0.48 | 0.33 | 0.02 | 0.03 | 100 | Obs | C3 |
| HiLDA | 0.05 | – |  |  |  | 150 | Obs | C3 |
| TCSM | 0.05 | 0.28 | 0.51 | 0.25 | 0.97 | 150 | Obs | C3 |
| diagREDM | 0.05 | 0.45 | 0.31 | 0.02 | 0.03 | 150 | Obs | C3 |

**Table S3:** Runtime, in seconds, for each of the models, in which diagREDM is shown to be the fastest model.

| Model | Dataset | N | T | Mean (s) | sd (s) | Min (s) | Max (s) |
| --- | --- | --- | --- | --- | --- | --- | --- |
| HiLDA | C1 | 100 | 50 | 431.39 | 15.49 | 412.30 | 554.90 |
| TCSM | C1 | 100 | 50 | 4.31 | 0.51 | 3.67 | 8.60 |
| diagREDM | C1 | 100 | 50 | 3.78 | 0.27 | 2.98 | 4.84 |
| TCSM | C1 | 100 | 1773 | 5.08 | 0.54 | 4.25 | 6.54 |
| diagREDM | C1 | 100 | 1773 | 3.61 | 0.20 | 3.16 | 4.48 |
| HiLDA | C2 | 20 | 50 | 18.42 | 0.87 | 16.86 | 20.95 |
| TCSM | C2 | 20 | 50 | 3.60 | 0.20 | 3.27 | 4.25 |
| diagREDM | C2 | 20 | 50 | 0.33 | 0.03 | 0.27 | 0.58 |
| HiLDA | C2 | 100 | 50 | 298.20 | 15.81 | 279.19 | 363.81 |
| TCSM | C2 | 100 | 50 | 3.95 | 0.26 | 3.59 | 5.93 |
| diagREDM | C2 | 100 | 50 | 1.50 | 0.12 | 1.21 | 1.89 |
| HiLDA | C3 | 20 | 50 | 17.89 | 0.67 | 16.65 | 20.17 |
| TCSM | C3 | 20 | 50 | 3.72 | 0.29 | 3.30 | 5.15 |
| diagREDM | C3 | 20 | 50 | 0.34 | 0.03 | 0.29 | 0.58 |
| HiLDA | C3 | 100 | 50 | 291.20 | 10.14 | 277.81 | 347.62 |
| TCSM | C3 | 100 | 50 | 4.06 | 0.42 | 3.60 | 8.25 |
| diagREDM | C3 | 100 | 50 | 1.54 | 0.12 | 1.33 | 1.97 |
| TCSM | C3 | 100 | Obs | 4.45 | 0.19 | 4.12 | 5.77 |
| diagREDM | C3 | 100 | Obs | 1.22 | 0.13 | 0.05 | 1.47 |
| TCSM | C3 | 150 | Obs | 4.63 | 0.21 | 4.29 | 6.30 |
| diagREDM | C3 | 150 | Obs | 1.95 | 0.16 | 1.64 | 2.78 |

**Table S4:** Fraction of zero exposures, and fraction of zero exposures in the signature with most zeros, in each PCAWG dataset.

|  | Fraction of zeros | Max. fraction of zeros in signatures |
| --- | --- | --- |
| Bone-Osteosarc | 0.12 | 0.37 |
| Breast-AdenoCA | 0.29 | 0.77 |
| CNS-GBM | 0.23 | 0.79 |
| CNS-Medullo | 0.13 | 0.44 |
| CNS-PiloAstro | 0.28 | 0.65 |
| ColoRect-AdenoCA | 0.27 | 0.82 |
| Eso-AdenoCA | 0.10 | 0.42 |
| Head-SCC | 0.27 | 0.81 |
| Kidney-ChRCC | 0.32 | 0.89 |
| Kidney-RCC.clearcell | 0.21 | 0.58 |
| Kidney-RCC.papillary | 0.22 | 0.67 |
| Liver-HCC | 0.39 | 0.93 |
| Lung-SCC | 0.03 | 0.13 |
| Lymph-BNHL | 0.22 | 0.88 |
| Lymph-CLL | 0.15 | 0.58 |
| Ovary-AdenoCA | 0.18 | 0.76 |
| Panc-AdenoCA | 0.19 | 0.61 |
| Panc-Endocrine | 0.24 | 0.59 |
| Prost-AdenoCA | 0.26 | 0.76 |
| Skin-Melanoma.cutaneous | 0.26 | 0.62 |
| Stomach-AdenoCA | 0.30 | 0.88 |
| Thy-AdenoCA | 0.22 | 0.52 |
| Uterus-AdenoCA | 0.31 | 0.79 |

**Table S5:** Active signatures in each of the cancer types, according to the PCAWG analyses. With asterisks, signatures which are considered exogenous for these analysis: those include signatures known to represent exogenous mutational processes (based on the COSMIC annotation of signatures), as well as SBS1 and SBS5. The last signature is used as baseline. Should this signature be exogenous, the last signature without an asterisk is used as baseline.

| Cancer type | Active signatures |
| --- | --- |
| Bone-Osteosarc | SBS1*, SBS2, SBS3, SBS5*, SBS8, SBS13, SBS17a, SBS17b, SBS30, SBS40 |
| Breast-AdenoCA | SBS1*, SBS2, SBS3, SBS5*, SBS8, SBS9, SBS13, SBS17a, SBS17b, SBS18, SBS37, SBS40, SBS41 |
| CNS-GBM | SBS1*, SBS5*, SBS11*, SBS30, SBS37, SBS40 |
| CNS-Medullo | SBS1*, SBS5*, SBS8, SBS18, SBS39, SBS40 |
| CNS-PiloAstro | SBS1*, SBS5*, SBS19, SBS23, SBS40 |
| ColoRect-AdenoCA | SBS1*, SBS5*, SBS10a, SBS10b, SBS15, SBS17a, SBS17b, SBS18, SBS28, SBS37, SBS40, SBS44, SBS45* |
| Eso-AdenoCA | SBS1*, SBS2, SBS3, SBS5*, SBS13, SBS17a, SBS17b, SBS18, SBS28, SBS40 |
| Head-SCC | SBS1*, SBS2, SBS3, SBS4*, SBS5*, SBS7a*, SBS7b*, SBS7d*, SBS13, SBS16, SBS17a, SBS17b, SBS18, SBS33, SBS40 |
| Kidney-ChRCC | SBS1*, SBS2, SBS5*, SBS13, SBS17a, SBS17b, SBS29*, SBS40 |
| Kidney-RCC.clearcell | SBS1*, SBS2, SBS5*, SBS13, SBS22, SBS29*, SBS40, SBS41 |
| Kidney-RCC.papillary | SBS1*, SBS2, SBS5*, SBS13, SBS22, SBS29*, SBS40, SBS41 |
| Liver-HCC | SBS1*, SBS4*, SBS5*, SBS6, SBS9, SBS12, SBS14, SBS16, SBS17a, SBS17b, SBS18, SBS19, SBS22, SBS24, SBS26, SBS28, SBS29*, SBS30, SBS31*, SBS35*, SBS40, SBS53, SBS54*, SBS56* |
| Lung-SCC | SBS1*, SBS2, SBS4*, SBS5*, SBS8, SBS13 |
| Lymph-BNHL | SBS1*, SBS2, SBS3, SBS5*, SBS6, SBS9, SBS13, SBS17a, SBS17b, SBS34, SBS36, SBS37, SBS40, SBS56* |
| Lymph-CLL | SBS1*, SBS5*, SBS9, SBS40 |
| Ovary-AdenoCA | SBS1*, SBS2, SBS3, SBS5*, SBS8, SBS13, SBS18, SBS26, SBS35*, SBS39, SBS40, SBS41 |
| Panc-AdenoCA | SBS1*, SBS2, SBS3, SBS5*, SBS6, SBS8, SBS13, SBS17a, SBS17b, SBS18, SBS20, SBS26, SBS28, SBS30, SBS40, SBS51* |
| Panc-Endocrine | SBS1*, SBS2, SBS3, SBS5*, SBS6, SBS8, SBS9, SBS11*, SBS13, SBS26, SBS30, SBS36, SBS39 |
| Prost-AdenoCA | SBS1*, SBS2, SBS3, SBS5*, SBS8, SBS13, SBS18, SBS33, SBS37, SBS40, SBS41, SBS45*, SBS52*, SBS58* |
| Skin-Melanoma.cutaneous | SBS1*, SBS2, SBS5*, SBS7a*, SBS7b*, SBS7c*, SBS7d*, SBS13, SBS17a, SBS17b, SBS38, SBS40, SBS58* |
| Stomach-AdenoCA | SBS1*, SBS2, SBS3, SBS5*, SBS9, SBS13, SBS15, SBS17a, SBS17b, SBS18, SBS20, SBS21, SBS26, SBS28, SBS40, SBS41, SBS43*, SBS44, SBS51*, SBS58* |
| Thy-AdenoCA | SBS1*, SBS2, SBS5*, SBS13, SBS40, SBS58* |
| Uterus-AdenoCA | SBS1*, SBS2, SBS3, SBS5*, SBS6, SBS10a, SBS10b, SBS13, SBS14, SBS15, SBS26, SBS28, SBS40, SBS44 |

**Table S6:** Summary statistics of the number of mutations in each cancer type and group, across patients.

| Cancer type | Group | Min | Max | Mean | Median |
| --- | --- | --- | --- | --- | --- |
| Bone-Osteosarc | Clonal | 152 | 11274 | 3393.78 | 2986 |
| Bone-Osteosarc | Subclonal | 227 | 3635 | 1370.59 | 1334 |
| Breast-AdenoCA | Clonal | 557 | 24655 | 4351.21 | 3170 |
| Breast-AdenoCA | Subclonal | 136 | 40540 | 2663.91 | 1643.50 |
| CNS-GBM | Clonal | 1477 | 9482 | 5119.32 | 5083.50 |
| CNS-GBM | Subclonal | 474 | 175464 | 6996 | 1479 |
| CNS-Medullo | Clonal | 121 | 3413 | 893 | 646.50 |
| CNS-Medullo | Subclonal | 57 | 2151 | 508.78 | 368 |
| CNS-PiloAstro | Clonal | 16 | 645 | 125.74 | 79.50 |
| CNS-PiloAstro | Subclonal | 31 | 776 | 160.95 | 87 |
| ColoRect-AdenoCA | Clonal | 4105 | 719890 | 73922.03 | 10462 |
| ColoRect-AdenoCA | Subclonal | 595 | 1938252 | 80149.54 | 2993 |
| Eso-AdenoCA | Clonal | 1839 | 62315 | 16195.14 | 15277 |
| Eso-AdenoCA | Subclonal | 551 | 36173 | 8683.60 | 6798 |
| Head-SCC | Clonal | 1496 | 42607 | 11117.69 | 6871.50 |
| Head-SCC | Subclonal | 211 | 18413 | 3773.16 | 3078 |
| Kidney-ChRCC | Clonal | 190 | 6044 | 988.26 | 709.50 |
| Kidney-ChRCC | Subclonal | 128 | 4078 | 896.79 | 836 |
| Kidney-RCC.clearcell | Clonal | 823 | 23326 | 5027.29 | 4014.50 |
| Kidney-RCC.clearcell | Subclonal | 249 | 12993 | 1704.60 | 1371 |
| Kidney-RCC.papillary | Clonal | 1268 | 9675 | 4435.30 | 4292 |
| Kidney-RCC.papillary | Subclonal | 251 | 5075 | 1046.67 | 756 |
| Liver-HCC | Clonal | 1420 | 50963 | 9800.13 | 9089 |
| Liver-HCC | Subclonal | 268 | 20069 | 2880.89 | 1968 |
| Lung-SCC | Clonal | 4811 | 52291 | 29414.12 | 27600 |
| Lung-SCC | Subclonal | 515 | 20221 | 7266.18 | 6332.50 |
| Lymph-BNHL | Clonal | 1281 | 47499 | 7076.04 | 4521 |
| Lymph-BNHL | Subclonal | 181 | 19374 | 1658.04 | 991 |
| Lymph-CLL | Clonal | 249 | 3231 | 1596.25 | 1529 |
| Lymph-CLL | Subclonal | 176 | 4083 | 883.57 | 649 |
| Ovary-AdenoCA | Clonal | 1003 | 29855 | 6480.45 | 5455 |
| Ovary-AdenoCA | Subclonal | 193 | 10605 | 2436.48 | 1802 |
| Panc-AdenoCA | Clonal | 1145 | 26491 | 4374.25 | 3472 |
| Panc-AdenoCA | Subclonal | 180 | 29039 | 2377.83 | 1641 |
| Panc-Endocrine | Clonal | 98 | 8483 | 1692.56 | 1178.50 |
| Panc-Endocrine | Subclonal | 104 | 17275 | 1636.61 | 891 |
| Prost-AdenoCA | Clonal | 118 | 13366 | 2495.90 | 1761 |
| Prost-AdenoCA | Subclonal | 382 | 7083 | 1527.89 | 1164 |
| Skin-Melanoma.cutaneous | Clonal | 2841 | 767734 | 75208.07 | 30374 |
| Skin-Melanoma.cutaneous | Subclonal | 367 | 82458 | 12788.90 | 4105 |
| Stomach-AdenoCA | Clonal | 1345 | 75901 | 14993.20 | 8200 |
| Stomach-AdenoCA | Subclonal | 429 | 246341 | 13698.73 | 4192 |
| Thy-AdenoCA | Clonal | 95 | 2322 | 828.51 | 614 |
| Thy-AdenoCA | Subclonal | 88 | 3177 | 641.27 | 533 |
| Uterus-AdenoCA | Clonal | 1689 | 207015 | 12120.40 | 3985.50 |
| Uterus-AdenoCA | Subclonal | 338 | 29388 | 5219.82 | 2652.50 |

## Supplementary methods

### S1 Signature extraction

Consider that we are given *N* samples or sets of mutations and have defined *F* features summarizing genomic changes (e.g., *F* = 96 in COSMIC signatures), indexed by *j ∈* [*N*] and *f ∈* [*F*], where [*x*] denote the sequence 1,…, *x*. The data set can then be written as a count matrix **V** *∈* ℕ^*F×N*^, where each row represents one class of mutations (e.g. one row could represent counts of `ACA`*→*`ATA` mutations for each sample). The signature definition matrix 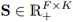 corresponds column-wise to a probability distribution over all features, for each signature, indexed by *k*′ ∈ [*K*], and as such ∑_*f*_ *s*_*fk*′_ = 1 *∀ k*′. Note that in this manuscript we use the index *k*′ for signatures and *k* for signature log-ratios. **S** was initially defined by non-negative matrix factorisation (NMF) with an initial set of samples, and it is *k*′ *j* redefined and maintained by COSMIC. We assume it is known. Let 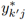 be the number of mutations attributed to signature *k*′ in observation *j*, i.e. **Y**^*∗*^ *∈* ℕ^*K×N*^ is the *exposure matrix*. Exposures are estimated by decomposing the count matrix **V** into the (given) signatures **S** and their exposures in each sample **Y**^*∗*^. **Y**^*∗*^ can be found from **V** and **S** via quadratic programming [18]. The exposure matrix **Y**^*∗*^ is our matrix of interest. In the methods section of this paper, for notational convenience, the transposed exposure matrix **Y** = (**Y**^*∗*^)^*T*^ is used instead of **Y**^*∗*^.

The signatures of PCAWG samples have been extracted using quadratic programming after extracting trinucleotide counts in a custom code, as well as using `mutSigExtractor` [19] on the vcf files – see Fig S1 for this comparison, as well as a comparison to values reported in [20] and extracted using `sigProfiler`. As patient ids are not available for download in the PCAWG data Synapse repository, these results are at the level of signature and cancer type.

### S2 Compositional data

In the case under consideration, for the *j*th clonality-based subsample, we can deduce *y*_*jK*_, the number of mutations corresponding to the last signature (also known as the exposure of signature *k* in subsample *j*), as 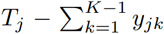, where *T*_*j*_ denotes the mutational toll of clonality-based subsample *j* (and to which exposures have been allocated). This makes such vectors (*K −* 1)-dimensional and compositional data, lying on a space called the simplex. The additive log-ratio transformation (ALR) and inverse additive log-ratio transformation (ALR^*−*1^) transformations correspond respectively to the logit and expit transformations often used in multinomial regressions. [42] address the common approaches and challenges related to the analysis of compositional data in more depth. Finally, we preferred the ALR transformation to the isometric log-ratio transformation (ILR) and centered log-ratio transformation (CLR) as CLR-transformed values cannot be used in regressions settings (as their covariance matrix is not full-rank), whilst the ALR and ILR yield the same results in this such setting, the ALR being a linear transformation of ILR. Moreover, if additional signatures are used in the model, the parameters in ALR space will not vary (provided the new signature is not used as baseline), whereas all parameters in ILR space would. In Section S2.2 we show the equivalence of results obtained when different signatures are used as ALR basis.

#### S2.1 Presence of zeros in signature exposures

The presence of zeros is a notorious complication for compositional models. In our model it is the coefficients ***β***_0_, ***β***_1_ (as well as the multivariate intercepts) which are in ALR space - note that we do not take the ALR transformation of the counts for each clonality-based subsample, where zeros do occur and where this would be problematic, possibly warranting replacement of zero exposures by a small non-zero value, which in turn could cause undesirable results in the model. Instead, the response of model is count data modelled by a multinomial distribution, where zero is included in the support.

It would be problematic, however, to include signatures of extremely low exposure, as it would lead to ***β*** coefficients being 0 or *→ ∞* (and this would apply to both ***β***_0_ and ***β***_1_). This is the reason why only active signatures are used. Similarly, if a signature is only present in the first group, ***β***_0_ can be estimated, but ***β***1 cannot, as *β*_1*j*_ *→ −∞*.

PCAWG exposures data do often contain zeros, as shown in Table S4: the fraction of zero exposures in a dataset varies from 2.7% to 38.6%, and, for a single signature, the fraction of samples with a zero exposure can be as high as 93%.

#### S2.2 Choice of baseline for ALR

The choice of baseline signature is not important for differential abundance results - both the softmax 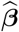 and the *p*-values are the same. We show these results by extending the simulation paradigm B2, with the following change: we modify the ***β***_1_ coefficients so that all ***β***_1_ are equal to a small number (0.1), which corresponds to only the last signature (SBS13) being differentially abundant. This allows us to have a scenario of subtle differential abundance, with a spread of *p*-values, but which also allows us to use as baseline either the signature that corresponds to the differentially-abundant signature, or another signature (here chosen as the fourth signature, SBS8, which is the signature of highest abundance in the whole dataset). The results are shown in Fig S26.

### S3 Assessment of the proposed models

#### S3.1 Bias and coverage of the estimator

The first set of simulations, referred to as **A1-A3**, are generated assuming

- a case involving *K* = 5 signatures and a number of mutations per sample of *T*_*j*_ = 180,
- a positive-definite matrix with no zero off-diagonal values in the random effect covariance matrix

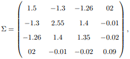
- varying overdispersion parameter values, *λ* = {2, 20, 100}, assumed equal per group (i.e., *λ*_*j*_ = *λ ∀ j*),
- ***β***_0_ = [5.5, 0.9, 0.35, 3] and ***β***_1_ = [*−*0.27, *−*1.28, 0.97, *−*1.28].

These sets of simulations, generated with lambda *λ* = {2, 20, 100} respectively. We focus here on the results corresponding to the models `diagREDM` and `fullREDM`. Both models allow the overdispersion parameter to be group-dependent as well as allowing for within-patient dependence, but differ in regards to the random effects correlation. In `diagREDM` and `fullREDM`, the random effects of the different signatures are respectively assumed to be independent and possibly dependent. Figs S2 and S3 respectively show the results for the `diagREDM` and `fullREDM` models. The upper plots show the bias (y-axes) for each of the *K −* 1 = 4 log-ratios of signatures (x-axes) for each element of ***β***_0_ (left) and ***β***_1_ (right) for three fixed values of *λ* (plots). The lower plots shows the coverage of 95% confidence intervals (y-axes) for the same configurations. The yellow bands correspond to the Monte Carlo tolerance area (defined by the quantiles 0.025 and 0.975 of a binomial with 1000 draws and a probability of success of 95%, the theoretical coverage). We can note that `diagREDM` shows biases for the ***β***_0_ elements, leading to very poor observed coverages. Estimates of the elements of ***β***_1_ are however reliably estimated with good coverages, suggesting that assuming an independence structure for the random effects is not problematic for the target parameter inference. Comparatively, `fullREDM` typically obtains less biases and better coverages for elements of ***β***_0_. For ***β***_1_, `fullREDM` and `diagREDM` lead to similar results for large values of *λ* and worse results for *λ* = 2, corresponding to a case with extreme overdispersion levels. However, in Fig S4, which shows the equivalent results for `singleREDM` we showcase the need for multivariate intercepts, as these biases and poor coverages are exhacerbated when using intercepts drawn from a single *σ*. In fact, as it follows from the softmax transformation, this model leads to the random effect only changing the abundance of the last signature, which is used as baseline. To summarise, our results suggest that, if assuming independence between the random effects lead to biases in the estimation of elements of ***β***_0_, this does not seem to translate to elements of ***β***_1_, our parameter of interest, provided multivariate random effects are used. Due to the strong reduction in the number of parameters to estimate, `diagREDM` thus appear as very attractive compared to `fullREDM`, whilst not being as restrictive as using a diagonal covariance matrix with shared elements along the diagonal.

Additionally, we performed equivalent simulations, considering `diagREDM` and `fullREDM`, for biologically- relevant parameters, taken from the PCAWG cohort by estimating the parameters using the `diagREDM`(in the case of uncorrelated data) and `fullREDM` models (in the case of correlated data). Three simulations are shown: (B1) for CNS-GBM without any signature correlations, (B2) for Lung-SCC without any signature correlations, and (B3) for Lung-SCC with positive and negative signature correlations. In all cases 1000 Monte Carlo simulations are performed for 200 samples.

The parameters are as follows. For simulation (**B1**) using parameters from CNS-GBM without any signature correlations, the parameters are

- *K* = 6 signatures and a number of mutations per sample of *T*_*j*_ = 3401 (median of the observed number of mutations),
- a positive-definite matrix of zero off-diagonal values in the random effect covariance matrix

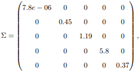
- overdispersion parameter value *λ* = 18, assumed equal per group (i.e., *λ*_*j*_ = *λ ∀ j*, corresponding to the average between the two estimated *λ*),
- ***β***_0_ = [*−*0.38, *−*1.13, *−*3.6, *−*6.96, *−*2.48] and ***β***_1_ = [*−*0.43, *−*0.17, 0.41, 1.03, *−*0.06].

For simulation (**B2**), using parameters from Lung-SCC without any signature correlations, the parameters are

- *K* = 6 signatures and a number of mutations per sample of *T*_*j*_ = 14072 (median of the observed number of mutations),
- a positive-definite matrix with no zero off-diagonal values in the random effect covariance matrix

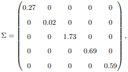
- overdispersion parameter value *λ* = 80, assumed equal per group (i.e., *λ*_*j*_ = *λ ∀ j*, corresponding to the average between the two estimated *λ*),
- ***β***_0_ = [*−*2.56, *−*0.39, 1.53, 1.77, 0.37] and ***β***_1_ = [1.07, 0.04, *−*0.74, *−*0.37, 07].

For simulation (**B3**) using parameters from Lung-SCC including a covariance matrix with positive and negative signature correlations, the same parameters as in (2) are used, except for ***λ*** and **Σ**:

- positive-definite matrix with no zero off-diagonal values in the random effect covariance matrix

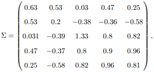
- an overdispersion parameter value *λ* = 87, assumed equal per group (i.e., *λ*_*j*_ = *λ ∀ j*), corresponding to the average between the two estimated *λ*).

Fig S5 shows the inference results using the un-correlated mixed-effects DM (`diagREDM`) and correlated mixed-effects DM (`fullREDM`) from simulation scenario (B1) of data simulated using previously-estimated parameters from the CNS-GBM cohort, with *N*_*s*_ = 200, no correlations, and a shared ***λ***. All remaining parameters are taken as estimated. Given that uncorrelated data are simulated, both models give the same results for bias and coverage. Both ***β***_0_ and ***β***_1_ are well recovered, although the penultimate element of ***β***_0_ has some biases. This corresponds to the ***β***_0_ in which the numerator of the log-ratio is the signature of lowest abundance.

In Fig S6 the equivalent simulation as above is carried out, but this time data are simulated from scenario (B2) using previously-estimated parameters from the Lung-SCC cohort, with *N*_*s*_ = 200, no correlations, and a shared ***λ*** (average between the two estimated *λ*). All remaining parameters are taken as estimated. Again, given that uncorrelated data are simulated, both models give the same results for bias and coverage. Bias and coverage results are satisfactory for both ***β***_0_ and ***β***_1_.

In Fig S7 data are simulated from the Lung-SCC cohort in simulation scenario (B3), with *N*_*s*_ = 200 and a shared ***λ*** (average between the two estimated *λ*). However, unlike (B1) and (B2), all remaining parameters are taken as estimated from the `fullREDM` model, and this includes correlations. The results for bias and coverage differ between the models owing to the presence of correlations. Bias and coverage results are satisfactory in both cases for ***β***_1_, but there is bias and low coverage for ***β***_0_ in the non-correlated `diagREDM` model (Figs S7a and S7c), as expected.

In conclusion, the results of these simulations – including simulations in which values are chosen to be representative of this type of data – indicate that ***β***_0_ might be more difficult to recover if the simpler model `diagREDM` is used, whereas ***β***_1_ is still well recovered. These results are consistent across simulations.

#### S3.2 Effect of the number of mutations on bias and coverage

The effect of the number of mutations on bias and coverage is assessed with biologically-informed simulation **B4**. Using parameters from Eso-AdenoCA without any signature correlations, the parameters are

- *K* = 10 signatures
- a varying number of mutations per sample: (1) equal to the observed number of mutations, and preserving the group (clonal/subclonal) information, ranging from 551 to 62315 (‘observed *T* ‘), (2) values half the size, ranging from 276 to 31158 (‘lower *T* ‘), (3) values ten times smaller, ranging from 55 to 6232 (‘lowest *T* ‘).
- a positive-definite matrix with no zero off-diagonal values in the random effect covariance matrix

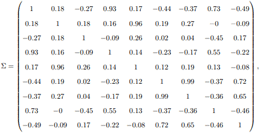
- two overdispersion parameters with values *λ*^(1)^ = 105.4 and *λ*^(2)^ = 50.4,
- ***β***_0_ = [*−*0.87, *−*2.2, 8 *−* 3.67, *−*0.10, *−*3.17, *−*0.76, *−*0.03, *−*0.84, *−*3.21]
- ***β***_1_ = [*−*0.27, 0.17, *−*0.19, *−*0.28, 0.05, *−*0.63, *−*0.7, 0 *−* 0.33, *−*0.21].

#### S3.3 Comparison with previous models

In this section we compare the output of our model to that of HiLDA and TCSM. We wanted to mention, additionally, [25], who they describe, from a theoretical perspective, a multivariate and unconstrained structure for the random effects as well as a covariate-dependent overdispersion parameter. Their estimator and implementation actually only consider a single random intercept parameter for all log-ratios of categories as well as a shared overdispersion parameter, thus imposing strong assumptions on the data dependence structure and variability, and the models we present here are extensions of this model.

##### S3.3.1 Data simulation

The simulation of data which can be used as input for TCSM and HiLDA is more complex than the simulation of data for `CompSign`, as both TCSM and HiLDA take as input substitution categories. In the case of TCSM these are trinucleotide substitutions (*F* = 96), and in the case of HiLDA it is a vcf file, which can be later converted to a *pmsignature* object, or a *pmsignature* directly. Moreover, in HiLDA signature extraction is performed using the pmsignature model by [43], which requires specifying the custom set of genomic features to use. Here we use, again, the single-base subtitution together with the two flanking bases, without any strand specificity, also leading to a maximum of *F* = 96 combinations of these features.

The substitution category matrix is, in turn, generated from PCAWG data (Lymph-CLL), by mixing mutations from clonal and subclonal mutations in certain proportions (this is a simulation framework very similar to the one used in [30]).

In particular, let 1 *− π, π ∈* [0, 1] be the mixing proportions. We simulate the same number of mutations as found in the Lymph-CLL PCAWG data. Given the exposures from clonal and sub-clonal mutations, we generate synthetic matrices **V** by drawing from a multinomial distribution and with probability vector the normalised signature exposures, for each patient-specific subsample, a number of mutations attributable to each signature (count exposures). For low values of *π*, these count exposures will be representative of clonal mutations (or mutations in the first group, more generally) in all patient-specific subsamples (no differential abundance), whereas for high values of *π* the count exposures will be representative of clonal mutations in patient-specific subsamples of one group and of subclonal mutations in patient-specific subsamples of the other (differential abundance). We then sample substitution categories with replacement from the 96-category vector, and with probabilities taken from the COSMIC signature definitions. This gives a vector of substitution categories for each patient-specific subsample. The mixing proportions *π* take the following values, in percentage: {0.034, 0.091, 0.250, 0.670, 2.3, 2.9, 3.7, 4.7, 6, 7.6, 9.5, 12, 27, 50, 62, 73}, and we have simulated 10 datasets with each set of parameters.

The count exposures from which we simulate mutations differ in simulated datasets C1-3: in (**C1**) they are a mixture of exposures from Lymph-CLL and Breast-AdenoCA (leading to a total of six signatures), i.e. a mixture of exposures where different signatures are active (only two are present in both datasets), in (**C2**) they are a mixture of exposures from the clonal and subclonal subsamples of Lymph-CLL, and (**C3**) is a variation of (C2) in which the patient pairing information of subsamples is preserved.

C1: data generated from a mixture of exposures from Lymph-CLL (four signatures: SBS1, SBS5, SBS9, SBS40) and Breast-AdenoCA (the most abundant four signatures: SBS3, SBS5, SBS40, SBS13)

C2: data generated from a mixture of clonal and subclonal exposures from Lymph-CLL

C3: data generated from a mixture of clonal and subclonal exposures from Lymph-CLL, preserving the information about the patient pairing

In the case of TCSM, which is run on the command line, this matrix and the covariate matrix are the only two input files. In the case of HiLDA, we synthetically create this *pmsignature* object from the substitution category matrix. The number of signatures which both HiLDA and TCSM need as input is given as the number of signatures used in the simulation. In the case of CompSign, signatures need to be extracted first -we do this using quadratic programming, exactly in the same way as it is done for the PCAWG samples in the Results section - also specifying the signatures from which we have simulated data. The extracted signature matrix is used as input for the `CompSign diagREDM` model.

##### S3.3.2 Comparison of runtime

We were unable to run several models with HiLDA, as its runtime exceeded the 10-hour and even 24- hour mark. These appear as missing in Table S3 and any figures relating to simulated datasets C1 or C3. For HiLDA, the function `hildaTest` was used and timed within R. For TCSM a two-step process is needed: the `run tcsm.R` was run, followed by `estimate significance.py`; both were timed.

##### S3.3.3 Comparison of *β*_1_ to equivalent parameters in HiLDA and TCSM

To assess the agreement in the estimated parameters of differential abundance, we have had to transform the output of each of the models. For `diagREDM` this has been done by taking the softmax-transformed 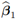, for HiLDA by computing the per-signature sum of squares of 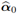 and 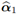, and for TCSM by plotting the estimated coefficients in the “effect” matrix. After extracting these parameters, and as we do not have an indication of the order in which signatures appear in each of the datasets, these estimates are sorted based on the *β*_0_-equivalent estimates of general signature abundance (see Section S3.3.4).

##### S3.3.4 Comparison of *β*_0_ to equivalent parameters in HiLDA and TCSM

To assess if the estimated abundance of signatures is in agreement, we have plotted the abundance of signatures in probability space. For `diagREDM` this has been done by taking the softmax-transformed 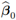, for HiLDA by normalising 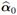 so that 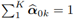, and for TCSM by plotting the estimated parameters in the “effect” matrix. After extracting these parameters, and as we do not have an indication of the order in which signatures appear in each of the datasets, these estimates are sorted – as HiLDA and TCSM extract signatures *de novo*, it might be the case that all models are working with completely different signatures.

##### S3.3.5 Assessment of signature recovery from TCSM

The cosine similarities from Fig S15 are computed as follows: for each estimated signature in the definition matrix 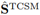 we compute its cosine similarity to all COSMIC signatures **S**^COSMIC^. We then pair each estimated signature to the COSMIC signature that represents it best, and ensuring that no two estimated signatures are paired to the same COSMIC signature, by sequentially pairing the estimated signature of highest cosine similarity. This gives, for each run of TCSM (equivalently, each simulated dataset) a set of cosine similarities of length *d*. High cosine similarities are indicative of a good recovery of signature definitions.

#### S3.4 Effect of active signature selection on differential abundance

We have extended the simulation strategy based on the simulation of mixtures so that we can compare the results of our models when we extract signatures using four types of subsets of active signatures. These datasets, (**C4**), have been created in the same way as (C2), but using exposures from the four most abundant signatures in Prost-Adenoca. We consider four strategies for active-signature selection: the “truth” subset of signatures used in the simulation (*Simulated signatures*), and three sets of active signatures determined by first extracting signatures de novo using all COSMIC signatures, and then refining them by selecting the top signatures that contribute to a total of 80% of mutations in a sample (*Signature sum > 80% samples*), or to a 60% of the samples (*Signature sum > 60% samples*), or signatures which are present (non-zero) in at last 80% of samples (*Signature active > 80% samples*). The signatures which are considered to be active in strategy 3 are a subset of those active in strategy 2.

